# Tau-induced mitochondrial reverse electron transport drives neurodegeneration

**DOI:** 10.64898/2026.04.04.716514

**Authors:** Wen Li, Suman Rimal, Sunil Bhurtel, Lucas Yeung, Benjamin G. Lu, Lea T. Grinberg, Salvatore Spina, Maria Inmaculada Cobos Sillero, William W. Seeley, Su Guo, Bingwei Lu

**Author notes:** These authors contributed equally (Wen Li, Suman Rimal).

## Abstract

Hyperphosphorylation and aggregation of the microtubule-associated protein tau are recognized as pathological hallmarks of tauopathies; however, the biological activity of tau that drives its pathophysiological effects remains poorly understood^1–6^. Mitochondrial dysfunction is a common feature of tauopathies^7,8^. Despite this, the mechanistic link between tau abnormalities and mitochondrial dysfunction, as well as its relationship to tau’s physiological function, remains unclear. Here, we demonstrate that tau regulates mitochondrial reverse electron transport (RET), which produces excess ROS, reduces the NAD^+^/NADH ratio, and is activated by aging or stress. In flies, mice, and human induced pluripotent stem cells (hiPSC)-derived neurons, tau depletion eliminates stress-induced RET and confers significant stress resistance. Mechanistically, tau enters mitochondria and directly interacts with the mitochondrial complex I (C-I) subunit NDUFS3, enhancing RET activation in a phosphorylation-dependent manner that correlates with tau pathogenicity. Elevated RET further drives tau hyperphosphorylation, establishing a self-perpetuating pathological loop. Blocking tau entry into mitochondria or disrupting tau/NDUFS3 interaction reduces tau-induced RET. Genetic or pharmacological inhibition of RET protects against tau-induced neurodegeneration across species. RET regulation represents a previously unrecognized normal function of tau that becomes pathological in disease, providing a therapeutic target for conditions characterized by tau abnormalities and mitochondrial dysfunction.

## Main

Many neurodegenerative diseases share overlapping neuropathological and clinical features. If this overlap reflects a shared pathogenic mechanism, targeting it therapeutically could potentially address multiple diseases. Hyperphosphorylation and aggregation of tau are pathological hallmarks of primary tauopathies, such as frontotemporal dementia (FTD) and progressive supranuclear palsy (PSP), as well as secondary tauopathies, which include common neurodegenerative diseases like Alzheimer’s disease (AD) and possibly Huntington’s disease and Parkinson’s disease^1–6^. Tau abnormalities are also observed in experimental models of autism^9^, epilepsy^10^, stroke^11^, and traumatic brain injury^12^, where tau reduction has shown beneficial effects – even in the absence of neurofibrillary tangles (NFTs), the prominent intraneuronal tau aggregates characteristic of AD and other tauopathies^1–3^. This suggests that non-aggregated soluble tau species may mediate the pathological effects of tau. Tau is normally a soluble protein enriched in neuronal axons but is also found in neuronal dendrites, cell bodies^13^, and non-neuronal cells^14,15^. Tau is phosphorylated at many sites under normal conditions but becomes hyperphosphorylated in disease, suggesting that phosphorylation regulates its pathophysiological activities^15,16^. Genetic mutations in tau causing FTD with Parkinsonism linked to chromosome 17^17–20^, the relationship between tau pathology and brain dysfunction in AD^21^, and the beneficial effects of tau reduction observed in experimental AD models^22,23^ collectively provide strong evidence that tau abnormalities are pathogenic.

Tau was originally identified as a protein essential for microtubule (MT) assembly^24^, and disease-associated mutations or post-translational modifications (PTMs) were thought to impair this function^25^, leading to the hypothesis that pathogenic tau causes neuronal damage by disrupting MT structure and function^26^. However, tau’s rapid kiss-and-hop interaction with MT^27^ suggests that it is readily available for interactions beyond MT. As an intrinsically disordered protein, tau engages in diverse protein-protein interactions^28,29^. Notably, the experimental reduction of tau *in vivo* did not disrupt MT-dependent neural processes^22,30^, and evidence indicates that tau-induced cellular damage may extend beyond its role in MT function^31–34^, pointing to alternative pathogenic mechanisms of tau.

Mitochondrial dysfunction is a prominent feature of tauopathies and other conditions associated with tau abnormalities^7,8^. Tau interacts with mitochondrial proteins, and recent studies in hiPSC-derived neurons suggest that the mitochondrial interactome of tau is extensive^34^. Overexpression studies in cell culture and animal models have linked pathogenic tau to disruptions in various mitochondrial processes^35–39^. However, an unresolved question remains: do these effects reflect tau’s normal physiological function^15^? Addressing this question is essential for effectively targeting tau in therapeutic development, as a deeper understanding of its normal function in relation to its pathological effects is critical.

During oxidative phosphorylation (OxPhos), electrons enter the electron transport chain (ETC) from C-I in the form of NADH. These electrons are transferred through carriers such as co-enzyme Q10 (CoQ) and cytochrome C, eventually reaching complex IV, where they react with O_2_ to form H_2_O. This forward electron transport (FET) process generates a proton gradient needed for ATP synthesis, converts NADH to NAD^+^, and produces small amounts of reactive oxygen species (ROS) as a byproduct^40^. In contrast, reverse electron transport (RET) generates large amounts of ROS and converts NAD^+^ back to NADH, thereby reducing the mitochondrial and cellular NAD^+^/NADH ratio^41–43^. RET occurs under conditions of high CoQH_2_ levels or elevated mitochondrial membrane potential (Δp), which provide the energy required to drive electrons backward against the redox potential gradient of the ETC, moving them in reverse from CoQH^44^. RET has been implicated in various disease states^42,43,45,46^, but its normal physiological role and regulatory mechanisms in health and disease remain poorly understood. Using *in vivo* models in *Drosophila* and mice, as well as hiPSC-derived neuronal systems, we have uncovered a conserved role of tau as a regulator of RET. This discovery provides new insights into tau’s physiological function and its relevance to tauopathies.

### RET activation in tauopathy models and patient brain tissues

To investigate the mitochondrial processes relevant to tauopathy, we examined RET in multiple models, including *Drosophila*, mice, hiPSC-derived neuronal systems, and patient brain tissues. RET activation was consistently observed across all models of tauopathies. We utilized hiPSCs carrying the tau-P301L mutation, a well-characterized pathogenic variant that induces tau hyperphosphorylation and mitochondrial dysfunction^35,47^. Isogenic wild-type iPSCs served as controls, enabling investigation in disease-relevant cells with physiological tau levels and patient-specific genetic backgrounds, while avoiding confounding factors inherent to transgenic overexpression models or iPSCs with unmatched genetic profiles. To induce neurons affected in tauopathy, NEUROGENIN 2 (NGN2) was used to drive neuronal differentiation^48^. Tau-P301L hiPSC-derived neurons exhibited increased ROS (Fig. 1a-c) and decreased NAD^+^/NADH ratio (Fig. 1a,b), key markers of RET activation^42^. To directly assess RET alterations, we used purified mitochondria supplemented with substrates driving RET (succinate) or FET (malate/glutamate)^43, 49–53^. Measurements revealed significantly elevated RET activity in tau-P301L hiPSC-derived neurons, while FET remained relatively unchanged (**Fig. 1a,b; Extended Data Fig. 1a,b**). Further validation came from experiments using RET inhibitors CPT-2008 (CPT)^42^ and low-concentration rotenone^54^, both of which abolished the ROS and NAD^+^/NADH alterations in tau-P301L mitochondria (Extended Data **Fig. 1c,d**), confirming RET as the source of the observed changes.

**Fig. 1.**
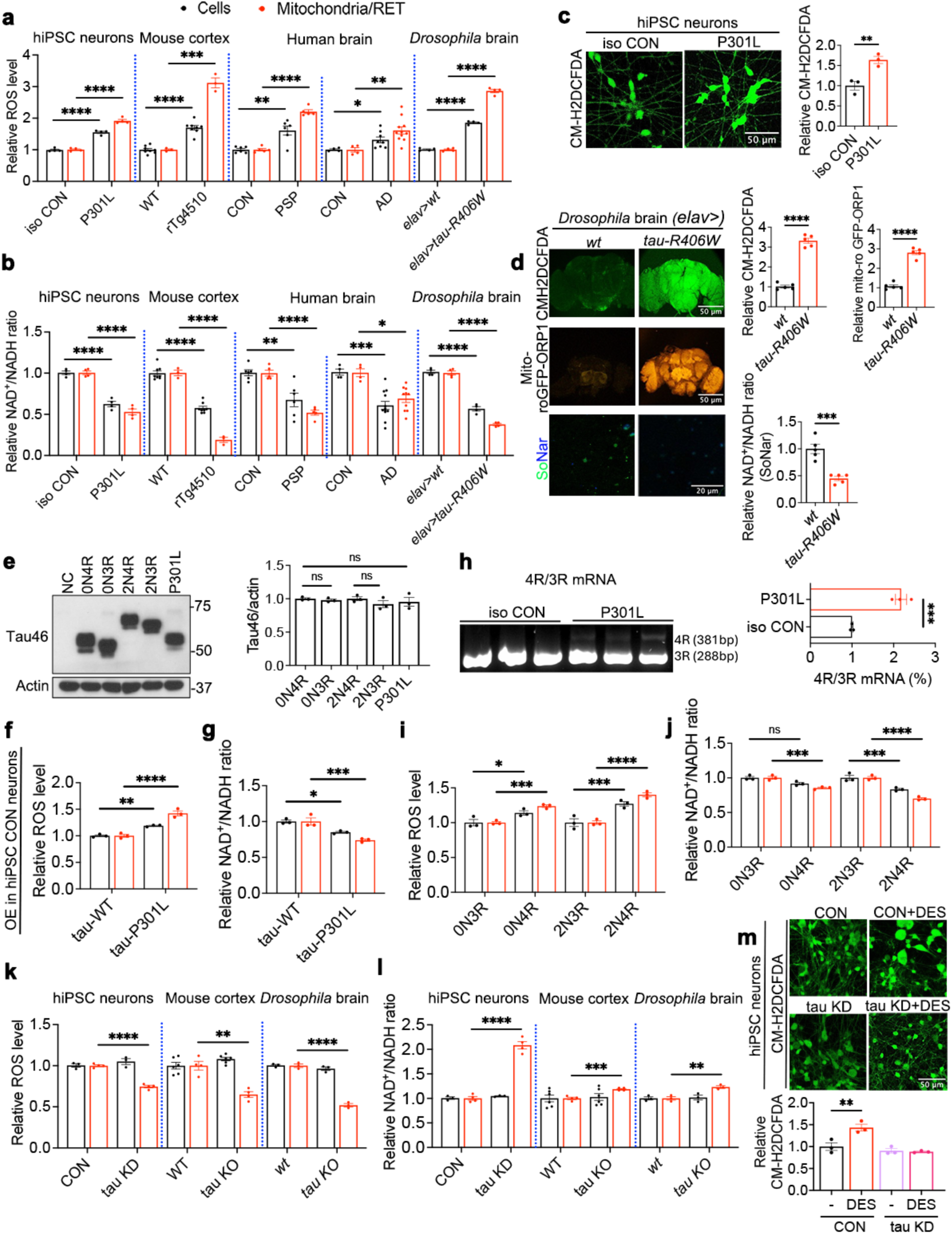
A physiological role of tau in RET that is deregulated in tauopathies. **a, b**, Measurements of ROS (**a**) or NAD^+^/NADH ratio (**b**) from cells (black bars) derived from cell culture or tissue homogenates of tauopathy human iPSC neurons, rTg4510 mice, human PSP and AD postmortem brain tissues, or transgenic *Drosophila* brains expressing human tau-R406W or from mitochondria (red bars) purified from these cells and respiring under the RET condition. Normalized values relative to their respective controls are shown. **c**, Representative images and quantification of CM-H2DCFDA staining of ROS in tau-P301L and isogenic WT control hiPSC-derived neurons. **d**, Representative images and quantification of CM-H2DCFDA, mito-roGFP-ORP1, and SoNar signals in WT control and *elav>tau-R406W Drosophila* brains. **e**, Representative western blot (WB) and quantification of tau isoform expression in healthy hiPSC-derived transfected with lenti-viruses expressing various tau isoforms. P301L is of the 0N4R isoform. **f**, **g**, Measurements of ROS (**f**) or NAD^+^/NADH ratio (**g**) from hiPSC-derived neurons overexpressing tau-WT or tau-P301L (black bars) or mitochondria purified from these cells and respiring under RET condition (red bars). **h**, RT-PCR analysis and quantification of mRNAs for 4R and 3R tau in control and tau-P301L hiPSC-derived neurons cultured for 3 weeks. **i**, **j**, Measurements of ROS (**i**) or NAD^+^/NADH ratio (**j**) from hiPSC-derived neurons overexpressing various tau isoforms (black bars) or mitochondria purified from these cells and respiring under the RET condition (red bars). **k**, **l**, ROS (**k**) or NAD^+^/NADH ratio (**l**) measurements in cells derived from cell culture or tissue homogenates of control vs. tau KD human iPSC neurons or control vs. tau KO mice or *Drosophila* (black bars) or mitochondria purified from these cells and respiring under the RET condition (red bars). **m**, Representative images and quantification of CM-H2DCFDA staining in control and tau KD hiPSC-derived neurons with or without DES (20 mM) treatment. All data are means ± SEM; statistical significance was determined by two-tailed unpaired Student’s *t* test. *p<0.05; **p<0.01; ***p<0.001; ****p<0.0001. For cell culture and fly model studies, n=3-5 biological repeats; for mouse and patient tissue studies, each data point represents an individual mouse or patient sample.

To test if RET activation occurs *in vivo*, we used the rTg4510 tauopathy mice expressing a regulatable tau-P301L transgene^55^. At 8 months of age, when tau-related neuropathology and behavioral deficits are prominent, brain tissues from rTg4510 mice showed increased ROS and decreased NAD^+^/NADH ratio, consistent with RET activation (**Fig. 1a,b**). RET assays on purified brain mitochondria confirmed CPT- (see later) and rotenone-inhibitable RET activity, while FET remained unaffected (Extended Data **Fig. 1a-d**).

To validate these findings, additional methods were employed to measure NAD^+^/NADH ratio, including enzymatic cycling reactions with alternative extraction protocols (Extended Data **Fig. 1e**) and mass spectrometry-based metabolomics to quantify NAD^+^ and NADH (Extended Data **Fig. 1f**). These results confirm RET activation in tauopathy models. Furthermore, the mitochondrial preparations displayed active respiration under both FET and RET conditions, fueled by different substrates, indicating their quality and structural integrity. This was corroborated by mitochondrial respirometry using a Seahorse XF analyzer (Extended Data **Fig. 1g**).

To strengthen findings across species, we examined transgenic flies expressing human tau with the R406W mutation. Brain tissues from tau-R406W flies demonstrated increased ROS and decreased NAD^+^/NADH ratios, indicative of RET activation. Mitochondrial assays confirmed elevated RET activity, while FET remained relatively unchanged (Fig. 1a,b; Extended Data Fig. 1a,b). ROS and NAD^+^/NADH changes were validated using distinct techniques, including the CM-H2DCFDA dye, a genetically encoded mitochondrial ROS reporter (mito-roGFP-ORP1), and a NAD^+^/NADH reporter (SoNar) ^43,56^ (Fig. 1d).

To investigate the clinical relevance of RET alterations, we examined frozen brain tissues from patients with progressive supranuclear palsy (PSP) and AD. Both PSP and AD brain tissues showed increased ROS and decreased NAD^+^/NADH ratios, alongside elevated RET activity in purified mitochondria respiring under RET conditions (Fig. 1a,b). In contrast, FET activity remained unaffected (Extended Data Fig.1a,b). These findings indicate that RET elevation observed in tauopathy models is relevant to human disease conditions.

The use of isogenic wild-type (WT) and tau-P301L hiPSC neurons expressing tau at physiological levels allowed us to draw a clear conclusion that tau-P301L exhibits a stronger ability to promote RET activity compared to tau-WT. When tau-WT and tau-P301L of the human 0N4R isoform were overexpressed at comparable levels in control hiPSC neurons (Fig. 1e), tau-P301L consistently induced higher ROS production and a greater reduction in the NAD^+^/NADH ratio under RET conditions, while FET remained unaffected (Fig. 1f,g; Extended Data Fig. 1h,i). These results demonstrate that the P301L mutation enhances tau’s RET-promoting activity in hiPSC-derived neurons.

In the adult human CNS, tau exists as six alternatively spliced isoforms that differ in the number of N-terminal inserts (0N, 1N, or 2N) and C-terminal repeats (3R or 4R). As previously reported, control hiPSC neurons predominantly expressed tau in the 3R isoform^57^. However, tau-P301L iPSC neurons expressed both 3R and noticeable levels of 4R isoforms (Fig. 1h), suggesting that tau-P301L may alter its own splicing, consistent with prior findings^58^. Since the P301L mutation is exclusively present in 4R tau, these results indicate that 4R tau plays a greater role in driving enhanced RET in tau-P301L neurons. Supporting this, control hiPSC neurons overexpressing different tau isoforms showed that 4R tau induced higher RET compared to 3R tau (Fig. 1i,j), while FET remained unaffected (Extended Data Fig. 2g,h)

To further investigate the effect of pathogenic tau mutations on RET *in vivo*, we used transgenic flies expressing similar levels of human tau-WT, tau-P301L, or tau-R406W (Extended Data Fig. 2a,b). Both tau-P301L (Extended Data Fig. 2c,d) and tau-R406W (Extended Data Fig. 2e,f) significantly increased RET activity compared to tau-WT, as evidenced by elevated ROS production and a reduced NAD^+^/NADH ratio. This provides additional support that these mutations amplify tau’s ability to drive RET dysfunction.

### RET activity is attenuated by tau depletion in *Drosophila*, mice, and hiPSC-derived neurons

The hyperactivation of RET observed in tauopathy may reflect either a toxic gain-of-function by pathogenic tau, unrelated to its normal role, or a physiological function of tau that becomes deregulated in tauopathy. To distinguish between these possibilities, we examined tau knockout (KO) and knockdown (KD) conditions. In hiPSC-derived neurons with tau KD by lentiviral delivery of shRNA, tau protein levels were significantly reduced (Extended Data Fig. 2i), and these neurons displayed normal ROS and NAD^+^/NADH ratio compared to control neurons (Fig. 1k,l). In contrast, mitochondrial RET activity was significantly impaired in tau KD neurons (Fig. 1k,l), while FET activity remained unaffected (Extended Data Fig. 2j,k). This effect was further confirmed by treating neurons with diethyl-succinate (DES), a substrate known to promote RET in mammalian cells^59^ and flies^43^. Control neurons treated with DES exhibited increased ROS production, whereas tau KD neurons failed to generate ROS under the same conditions (Fig. 1m).

Similarly, brain tissues from a well-characterized tau KO mouse model^60^ showed normal ROS levels and the NAD^+^/NADH ratio (Fig. 1k,l), However, when purified brain mitochondria were analyzed, RET activity was reduced while FET activity remained unchanged (Fig. 1k,l; Extended Data Fig. 2j,k). Comparable findings were observed in tau KO flies^61^ (Fig. 1k,l; Extended Data Fig. 2j,k). These results indicate that tau-deficient mitochondria have compromised RET activity, which becomes evident when mitochondria are fueled by RET-specific substrates *in vitro*. Under normal conditions, mitochondria predominantly rely on FET, with RET contributing minimally to cellular ROS production and NAD^+^/NADH ratio. This explains the relatively unchanged ROS levels and NAD^+^/NADH ratio in tau KO brain tissues or tau KD hiPSC-derived neurons. Taken together, these findings demonstrate a conserved physiological role for tau in regulating RET activity across species.

### Tau actively promotes RET in a phosphorylation-dependent manner

The mechanism by which tau regulates RET could be either direct or indirect. To explore this, we treated mitochondria isolated from 6-month-old WT mouse (C57/BL6) muscle with soluble recombinant human tau (2N4R form), pretreated with glycogen synthase kinase 3β (GSK-3β) under conditions conducive to protein import. This recombinant tau was recognized by the PHF-1 p-tau antibody (p-S396/S404), but not by antibodies specific for other phosphorylation sites, including p-S262, AT-100 (p-T212/S214), or CP13 (p-S202) (Extended Data Fig. 3a). These results are consistent with GSK-3β being a kinase generating the PHF-1 epitope^62^. Notably, while a non-phosphorylatable tau mutant (tau-S404A) failed to induce RET, an equivalent amount of GSK-3β treated tau-WT robustly promoted RET-ROS production (Fig. 2a). Since phosphorylation at S404 primes tau for phosphorylation at S396^63^, tau-S404A is likely deficient in S936 phosphorylation. These findings suggest that tau promotes RET in a PHF-1 site-dependent manner.

**Fig. 2.**
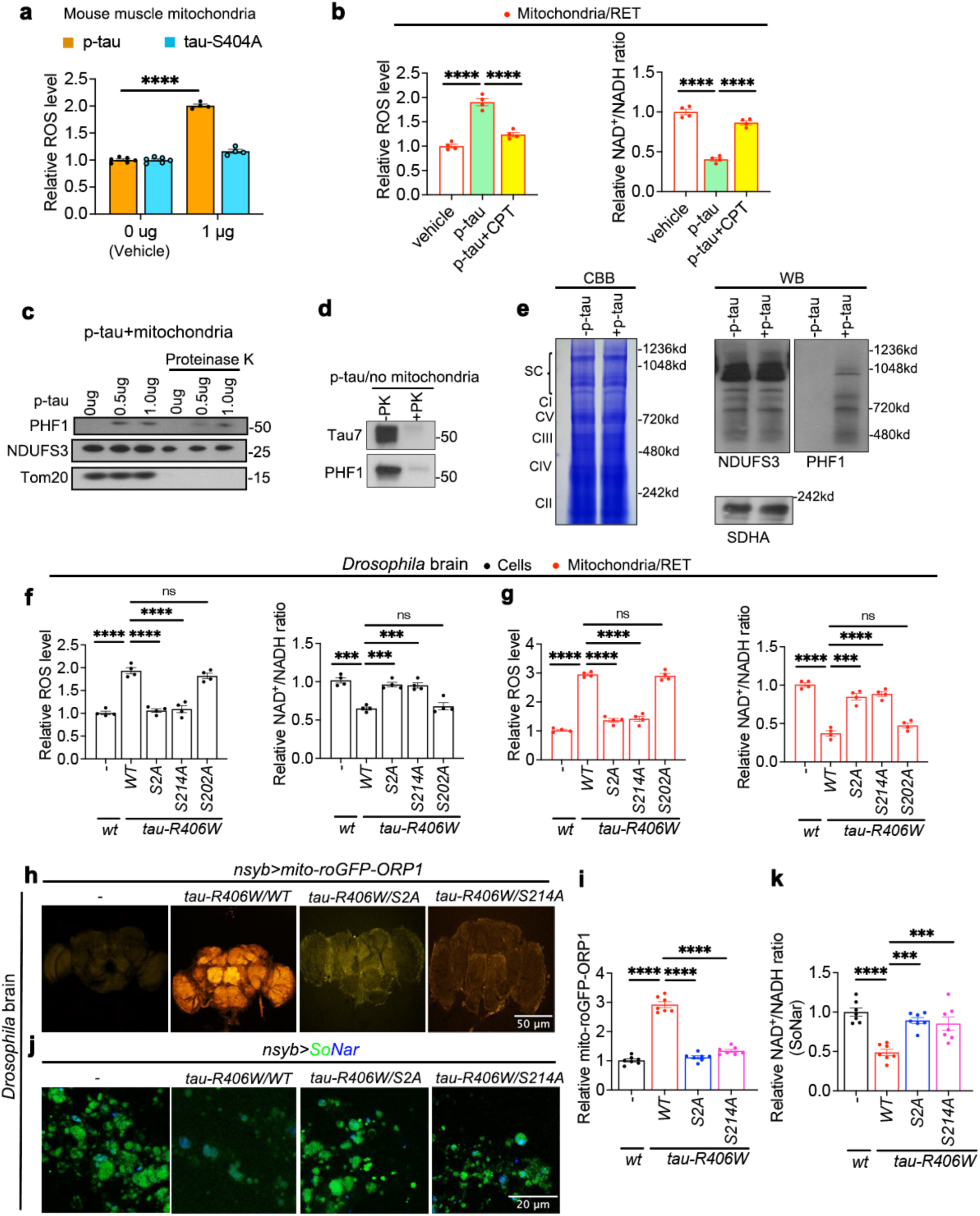
Phosphorylation-dependent RET induction by tau. **a**, Measurement of mitochondrial RET-ROS induced by phosphorylated or phospho-mutant (S404A) recombinant human tau *in vitro*. **b**, Measurement of ROS and NAD^+^/NADH in mitochondria treated with p-tau and respiring under the RET condition. Assays were performed with or without the RET inhibitor CPT. **c**, WB of mitochondria treated with PK showing the presence of p-tau inside mitochondria. Recombinant human 2N4R tau pretreated with GSK-3β and muscle mitochondria purified from 6-month-old C57BL6 mice were used in **a**-**c**. **d**, PK treatment of recombinant p-tau in the absence of mitochondria. **e**, Coomassie brilliant blue (CBB) staining (left) and WB analysis (right) of BN-PAGE separated mitochondrial proteins with or without prior incubation with p-tau. SC: supercomplex. **f**, **g**, Relative ROS and NAD^+^/NADH ratio of cells from *Drosophila* brain tissues expressing WT or phospho-mutant forms of human tau-R406W (**f**) or mitochondria purified from these cells and respiring under RET condition (**g**). **h-k**, Live imaging and quantification of genetically encoded mito-roGFP-ORP1 (**h**, **i**) or SoNar (**j**, **k**) reporters assessing mitochondrial ROS or NAD^+^/NADH levels in *Drosophila* brain tissues expressing WT or phospho-mutant forms of tau-R406W driven by neuronal *nsyb-Gal4*. All data are means ± SEM; statistical significance was determined by one-way ANOVA with Tukey’s post hoc test (**b, f, g, i, k**), or two-tailed unpaired Student’s *t* test (**a**). *p<0.05; **p<0.01; ***p<0.001; ****p<0.0001. Each data point represents an independent experimental repeat.

The effect of phosphorylated tau (p-tau) on ROS production and NAD^+^/NADH ratio was dose-dependent (Extended Data Fig. 3b) and was inhibited by the RET inhibitor CPT (Fig. 2b), supporting the notion that p-tau acts via RET. Additionally, p-tau was efficiently imported into mitochondria, as demonstrated by its protection from Proteinase K (PK) digestion - a pattern resembling that of the inner mitochondrial membrane protein NDUFS3 - while the outer membrane protein Tom20 was completely digested (Fig. 2c). In contrast, p-tau was sensitive to PK digestion when not mixed with mitochondria (Fig. 2d), confirming that its resistance to PK digestion was due to mitochondrial import rather than intrinsic properties of the protein. Furthermore, after mitochondrial import, PHF-1 tau was associated with the supercomplex, C-I holoenzyme, and C-I subcomplexes, as shown by Blue Native Polyacrylamide Gel Electrophoresis (BN-PAGE) (Fig. 2e). These observations are consistent with previous reports of p-tau localization to mitochondria^64^ and tau interaction with mitochondrial proteins^34^, although the intramitochondrial function of tau has remained unclear. Our data provide the first evidence that soluble tau species phosphorylated at the PHF-1 epitope can access C-I and actively promote RET.

To test the *in vivo* relevance of this phospho-dependent regulation of RET by tau, we utilized transgenic flies expressing mutant human tau transgenes. Specifically, we generated flies expressing tau-R406W-S2A, in which two PAR-1/MARK kinase target sites (S262 and S356) were rendered unphosphorylatable^65^. PAR-1/MARK kinase is known to orchestrate temporally ordered tau phosphorylation, with S262/S356 phosphorylation serving as a prerequisite for downstream tau kinases to generate PHF-1 and AT100 epitopes^65^. In flies expressing tau-R406W-S2A, RET promotion was completely abolished compared to flies expressing tau-R406W at equivalent levels (Fig. 2f,g; Extended Data Fig. 3d,e). Similarly, flies expressing tau-R406W-S214A, in which the AT100 site was mutated, also failed to promote RET (Fig. 2f,g; Extended Data Fig. 3d,e). *In vivo* mitochondrial ROS (Fig. 2h,i) and NAD^+^/NADH ratio (Fig. 2j,k) assays using genetically encoded reporters confirmed these findings. In contrast, flies expressing tau-R406W-S202A, in which the CP13 site was mutated, retained the ability to promote RET (Fig. 2f,g; Extended Data Fig. 3d,e). Importantly, tau-R406W-S214A and tau-R406W-S202A were expressed at similar levels as tau-R406W (Extended Data Fig. 3c,f). Interestingly, tau-R406W-S202A exhibited similar neurotoxicity to tau-R406W^65^, whereas tau-R406W-S2A^65^ and tau-R406W-S214A were largely non-toxic (Extended Data Fig. 3g). Furthermore, PHF-1 phosphorylation was greatly reduced in tau-R406W-S2A and tau-R406W-S214A, but less affected in tau-R406W-S202A (Extended Data Fig. 3c). Together, these results suggest that tau’s ability to promote RET correlates with its neurotoxicity, and that different phosphorylation sites contribute variably to RET regulation and tau toxicity - likely via their impact on PHF-1 p-tau formation.

### Aging and hyperthermic stress, which promote RET, also promote tau phosphorylation

If p-tau plays a key role in RET regulation, tau phosphorylation status may be altered under conditions that activate RET. RET is known to be activated in aged flies^43^, providing an opportunity to test this hypothesis. Although most available p-tau antibodies generated against human tau do not recognize fly tau, an antibody targeting the conserved p-S262 site detected an age-related increase in fly tau phosphorylation, when normalized to total tau or actin levels (Extended Data Fig. 4a). Similarly, in aged mouse brains, we observed increased endogenous p-tau positive for p-S262, PHF-1, and AT100 epitopes. These increases were evident when p-tau levels were quantified relative to actin or normalized by the p-tau/total tau ratios (Extended Data Fig. 4b).

Next, we examined the impact of stress on tau phosphorylation. Mice subjected to hyperthermic stress showed a significant increase in p-S262, PHF-1, and CP13 tau phosphorylation (Extended Data Fig. 4c). This increase in tau phosphorylation was accompanied by enhanced RET activity, while FET remained unchanged in heat-stressed mouse brains (Extended Data Fig. 4d,e). Similarly, flies exposed to heat stress displayed elevated RET activity without changes in FET (Extended Data Fig. 4f,g), along with increased phosphorylation of tau at the p-S262 site (Extended Data Fig. 4h). These findings demonstrate that stress conditions, such as hyperthermia, promote tau phosphorylation and enhance RET activity, suggesting a link between stress-induced tau phosphorylation and RET activation.

### Tau compromises survival and behavioral performance under stress via RET-related mechanisms

To investigate the physiological role of tau, we first analyzed flies with tau deletion. Tau deletion conferred resistance to heat stress, significantly extending lifespan when flies were raised at 32°C (Extended Data Fig. 5a). Consistent with a previous study^61^, tau deletion flies raised at room temperature (25°C) did not exhibit any lifespan changes (Extended Data Fig. 5b). To identify tissue- and cell type-specific roles of tau in the stress response, we used RNAi to knock down tau expression in specific tissues, with RNAi efficiency confirmed by western blotting (Extended Data Fig. 5c). Tau RNAi in neuronal or muscle tissues increased survival under heat stress (Extended Data Fig. 5d,e), whereas knockdown in other tissues had no effect (Extended Data Fig. 5f). These findings highlight the importance of neuromuscular tau in the stress response. The significant effect of muscle-specific tau RNAi aligns with tau expression in mouse and fly muscle tissues (Extended Data Fig. 5g,i), where it promotes RET (Extended Data Fig. 5h,j,k).

We next explored whether the stress resistance phenotype associated with tau deletion was related to RET. Inhibition of RET with CPT mimicked the survival benefits of tau RNAi under heat stress (Extended Data Fig. 6a). Furthermore, CPT treatment of tau KO flies did not provide additional lifespan benefits (Extended Data Fig. 6b), suggesting that tau KO and CPT act through a shared mechanism - RET inhibition - to enhance stress resistance and extend lifespan. Conversely, treatment with the RET inducer DES partially blocked the stress resistance conferred by tau KO (Extended Data Fig. 6c), further supporting the connection between tau’s role in RET regulation and stress response.

To assess the effect of tau deletion on stress resistance in mice, we subjected WT and tau KO mice to heat stress. Heat stress was applied for two weeks by placing the mice in an incubator set to 43°C with 60 ± 10% humidity for 15 minutes each day. After the heat stress period, cognitive performance was assessed using the novel object recognition (NOR) assay, and anxiety-related behavior was evaluated using the elevated plus maze (EPM) assay. WT mice exposed to heat stress performed poorly compared to those maintained at room temperature (25°C), as indicated by a lower discrimination index in NOR (Extended Data Fig. 6d) and reduced time spent in the open arm in the EPM (Extended Data Fig. 6e). Notably, these behavioral defects were ameliorated by CPT treatment (Extended Data Fig. 6d,e), supporting a RET-dependent origin of the stress-related impairments. Similarly, tau KO mice were behaviorally resilient to heat stress, and CPT did not further improve their behavior (Extended Data Fig. 6d,e), reinforcing the conclusion that tau KO and CPT provide stress resistance through similar RET-related mechanisms. Consistent with these observations, *in vitro* mitochondrial RET assays revealed that both CPT treatment and tau KO similarly inhibited heat stress-induced RET activation (Extended Data Fig. 6f,g). Comparable results were observed in tau RNAi (Extended Data Fig. 6h,i) or tau KO flies (Extended Data Fig. 6j), further confirming that tau deletion enhances stress resistance by inhibiting RET. Together, these findings reveal a detrimental effect of tau under chronic stress conditions, primarily through its ability to promote RET activity.

### Tau directly interacts with NDUFS3 and alters its RET-related interactions within C-I

To understand the molecular mechanisms by which tau promotes RET, we made several predictions: 1) tau/p-tau should be present inside mitochondria; 2) tau/p-tau may directly interact with C-I proteins; and 3) tau/p-tau may induce conformational changes in C-I that influence RET. We tested these predictions, focusing on PHF-1 tau, which is important for RET.

First, we analyzed mitochondrial fractions and subfractions from rTg4510 mouse brains. We observed p-tau inside mitochondria, with a distribution pattern encompassing the inner mitochondrial membrane (IMM) where the C-I marker NDUFS3 is localized (Fig. 3a,b). p-tau was also present in mitochondria purified from normal mouse brains, with levels increasing with age (Extended Data Fig. 7a). PK treatment of purified mitochondria reduced p-tau levels, but a substantial fraction remained protected (Fig. 3c). This protection was not due to intrinsic resistance of p-tau to PK digestion, as p-tau was efficiently digested in the absence of mitochondria (Fig. 2d). When Triton X-100 was added, both PHF-1 p-tau and NDUFS3 were completely digested by PK (Fig. 3c), confirming that p-tau is present inside mitochondria. These findings align with previous reports of tau’s presence in mitochondria^64^. Proteomic databases, such as MitoCoP, have also identified tau in purified mitochondria from cultured cells. However, tau did not appear in the high-confidence list due to its non-exclusive mitochondrial localization and cell line-specific nature^66^.

**Fig. 3.**
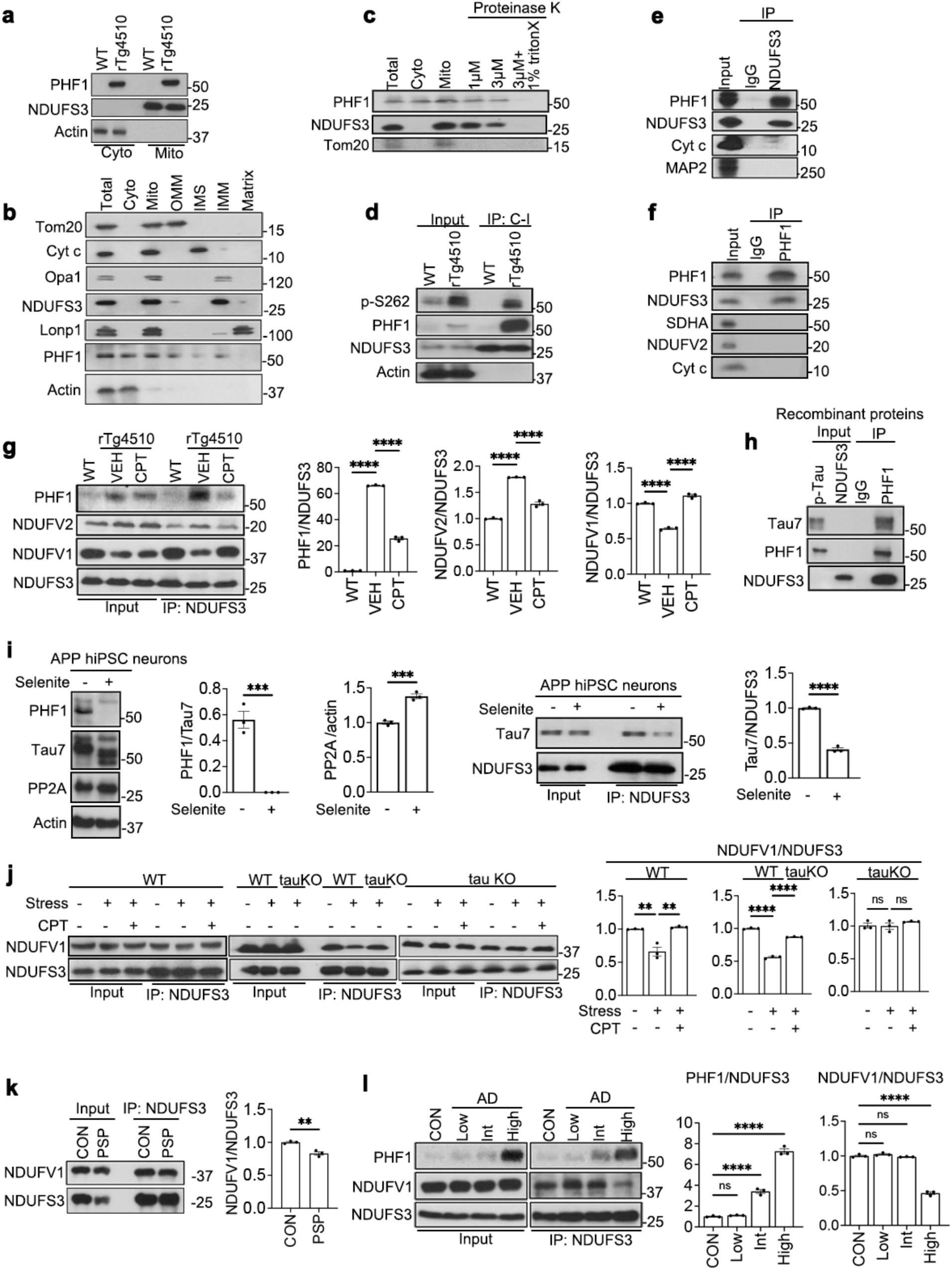
tau is localized inside mitochondria and interacts with NDUFS3 to alter RET-related protein-protein interactions. **a**, WB showing the presence of p-tau inside mitochondria of rTg4510 mouse brain. **b**, mitochondrial fractionation showing the presence of p-tau in mitochondrial subfractions. Tom20, Cyt *c*, Opa1, NDUFS3, and LonP1 are outer membrane (OMM), intermembrane space (IMS), inner membrane (IMM), C-I, and matrix markers, respectively. **c**, PK digestion of purified mitochondria from rTg4510 mice demonstrating the presence of p-tau inside mitochondria. **d**, Co-IP assay showing the presence of p-tau in affinity-purified C-I from rTg4510 mice. **e**, IP with NDUFS3 antibody showing specific interaction of p-tau with NDUFS3 in rTg4510 brain. **f**, Reverse IP with PHF-1 antibody showing specific interaction of NDUFS3 with p-tau in rTg4510 brain. **g**, Co-IP assay and quantification showing increased PHF-1 tau/NDUFS3 and NDUFV2/NDUFS3 and decreased NDUFV1/NDUFS3 interactions in rTg4510 mouse brain and their rescue by CPT. **h**, Co-IP assay showing *in vitro* binding between p-tau and NDUFS3 recombinant proteins. **i**, WBs showing dephosphorylation of tau and reduced tau/NDUFS3 co-IP in APP hiPSC-derived neurons treated with the PP2A activator sodium selenite. **j**, Co-IP and quantification showing altered NDUFS3/NDUFV1 interaction in WT or tau KO mice exposed to heat stress, and with or without CPT treatment. **k**, Analysis of NDUFS3/NDUFV1 interaction in PSP patient brain samples by co-IP. **l**, Co-IP assay and quantification of NDUFS3/NDUFV1 and PHF-1 tau/NDUFS3 interactions in brain samples of patients with different degrees of AD (low, intermediate, and high). All data are means ± SEM; statistical significance was determined by one-way ANOVA with Tukey’s post hoc test (**g, j, l**), or two-tailed unpaired Student’s *t* test (**i**,**k**). *p<0.05; **p<0.01; ***p<0.001; ****p<0.0001. Each data point represents an independent experimental repeat.

Next, we tested whether p-tau directly associates with C-I. Using a well-established affinity purification method^67^, we isolated C-I from rTg4510 brain tissues and detected p-tau in purified samples (Fig. 3d). Co-immunoprecipitation (co-IP) experiments in rTg4510 brain lysates revealed p-tau interaction with NDUFS3 (Fig. 3e-f) - the C-I subunit critical for RET and targeted by CPT^42^. The specificity was demonstrated by the lack of PHF-1 p-tau interaction with another C-I protein NDUFV2 or other ETC proteins (Fig. 3f). Proximity ligation assay (PLA), a technique that uses a combination of antibodies and rolling circle amplification to visualize and quantify protein-protein interactions, further verified the interaction between p-tau and NDUFS3, which was reduced by CPT treatment (Extended Data Fig. 7b). Similarly, CPT weakened the interaction between PHF-1 p-tau and NDUFS3 in co-IP experiments (Fig. 3g), suggesting that CPT and p-tau may compete for binding to NDUFS3.

To determine whether the interaction between p-tau and NDUFS3 is direct, we performed *in vitro* binding assays using recombinant proteins, which confirmed a robust interaction (Fig. 3h). In a hiPSC-derived neuronal model of AD carrying APP duplication^68^, activating phosphatase 2A to dephosphorylate tau significantly reduced the interaction between tau and NDUFS3 in co-IP assays (Fig. 3i). These results demonstrate that tau phosphorylation promotes its interaction with NDUFS3.

Although soluble tau is an intrinsically disordered protein that adopts largely random coil structures^69^, AlphaFold-Multimer^70,71^ was able to predict specific interactions between tau and NDUFS3. These included β-strand interactions between NDUFS3 residues 72-77 (VQQVQV) and tau C-terminal residues 375-381 (KLTFREN), as well as electrostatic interactions between phosphorylated serine residues in the PHF-1 epitope (p-S396, p-S400, and p-S404) and positively charged residues (R43, R49, and R245) in NDUFS3 (Extended Data Fig. 7c,d). R43 and R49 of NDUFS3 are surface-exposed in the cryo-EM structure of mouse C-I^72^, making them accessible to p-tau. To test these predictions, we mutated NDUFS3 residues 72-77 (VQQVQV) individually to Pro to disrupt β-strand interactions, and mutated R43 and R49 individually or together to Ala to impair electrostatic interactions. Both types of interactions were important for NDUFS3/p-tau binding, with residues such as R49 and Q73 playing a more significant role in binding PHF-1 tau and inducing RET-ROS (Extended Data Fig. 7e,f). AlphaFold predictions explain the importance of Q73, which engages in extensive hydrogen bonding with tau residues H388, D387, and N381, as well as NDUFS3 residue C86 (Extended Data Fig. 7g). The neighboring V72 and Q74 residues likely stabilize these interactions. R49A and R43/49A double mutants of NDUFS3 disrupted NDUFS3/p-tau interaction, highlighting the critical role of R49/p-S396 electrostatic interactions.

To further verify p-tau’s association with C-I, we performed BN-PAGE of rTg4510 brain mitochondria and observed a distribution pattern of endogenous PHF-1 tau resembling that of NDUFS3 (Extended Data Fig. 8a). Next, 2D-gel electrophoresis was performed in which mitochondrial protein complexes were first separated by BN-PAGE in the 1^st^ dimension, followed by SDS-PAGE in the 2^nd^ dimension. PHF-1 tau was detected in BN-PAGE-separated C-I complexes, as indicated by the presence of NDUFS3 but the absence of C-II protein SDHA (Extended Data Fig. 8b). BN-PAGE in-gel activity assays showed that rTg4510 and control mice had comparable C-I activities (Extended Data Fig. 8c). Additional co-IP experiments using PK-treated mitochondria confirmed the specificity of the tau-NDUFS3 interaction (Extended Data Fig. 8d). Furthermore, purification of C-I from rTg4510 mouse brains using ion exchange and size exclusion chromatography revealed PHF-1 tau co-purifying with C-I (Extended Data Fig. 8e).

Finally, we examined how p-tau affects RET-related protein-protein interactions within C-I. RET activation has been associated with decreased NDUFS3/NDUFV1 interaction and increased NDUFS3/NDUFV2 interaction, indicating conformational changes within C-I that mediate RET^42,43^. In rTg4510 mouse brains, correlating with RET activation, the NDUFS3/NDUFV1 interaction was decreased and NDUFS3/NDUFV2 interaction was increased; both changes were reversed by CPT treatment (Fig. 3g). Similarly, heat stress-induced RET activation in WT mice was associated with decreased NDUFS3/NDUFV1 interaction (Fig. 3j), as confirmed using antibodies verified for specificity (Extended Data Fig. 8f). This effect was absent in tau KO mice (Fig. 3j) and tau KO flies (Extended Data Fig. 8g) subjected to heat stress and with or without CPT treatment, suggesting that tau is necessary for these RET-related changes within C-I.

To further validate these findings, we examined patient samples. Compared to normal brain, the PSP brain tissues showed increased PHF-1 tau (Extended Data Fig. 8h), decreased NDUFS3/NDUFV1 interaction (Fig. 3k), and enhanced p-tau/NDUFS3 interaction (Extended Data Fig. 8i). Similarly, AD brain tissues exhibited increased p-tau/NDUFS3 interaction and reduced NDUFS3/NDUFV1 interaction (Fig. 3l). Moreover, tau-WT inhibited the NDUFS3/NDUFV1 interaction in a phosphorylation-dependent manner (Extended Data Fig. 8j). Taken together, these findings demonstrate that tau acts in a phosphorylation-dependent manner to directly interact with NDUFS3, altering RET-related protein-protein interactions within C-I. Through these interactions, tau promotes RET activity, providing insight into its detrimental role in stress response and neurodegeneration.

### Inhibiting the mitochondrial entry of tau diminishes its RET-promoting effect

To further substantiate the intramitochondrial action of tau in promoting RET, we explored ways to inhibit its mitochondrial entry. Although tau has been reported to localize to mitochondria and interact with various mitochondrial proteins, the mechanism by which it enters mitochondria remains unknown. Proteins such as voltage-dependent anion channel (VDAC1) and molecular chaperones Hsp70 and Hsp90 - known to facilitate mitochondrial protein import^73, 74^ - have been shown to interact with tau^75–77^. We found that mitochondria purified from HEK293 cells treated with specific inhibitors of VDAC ^78^, Hsp70 ^79^, or Hsp90 ^80^ were compromised in importing recombinant PHF-1 tau and exhibited lower RET activity than control mitochondria after treatment with PHF-1 tau (Extended Data Fig. 9a,b), despite similar amounts of total PHF-1 tau in the reaction and unaltered basal RET activity of inhibitor-treated mitochondria (i.e., without PHF1 tau import) (Extended Data Fig. 9c,d). These findings demonstrate that PHF-1 tau needs to enter mitochondria to promote RET.

Additionally, genetic manipulation of VDAC1, Hsp70, or Hsp90 via RNAi, which reduced target protein expression (Extended Data Fig. 9e), produced effects similar to those observed with chemical inhibitors. RNAi-mediated knockdown of these proteins hindered tau’s mitochondrial entry and its RET-promoting activity (Extended Data Fig. 9f,g), while leaving basal mitochondrial RET activity unaffected (Extended Data Fig. 9h). These results further support the involvement of VDAC1, Hsp70, and Hsp90 in regulating tau’s mitochondrial entry and confirm an intramitochondrial role for tau in promoting RET.

### Genetic or pharmacological inhibition of RET rescues tauopathy in a fly model

Our findings suggest that the regulation of RET is a physiological function of tau, which becomes exacerbated in tauopathy. To investigate the contribution of RET to tau toxicity, we evaluated its inhibition in tauopathy models. Transgenic flies with pan-neuronal expression of FTD-associated tau-R406W exhibited progressive neurodegeneration and shortened lifespan. This model has been widely used to study the mechanisms underlying phospho-regulation of tau toxicity^65^. Treating these flies at the adult stage with food containing 10 µM CPT - a regimen known to effectively inhibit RET^43^ - rescued locomotor deficits and extended lifespan (Extended Data Fig. 10a,b). These tau-R406W flies also presented learning and memory defects as detected in the aversive taste memory assay, in which the proboscis extension reflex (PER) in response to sucrose indicates how well the flies remember an aversive taste co-applied with sucrose during training^81^. CPT treatment rescued the memory deficits (Extended Data Fig. 10c) and neuronal loss (Extended Data Fig. 10d,e), while also restoring proteostasis (Extended Data Fig. 10d,e). Furthermore, CPT treatment reduced p-S262 and PHF-1 tau levels (Extended Data Fig. 10f,g), a point we will address later.

To complement the pharmacological approach, we employed a genetic strategy. Partial knockdown of NDUFS3 - the target site of CPT - has previously been shown to mimic the effect of CPT *in vivo*^42,43^. Using NDUFS3 RNAi, we observed similar rescue effects as CPT on locomotor activity, lifespan, learning and memory, and p-tau level in tau-R406W flies (Extended Data Fig. 10h-k).

Importantly, inhibition of VDAC, Hsp70, or Hsp90 diminished PHF-1 tau entry into mitochondria without affecting total PHF-1 tau level (Extended Data Fig. 11a), reduced RET activity (Extended Data Fig. 11b), and rescued learning and memory deficit (Extended Data Fig. 11c-e) as well as locomotor activity (Extended Data Fig. 11f) in tau-R406W flies. Notably, inhibition of VDAC, Hsp70, or Hsp90 had no significant impact on basal RET activity or behavior in control flies under the same experimental conditions (Extended Data Fig. 11g,h). These results strongly support an intramitochondrial role of tau in RET regulation *in vivo* and highlight the pathological consequences of increased RET in tauopathy.

### Inhibition of RET rescues neuroinflammation, neurodegeneration, and cognitive deficits in a mouse tauopathy model

To evaluate the impact of RET inhibition in a mammalian tauopathy model, we used rTg4510 mice, which develop tau neuropathology by 2 months of age and cognitive deficits at 4-5 months^55^. Starting at 4 months of age, rTg4510 mice were treated with CPT via daily oral gavage for 2–4 months. CPT-treated rTg4510 mice exhibited significant improvement compared to vehicle-treated controls across a range of behavioral tests. In the Morris water maze (MWM) assay for spatial memory, CPT-treated rTg4510 mice showed significantly shorter escape latencies to find the hidden platform on training days 4 and 5, along with increased target quadrant occupancy during the probe trial (Extended Data Fig. 12a,b). In the contextual fear conditioning test for associative learning, CPT-treated rTg4510 mice spent significantly more time freezing when exposed to an environment where they had previously received a mild foot shock (Extended Data Fig. 12c). Hyperactivity, as measured by total distance traveled in the open field test, was rescued by CPT treatment (Extended Data Fig. 12d). Similarly, in the EPM assay for anxiety-like behavior, rTg4510 mice spent significantly more time in the open arm, indicating subdued fear or anxiety, which was also rescued by CPT treatment (Extended Data Fig. 12e,f). CPT exhibited a dose-dependent response across these assays (Extended Data Fig. 12c-f).

Consistent with previous reports showing CPT does not affect cognition or behavior in WT mice^42,43^, CPT treatment had no observable effects on C57BL/6 control mice in fear conditioning, open field, or EPM assays (Extended Data Fig. 12g-j). Additionally, CPT treatment reduced brain ROS levels and increased NAD^+^/NADH ratio in rTg4510 mice, demonstrating RET target engagement, while FET activity remained unaffected (Extended Data Fig. 12k-n). These findings indicate that RET contributes significantly to the behavioral deficits caused by tau-P301L in mammals. Importantly, behavioral differences between CPT-treated and vehicle-treated rTg4510 mice remained significant when males and females were analyzed separately, ruling out sex-specific effects (Extended Data Fig. 13a-l).

We next examined the effect of RET inhibition on tau-induced neurodegeneration. Extensive brain atrophy in rTg4510 mice, a hallmark of tauopathy, can be detected via MRI^82^. CPT treatment mitigated brain atrophy, as evidenced by increased cortical thickness, hippocampal and total brain volumes, and reduced ventricle size compared to vehicle-treated rTg4510 mice (Fig. 4a,b). Notably, the olfactory bulb area was not affected in rTg4510 mice (Extended Data Fig. 14a,b). Consistent with MRI findings, immunostaining of the cortex and hippocampus revealed that CPT treatment reduced markers of DNA-damage (γH2AX, Extended Data Fig. 14c) and apoptosis (cleaved caspase 3, Extended Data Fig. 14d) while preserving NeuN-positive neurons (Fig. 4c,d), indicating attenuation of tau-induced neurodegeneration. Furthermore, CPT treatment reduced Thio-S-positive tau aggregates (Extended Data Fig. 14e) and p-tau levels (Extended Data Fig. 14f-i), a point we will address later.

**Fig. 4.**
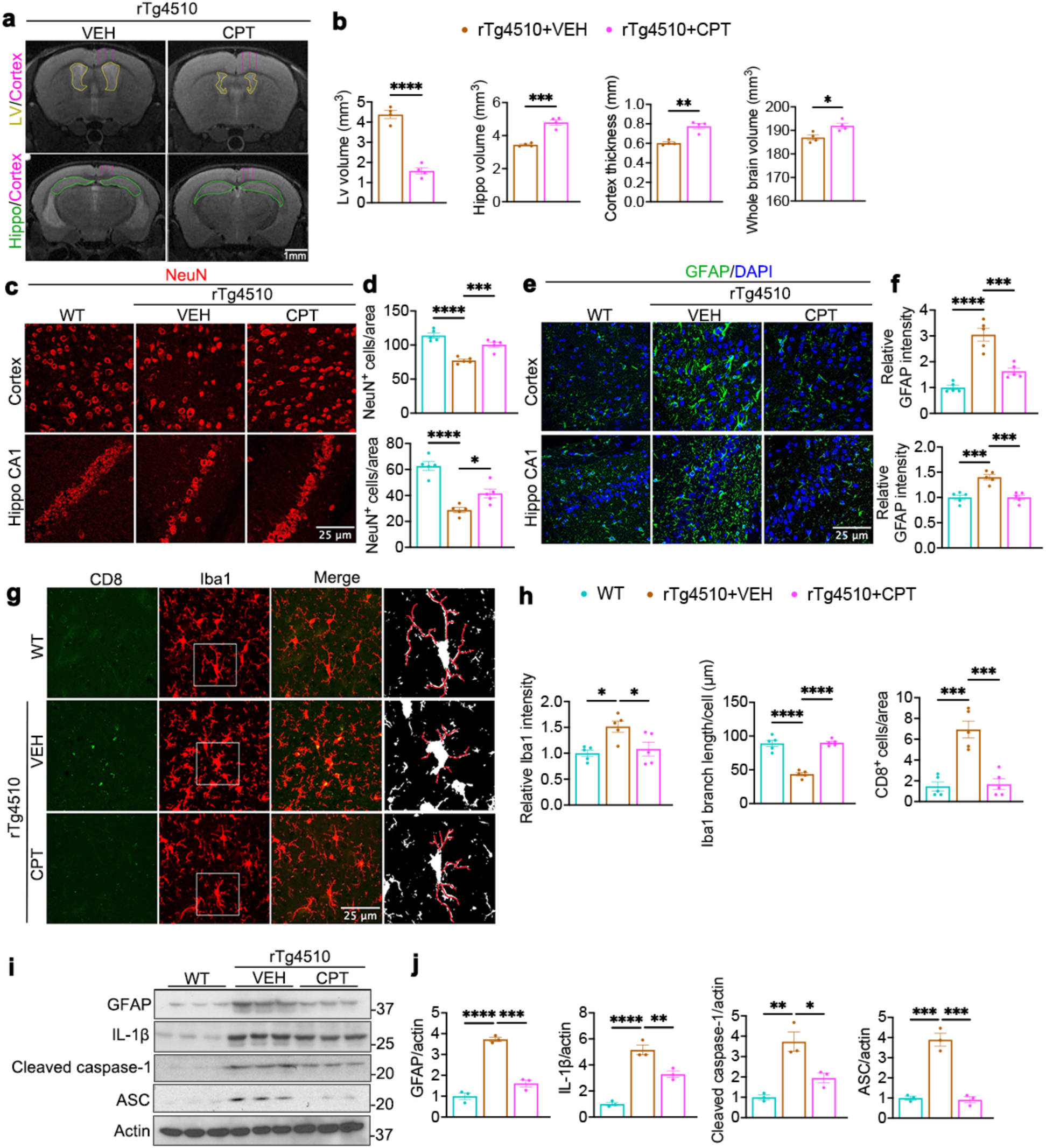
Effect of RET inhibition on neuroinflammation and neurodegeneration in rTg4510 tauopathy mice. **a, b**, Representative MRI images (**a**) and data quantification (**b**) showing effect of RET inhibition by CPT on brain atrophy in 8-month-old rTg4510 mice. Volume information was obtained from the lateral ventricles (Lv), hippocampus (Hippo), and whole brain. Arrows in the cortical area mark cortical thickness. **c, d,** Representative images (**c**) and data quantification (**d**) of NeuN staining of neurons in the Hippo CA1 and cortex regions. (**e, f**) Representative images (**e**) and data quantification (**f**) of GFAP staining in the hippocampal CA1 and cortex regions. (**g, h**) Representative images (**g**) and data quantification (**h**) of CD8 and Iba1 immunostaining in the cortex region. (**i, j**) Immunoblots (**i**) and data quantification (**j**) of inflammation-related markers GFAP, IL-1β, cleaved caspase-1, and ASC in the cortex tissues of 8-month-old mice. WT: wildtype littermates of rTg4510. VEH and CPT: rTg4510 mice treated with vehicle (VEH) or CPT. All data are means ± SEM; statistical significance was determined by one-way ANOVA with Tukey’s post hoc test (**d, f, h, j**), or two-tailed unpaired Student’s *t* test (**b**). * p<0.05; **p<0.01; ***p<0.001; ****p<0.0001 (**b**: n=4 mice/group; **d, f, h**: n=5 mice/group, each data point represents an average of 6 images/mouse; for Iba1 branch length measurement, each data point represents an average of 10 cells/mouse; **j**: n=3 mice/group).

Reduced cytoplasmic occupancy of mitochondria within neuronal soma was previously seen in rTg4510 mice^83^. A similar defect was identified in PSP human samples (Extended Data Fig. 15b), indicating a conserved tauopathy phenotype potentially linked to RET activation. Notably, this mitochondrial defect was reversed by CPT treatment (Extended Data Fig. 15a).

To further characterize the effect of CPT on tau-induced mitochondrial abnormalities, we analyzed mitochondrial dynamics. In rTg4510 mouse brains, levels of the mitochondrial fission regulator Drp1 were increased, while levels of fusion regulators Opa1 and Mfn2 were decreased. CPT treatment alleviated these defects (Extended Data Fig. 15c). Synaptic health is particularly vulnerable to mitochondrial dysfunction^84^, as demonstrated by reduced levels of synaptic markers Glu2B, PSD-95, and Synapsin-1 in rTg4510 mice. These reductions were also rescued by CPT treatment (Extended Data Fig. 15d).

Neuroinflammation is a pathological feature of tauopathies and is thought to contribute to tau pathology^85^. Glial activation and T cell infiltration have been observed in the brains of tauopathy patients^86^ and mouse models^85,87^. In rTg4510 mice, CPT treatment reduced microglial and astrocyte activation, as demonstrated by Iba1 and GFAP immunostaining, respectively. Cytotoxic T cell infiltration, assessed via CD8 immunostaining, was also diminished (Fig. 4e-h). These results suggest that neuroinflammation in the rTg4510 model is partially driven by RET. Consistent with this, western blot and qRT-PCR analyses revealed upregulation of certain neuroinflammation-related genes in rTg4510 mice, which was attenuated by CPT treatment (Fig. 4i, j; Extended Data Fig. 15e). The mRNA levels of *Alx*, *Ank*, *NF-kB,* and *Cox6a2* were unaffected by CPT treatment or treatment with the RET inducer DES (Extended Data Fig. 15f), suggesting that the upregulation of these genes in rTg4510 mice may be RET-independent. Overall, these findings suggest that RET activation contributes to key aspects of neuroinflammation in rTg4510 tauopathy mice, while CPT mitigates these pathological processes.

### Inhibition of RET is protective in human iPSC-derived neuronal models of tauopathies

To investigate the effects of RET inhibition on tau toxicity in human neurons, we used patient hiPSC-derived neurons carrying the P301L tau mutation alongside isogenic WT controls. Under standard culture conditions, no significant differences in differentiation or survival were observed between control and tau-P301L neurons. To test whether tau-P301L neurons might exhibit increased vulnerability to cellular stressors, we applied oxidative stress (via H_2_O_2_) and disrupted the NAD^+^/NADH ratio (via the NAD^+^ synthase inhibitor FK866), both of which are relevant to RET activation. Control neurons exhibited baseline H_2_O_2_ concentrations at around 100 µM (Extended Data Fig. 16a), while oxidative stress and cytotoxicity are typically induced in cells with H_2_O_2_ concentrations between 100 - 500 µM^88^. After testing a range of 25 - 400 µM, we selected 200 µM H_2_O_2_ for subsequent analyses, as it reduced tau-P301L neuronal viability by ∼50% without significantly affecting control neurons (Extended Data Fig. 16b). Similarly, 200 nM FK866 was chosen because it caused a comparable reduction in tau-P301L neuronal viability while sparing control neurons (Extended Data Fig. 16c). CPT treatment partially rescued tau-P301L neurons from the toxic effects of H_2_O_2_ and FK866 on (Fig. 5a). This rescue likely reflects CPT’s ability to block endogenous RET-induced changes in ROS and NAD⁺/NADH ratios, while leaving much of the exogenous stress unaltered. While control neurons did not show significant changes of steady state ROS in response to the stressors, tau-P301L neurons had increased ROS, which was partially rescued by CPT (Extended Data Fig. 16d). Similarly, stress-induced cleaved caspase-3 levels were elevated in tau-P301L neurons and partially rescued by CPT (Extended Data Fig. 16e). Tau-P301L neurons treated with stressors also showed increased colocalization of the autophagy receptor p62 with mitochondria, indicating a defect in mitophagy flux, which CPT ameliorated (Extended Data Fig. 16f). At the synaptic level, tau-P301L neurons exhibited fewer Synapsin-positive puncta compared to controls, with stressors further reducing Synapsin levels - a defect that CPT reversed (Extended Data Fig. 16g). These results confirm that mutant tau increases stress vulnerability in tau-P301L neurons and that CPT mitigates most of these cellular abnormalities (Extended Data Fig. 16d–g) by targeting RET (as indicated by changes in ROS and NAD⁺/NADH ratios; Extended Data Fig. 1c).

**Fig. 5.**
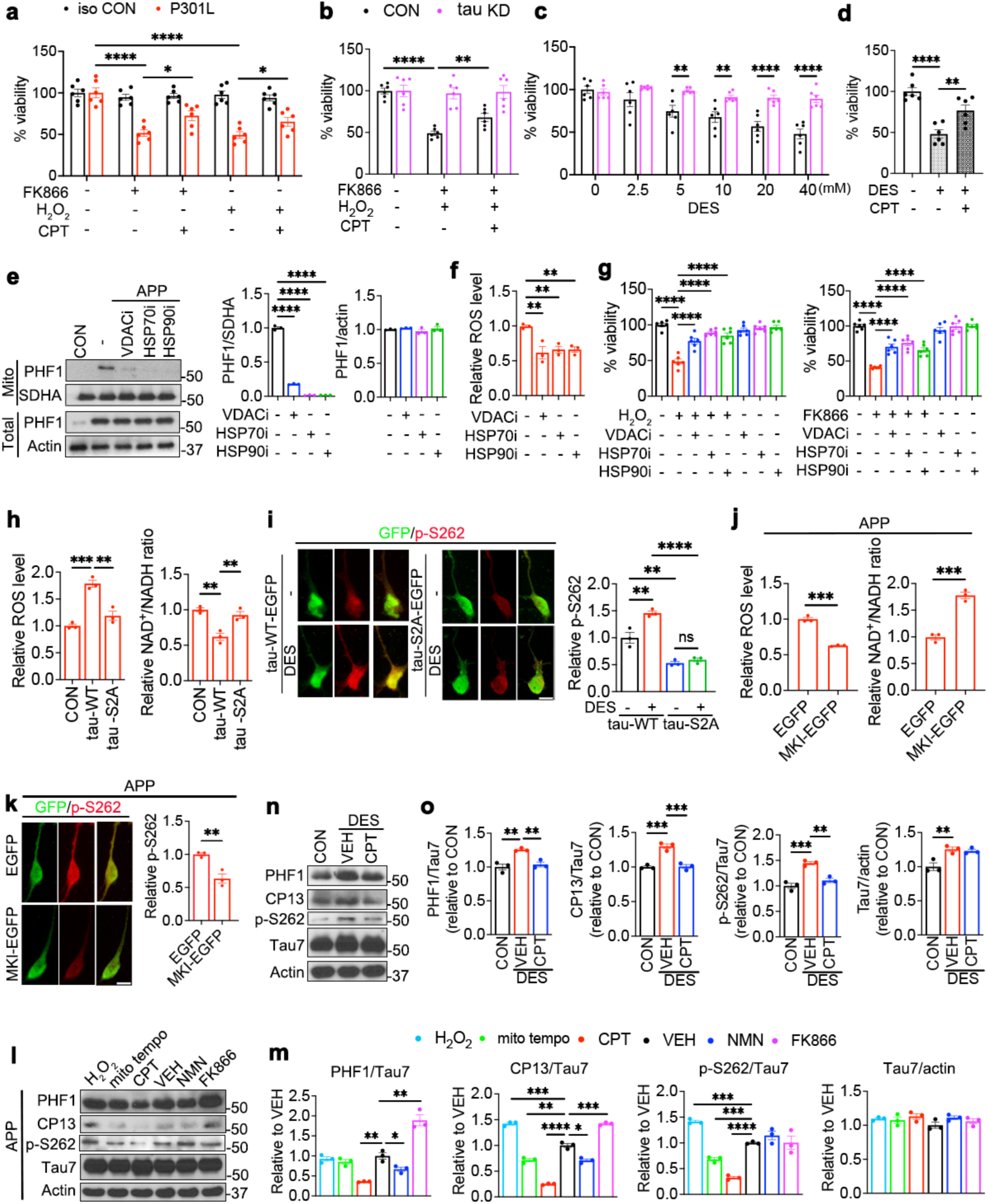
Effect of RET inhibition and tau knockdown on tau phosphorylation and stress response in hiPSC-derived neuronal models of tauopathies. **a**, Effect of H_2_O_2_ and FK866 on the viability of tau-P301L hiPSC neurons and isogenic wildtype controls after 24h treatment, and the rescuing effect of CPT. **b**, Effect of H_2_O_2_ and FK866 co-treatment on the viability of control and tau KD hiPSC neurons and the rescuing effect of CPT. **c, d**, Effect of DES treatment on the viability of control and tau KD hiPSC neurons (**c**) and the rescuing effect of CPT in DES (20 mM) treated control hiPSC neurons (**d**). **e**-**g**, WB and quantification showing the effect of inhibiting VDAC, Hsp70, or Hsp90 on PHF-1 tau entry into mitochondria without affecting total PHF-1 tau levels (**e**), and quantification of the effect of inhibitor treatment on RET activity (**f**) and stress sensitivity (**g**) of APP hiPSC neurons. **h**, Measurement of RET ROS and NAD^+^/NADH in purified mitochondria from control and tau-WT-EGFP or tau-S2A-EGFP-transfected normal hiPSC neurons. **i**, Representative images and quantification of p-S262 tau in tau-WT-EGFP or tau-S2A-EGFP transfected control hiPSC neurons with or without DES treatment. **j**, Measurement of RET ROS and NAD^+^/NADH in purified mitochondria from EGFP or MKI-EGFP transduced APP hiPSC neurons. **k**, Representative images and quantification of p-S262 tau in EGFP or MKI-EGFP transfected APP hiPSC neurons. **l, m**, Immunoblots (**l**) and quantification (**m**) showing effects of the various treatments on normalized levels of p-tau species in APP hiPSC neurons. **n, o**, Immunoblots (**n**) and quantification (**o**) showing effect of DES or DES/CPT co-treatment on normalized levels of p-tau species in control hiPSC neurons. All data are means ± SEM; statistical significance was determined by one-way ANOVA with Tukey’s post hoc test (**a, b, d, g, h, i, o**), two-way ANOVA with Tukey’s post hoc test (**c**), two-tailed unpaired Student’s *t* test (**j, k**), or one-way ANOVA with Dunnett’s multiple test (**e, f**, **m**). *p<0.05; **p<0.01; ***p<0.001; ****p<0.0001 (**a, b, c, d, g**: n=6/group from 3 biological replicates and 2 wells/experiment; **e, f, h, j, m, o**: n=3 biological replicates; **i, k**: n=3 biological replicates, and each data point represents an average of 6 cells/experiment).

To further validate the role of RET in tau-mediated neurotoxicity, we conducted two additional experiments using control and tau KD neurons. First, we tested whether combining H_2_O_2_ (200 µM) and FK866 (200 nM) to simultaneously increase ROS and decrease NAD^+^/NADH ratio, thus mimicking RET activation, would further compromise neuronal health. Indeed, this combined treatment dose-dependently reduced the viability of control neurons, while tau KD neurons were unaffected (Extended Data Fig. 16h; Fig. 5b). Second, we evaluated the effect of RET activation via DES treatment, which induced ROS and NAD⁺/NADH ratio changes that were inhibited by genetic RET knockdown using NDUFS3 RNAi (Extended Data Fig. 17a). Similar to the combined stressor treatment, DES reduced the viability of control neurons but not tau KD neurons, further supporting the role of tau in mediating RET activation and its detrimental effects (Fig. 5c,d). CPT also protected neurons from both treatments (Fig. 5b,d).

Control neurons subjected to the combined H_2_O_2_/FK866 treatment showed increased ROS levels, cleaved caspase-3, and p62-positive puncta associated with mitochondria, all of which were rescued by CPT or tau KD (Extended Data Fig. 17b-d). Comparable results were observed in APP hiPSC-derived neuron models of AD, where increased ROS and decreased NAD⁺/NADH ratios were rescued by genetic inhibition of RET via NDUFS3 RNAi (Extended Data Fig. 17e–h). Tau knockdown similarly mitigated RET activation in APP neurons (Extended Data Fig. 17i–k) and in transgenic flies overexpressing human APP (Extended Data Fig. 17l,m), demonstrating that endogenous tau acts downstream of APP to activate RET in AD contexts.

Notably, inhibition of VDAC, Hsp70, or Hsp90 reduced PHF-1 tau entry into mitochondria without affecting total PHF-1 tau levels (Fig. 5e). This in turn decreased RET activation (Fig. 5f) and rescued the sensitivity of APP neurons to H_2_O_2_ and FK866 (Fig. 5g), indicating that endogenous p-tau acts intramitochondrially to drive RET-related toxicity. Genetic knockdown of VDAC1, Hsp70, or Hsp90 similarly reduced PHF-1 tau entry into mitochondria without affecting total PHF-1 tau level (Extended Data Fig. 18a-c). Inhibition of VDAC, Hsp70, or Hsp90 did not affect basal RET activity (Extended Data Fig. 18d) or stressor sensitivity in control neurons (Extended Data Fig. 18e). Together, these findings establish that RET inhibition mitigates tau-mediated neurotoxicity and provides protective benefits in hiPSC models of both primary (tau-P301L) and secondary (APP) tauopathies.

### Feedforward regulation between RET and tau phosphorylation in a vicious cycle

To explore whether phospho-tau regulates RET in human neurons, we transfected control neurons with tau constructs. Tau-WT strongly promoted RET activation, while tau-S2A - a mutant lacking phosphorylation sites at S262 and S356 - failed to do so (Fig. 5h). DES treatment induced a significant increase in p-S262 tau in tau-WT neurons but not in tau-S2A neurons (Fig. 5i). Furthermore, APP hiPSC-derived neurons transfected with EGFP-fused MKI^89^, a peptide inhibitor of PAR-1/MARK kinases that phosphorylates the S262 and S356, showed attenuated RET activation (Fig. 5j,k). These findings highlight the critical role of tau phosphorylation by PAR-1/MARK at S262 and S356 in RET activation. Consistently, PAR-1 RNAi inhibited RET in tau-R406W transgenic flies (Extended Data Fig. 17n,o).

Interestingly, CPT treatment reduced p-tau levels without affecting total tau in tau-P301L neurons (Extended Data Fig. 19a) and APP neurons (Fig. 5l, m). Similarly, CPT reduced p-tau in tau-R406W flies (Extended Data Fig. 10f,g) and rTg4510 mice (Extended Data Fig. 14h,i). Genetic inhibition of RET by NDUFS3 RNAi also lowered p-tau levels in APP neurons (Extended Data Fig. 19b). These results suggest the existence of a feedforward loop between RET activation and tau phosphorylation. To test this hypothesis further, RET was induced in control neurons using DES, which promoted tau phosphorylation. CPT treatment reduced this DES-induced tau phosphorylation (Fig. 5n, o). Acute DES treatment in WT mice also increased p-tau accumulation in the brain mitochondrial fraction (Extended Data Fig. 19c).

To probe the mechanisms driving RET-related p-tau changes, we examined tau kinases and phosphatases. In the rTg4510 mouse brains, phosphorylation of MARK at its activation site in the kinase domain (corresponding to T408 in PAR-1^90^) was increased, while phosphorylation of GSK-3β at its inhibitory S9 site was decreased, indicating activation of both tau kinases. CPT treatment inhibited these kinase activations. In contrast, the level of the PP2A phosphatase remained unchanged (Extended Data Fig. 19d). These findings implicate PAR-1/MARK and GSK-3β in RET-related tau hyperphosphorylation.

We also tested the relative contribution of the two outcomes of RET - ROS increase and NAD^+^/NADH imbalance - to RET-induced tau phosphorylation. While RET-induced tau phosphorylation at the CP13 site was reduced by either antioxidant treatment or supplementation with the NAD^+^ precursor NMN, the PHF-1 site responded to NMN but not antioxidants (Fig. 5l, m; Extended Data Fig. 19e). These results suggest that RET-induced changes in ROS and NAD⁺/NADH ratios regulate tau phosphorylation through distinct pathways, contributing to site-specific tau phosphorylation. Together, these findings reveal a self-perpetuating pathological loop in which p-tau drives RET activation, while RET-induced ROS and NAD⁺/NADH imbalances feed forward to further increase tau phosphorylation. This vicious cycle exacerbates tau pathology. Inhibition of RET with CPT effectively disrupts this cycle, offering therapeutic benefits.

## Discussion

Despite significant efforts to develop treatments for neurodegenerative diseases, therapeutic success has been limited, largely due to an incomplete understanding of underlying disease mechanisms. In tauopathies, while considerable progress has been made in elucidating tau’s role in microtubule-related function and dysfunction, the normal physiological function of tau in the adult brain - particularly in relation to disease - remains uncertain. This knowledge gap has hindered efforts to uncover the core mechanisms of tau-induced neurotoxicity and develop effective therapies. Here, we identify a previously unrecognized physiological function of tau as a regulator of RET through its direct interaction with the mitochondrial C-I protein NDUFS3. Several lines of evidence strongly support tau’s intramitochondrial role in promoting RET: blocking tau’s mitochondrial entry diminishes RET activation, and mutations in NDUFS3 that disrupt tau interaction reduce RET activity. Using AlphaFold-based structural predictions combined with experimental validation, we provide mechanistic insights into tau’s interaction with C-I and regulation of RET. Furthermore, RET activation is shown to be dependent on p-tau - widely implicated in tau pathogenesis, and is deregulated across tauopathy models, human iPSC-derived neurons, and patient brain samples. Crucially, therapeutic inhibition of RET mitigates tau-induced neurotoxicity in multiple models, without observable detrimental effects on normal animals. These findings suggest that RET inhibition holds promise as a therapeutic strategy not only for tauopathies but potentially for other brain diseases characterized by aberrant tau phosphorylation and mitochondrial dysfunction.

A key finding of this study is the identification of a feedforward loop between RET and tau phosphorylation (Extended Data **Fig. 20**). While phospho-tau promotes RET activation, RET-induced changes in ROS and NAD^+^/NADH ratios further enhance tau phosphorylation, creating a self-perpetuating pathological cycle. RET-induced tau phosphorylation is mediated by distinct signaling pathways: ROS appears to act through stress and inflammatory pathways, while NAD^+^/NADH imbalance likely influences tau acetylation and phosphorylation via Sirtuin signaling. These insights begin to unravel the molecular mechanisms underlying RET-induced tau hyperphosphorylation and highlight site-specific effects of tau phosphorylation on RET. For example, phosphorylation at the PHF-1 site is particularly critical for tau’s interaction with NDUFS3 and RET promotion, while mutations at the AT8 (S202) site have little impact on RET.

Our findings demonstrate that RET inhibition provides broad therapeutic benefits in tauopathy models. The RET inhibitor CPT effectively disrupts the pathological loop between RET and tau phosphorylation, ameliorating neurotoxicity across species. In fly and mouse models of tauopathy, CPT treatment rescues behavioral deficits, reduces neuroinflammation, and mitigates neurodegeneration. In hiPSC-derived neurons carrying pathogenic tau mutations, CPT protects against stress-induced cellular abnormalities, including ROS overproduction, mitochondrial dysfunction, synaptic deficits, and impaired mitophagy. Notably, RET inhibition reduces p-tau levels without affecting total tau, suggesting that targeting RET may avoid potential adverse effects associated with complete tau depletion.

The therapeutic potential of RET inhibition extends beyond tauopathies. Mitochondrial dysfunction is a hallmark of aging and age-related diseases^91^, and RET is emerging as a driver of aging and aging-related pathology^43^. Our findings resonate with the geroscience hypothesis, which posits that targeting aging mechanisms may prevent or mitigate multiple age-related diseases^92,93^. Evidence supporting this hypothesis includes the association of C-I subunit abnormalities (including NDUFS3) with longevity and neuroprotection in experimental models^94, 95^; lifespan extension by partial knockdown of NDUFS3 and select C-I subunits in worms^96^ and flies^97^; neuroprotection by mild inhibition of C-I in mouse AD models^98^; and association of *NDUFS3* with late onset AD in human genome-wide association studies, with NDUFS3 RNAi being protective against AD related toxicity^99^. These observations suggest that mild inhibition of NDUFS3 may confer therapeutic benefits by inhibiting pathological RET without compromising essential mitochondrial functions.

This study sheds light on tau’s normal physiological role in regulating RET and its pathological transition in disease. Under normal conditions, tau may act through RET to mediate stress adaptation, particularly during aging. However, persistent stress stimuli or aging-related changes can lead to supraphysiological levels of RET and p-tau, triggering a vicious cycle that drives neurotoxicity (Extended Data **Fig. 20**). The observation that RET and tau phosphorylation are upregulated during aging and stress suggests that tauopathies may arise from the deregulation of normal physiological processes. Our findings also suggest that therapeutic strategies targeting tau phosphorylation must account for site-specific effects. Not all phospho-tau sites contribute equally to RET regulation or tau toxicity, as evidenced by the differential effects of mutations at the PHF-1, AT100, and S262/S356 sites. Further studies are needed to identify modulators that selectively target pathological tau species while preserving normal tau function.

The identification of tau as a modulator of RET has broader implications for neurodegenerative diseases. Aberrant tau phosphorylation and mitochondrial dysfunction co-occur in numerous brain disorders, including AD, frontotemporal dementia (FTD), and Parkinson’s disease. Our results suggest that RET may serve as a common pathogenic mechanism linking tau abnormalities to mitochondrial dysfunction across diseases. For example, in AD models, endogenous tau acts downstream of APP to activate RET, and RET inhibition mitigates APP-related neurotoxicity. This highlights the potential of RET-targeted therapies to address both primary tauopathies and secondary tauopathies associated with other disease contexts.

Future studies should explore whether other FTD-related tau mutations and disease-associated tau variants similarly promote RET. Additionally, the cell-type specificity of RET and tau phosphorylation in different neurodegenerative diseases remains an open question. It is possible that certain cell types are more vulnerable to RET-induced changes in ROS and NAD^+^/NADH metabolism, or that disease-specific factors selectively upregulate RET and phospho-tau. Addressing these questions will deepen our understanding of tau’s role in disease and refine therapeutic strategies.

## Acknowledgements

We are grateful to Dr. Edward Owusu-Ansah, Charles Glabe, and Peter Davies for antibodies; the Vienna *Drosophila* RNAi Center, FlyORF, and the Bloomington *Drosophila* Stock Center for fly stocks; Stanford PAN facility for primer synthesis; The Axelrod, Bogyo, Lipsick, Cobos, and Svensson labs in the Department of Pathology, and the Kebebew lab in the Department of Surgery, Stanford University School of Medicine, for sharing reagents and equipment; Dr. Laurel Kinman for advice on AlphaFold; J. Gaunce for maintaining fly stocks and providing technical supports, and members of the Lu lab for discussions. This work was supported by the NIH (R21AG083863, R01NS084412, R01AG089752, and R37NS083417 to B.L. and R01NS120219 to S.G.). The UCSF Neurodegenerative Disease Brain Bank receives funding support from NIH grants P30AG062422, P01AG019724, U01AG057195, and U19AG063911, as well as the Rainwater Charitable Foundation and the Bluefield Project to Cure FTD. Research organization/authors hereby express its thanks for the cooperation of Donor Network West and all of the organ and tissue donors and their families, for giving the gift of life and the gift of knowledge, by their generous donation.

## Author contributions

W.L and S.R designed study, performed experiments, analyzed data, wrote the manuscript, and contributed equally to the study; S.B, L.Y, and B.G.L performed experiments and analyzed data; L.T.G, S.S, M.I.C.S, W.W.S provided key reagents; S.G provided key reagents and advice, wrote the manuscript, and provided funding; B.L conceived and supervised the study, performed experiments, wrote the manuscript, and provided funding.

## Methods

### Mouse genetics

All procedures followed animal care and biosafety protocols approved by the Stanford University Administrative Panel on Animal Care (APLAC). The following mouse strains from Jackson Laboratory were used in this study: C57BL6 (Cat#JAX:000664), rTg4510 (Cat#JAX:015815), CaMKII-tTA (Cat#JAX:007004), and tau KO (Cat#JAX:004779).

For the tau-P301L transgene, genomic DNA isolated from tail snips was used for genotyping by PCR with the following primers: forward primer, 5’- GATCTTAGCAACGTCCAGTCC -3’, reverse primer, 5’- TGCCTAATGAGCCACACTTG -3’. For CaMKII-tTA transgenic mice, the PCR primers used for genotyping were forward primer: 5’- GACCTGGATGCTGACGAAGG -3’; reverse primer: 5’- GCAGCTCTAATGCGCT GT -3’. The tau-P301L mice were crossed with CaMKII-tTA mice to generate the rTg4510 bi-transgenic model with conditional (inducible/reversible) tau-P301L expression, and the tau P301L, CaMKII-tTA, and noncarrier littermates that served as controls. Homozygous tau KO mice were self-crossed to retain the homozygotes colonies.

### Cell lines

HEK293T cells (Cat# CRC-1573, ATCC) were cultured in DMEM high-glucose media supplemented with 10% FBS. All iPSCs were maintained in mTeSR medium. The following iPSC lines were used: Healthy control (Cat# AG28262, Coriell Medical Institute), APP duplication (UCSD239i-APP2-1, WiCell), tau-P301L and isogenic control (ND50098 and ND50046, NINDS Human Cell Repository). iPSC-induced neuronal culture is described in the following sections.

### Human brain tissues

Frozen cortex tissues from healthy and AD patients were obtained from The Memory and Aging Center and Neurodegenerative Disease Brain Bank, University of California San Francisco, and frozen pons tissues from healthy and PSP patients were obtained from the Department of Pathology Brain Bank, Stanford University School of Medicine. The primary neuropathologic diagnosis confirming AD and PSP patient status, PMI, sex, and age at death are summarized in Extended Data Table S1. Informed consent to undergo autopsy was provided by patients and/or their surrogates, following the principles outlined in the Declaration of Helsinki.

### Lentivirus production, primary mouse glia cell culture, and human iPSC neuron generation

The plasmid Tet-Ngn2-puro (Addgene #52047) or FUW-M2rtTA (Addgene #20342) (20 µg) and three helper plasmids (pRSV–REV, pMDLg/pRRE, and pMD2.G, Addgene#12253, 12251, and 12259, respectively; 10 µg each) were co-transfected with polyethylenimine into 80%–90% confluent HEK cells plated on polyornithine-coated 10 cm dishes. 6-8h after transfection, the HEK cell culture medium was completely replaced with fresh media and viral supernatant harvested at 24h and 48h. Lenti-X concentrator (Cat#631231, Takara) was used to concentrate the viral particles at 4°C for 24h. Then the mixture was centrifuged for 45 min at 1500 x g and the pellet reconstituted in 1/100th of the original volume using DMEM, snap frozen and stored at -80°C.

For making mouse glial cell culture, two postnatal day 0-3 C57BL/6J pups were anesthetized on ice. Heads were removed from pups and the cortices dissected from the brain and put into cold HBSS. HBSS was removed and 5 ml dissociation solution (5 ml HBSS, 80 µl Papain (Cat# LS003127, Worthington Biochemical), 5 µl EDTA (0.5 µM), 5 µl CaCl_2_ (1 µM) was added. After incubation for 20 min in 37°C incubator, the dissociation solution was removed with caution and the tissue triturated to single cells with a pipette in DMEM high-glucose media supplemented with 10% FBS and cells were plated on a 10cm dish. The media was changed on the next day, and cells were used after two passages.

For iPSC neuronal culture, iPSCs were dissociated with accutase and plated on Matrigel-coated plates. Cells were infected with lentiviruses containing expression constructs of rtTA and Ngn2, in the presence of Rock inhibitor. The next day, media was replaced with N2 media (DMEM/F12, N2, and insulin 10 ng/ml supplemented with doxycycline 2 µg/ml). 24 hours later, start the puromycin selection for 2 days. Then media was replaced with B27 media (Neurobasal-A, L-glutamine, B27, doxycycline 2 µg/ml, NT3 10 ng/ml, laminin 20 ng/ml and BDNF 10 ng/ml). For capturing neuronal Synapsin signals, neurons were dissociated using accutase and seeded on mouse glial culture and additionally supplemented with 5% FBS in B27 media to support glia cell survival. Media was half-exchanged every 3-4 days. Matured neurons were used after 4 weeks.

The following reagents were used in iPSC neuronal culture: Accutase (Cat#07920, Stem Cell Technologies); Ara-C (Cat#C6645, Sigma-Aldrich); B-27 supplement (Cat#12587010, Thermo Fisher); BDNF (Cat#450-02, PeproTech); BS3 (Cat#S5799, Sigma-Aldrich); Rock inhibitor (Axon Medchem, Cat#1683; 10 µM); DMEM/F-12 media (Cat#11320082, Thermo Fisher); DMEM/F-12 with 15 mM HEPES (Cat#36254, Stem Cell Technologies); ECDM (Cat#165344-1G, Millipore); Insulin (Cat# I0516, Sigma-Aldrich); L-Glutamine (Cat#25030164, Thermo Fisher); Matrigel (Cat#356230, BD Bioscience); mTeSR1 media (Cat#85850, Stem Cell Technologies); N-2 supplement (Cat#17502048, Thermo Fisher); Neurobasal-A Medium (Cat#10888022, Thermo Fisher); NT-3 (Cat#450-03, PeproTech); Laminin/Poly-D-lysine Coating Solution (Cat#LPDL001, Sigma-Aldrich); Polybrene (Cat#H9268, Sigma-Aldrich); Polyethylenimine (PEI) (Cat#23966, Polysciences Inc); Puromycin (Cat#P8833, Sigma-Aldrich); Fetal Bovine Serum (FBS) (Cat#S11550, ATLANTA Biological); Recombinant mouse laminin (Cat#23017-015, Invitrogen).

### Neuronal viability assay

After the puromycin selection, neurons were further dissociated into single cells using accutase and cultured in Laminin/Poly-D-lysine coated black 96-well plates (Corning, Cat#3603). Four weeks later, neurons were treated with different concentrations of chemicals for 24h. Then cell viability was measured with the Alamar Blue Cell viability reagent (Cat#DAL1025, Invitrogen) at 1:10 dilution, after 1h incubation at 37°C according to manufacturer’s instructions.

### Treatment of neurons with chemicals and viruses

Neurons derived from tau-P301L with isogenic wildtype control iPSCs, and APP duplication with age-, gender-, and ethnicity-matched normal control iPSCs were treated with the indicated concentrations of FK866 (Cat#S2799, Sellekchem), H_2_O_2_ (Cat#H1009, Sigma-Aldrich), DES (Cat#112402, Sigma-Aldrich), MitoTEMPO (Cat#SML0737, Sigma-Aldrich; 10 µM), CPT (2.5 µM), NMN (Sigma, Cat#N3501, 20 µM) for 24h; PP2A activator sodium selenite (Cat#S5262, Sigma-Aldrich) 10 µM for 6h; VDAC inhibitor DIDS (Sigma, Cat#309795, 50 µM), Hsp70 inhibitor PES-CI (MedChemExpress, Cat#HY-15484, 20µM), Hsp90 inhibitor geldanamycin (Selleckchem, Cat#S2713, 1 µM) for 6h. Lenti-sh-Tau (Cat#sc-36614-V, Santa Cruz Biotechnology), lenti-Tau-WT^89^, and lenti-Tau-S2A^89^ were transfected to control neurons in the presence of polybrene (8 µg/ml) and media changed after 24h. Lenti-EGFP^89^, lenti-MKI-EGFP^89^, lenti-shNC^100^, and lenti-shNDUFS3 (#31663171, Abmgood), lenti-shVDAC1, lenti-shHSP70, lenti-shHSP90 (Genscript) were transfected into APP neurons in the presence of polybrene (8 µg/ml) and media changed after 24h. Lenti-P301L-tau, lenti-0N3R-tau, lenti-0N4R-tau, lenti-2N3R-tau, and lenti-2N4R-tau viruses were generated by Genscript and used to transfect control iPSC neurons.

### Chemical treatment and transfection of cultured cells

HEK293T cells were treated with the VDAC inhibitor DIDS (50 µM, Sigma, Cat#309795), Hsp70 inhibitor pifithrin-α hydrobromide (20µM, MedChemExpress, Cat#HY-15484), Hsp90 inhibitor Geldanamycin (1 µM, SelleckChem, Cat#S2713) for 6h. To perform RNAi, HEK293T cells were transfected with Invitrogen Silencer siRNAs for VDAC1 (ID: 4728), HSP70 siRNA (ID: 16306), or HSP90 siRNA (ID: 122318) at 20 nM using Lipofectamine™ RNAiMAX (Invitrogen, Cat#13778030) for 48h prior to subsequent analyses. To overexpress WT and mutant NDUFS3 in HEK293 cells, WT and mutant (R43A, R49A, R43A/R49A, V72P, Q73P, Q74P, V75P, Q76P, or V77P) constructs of mouse NDUFS3 synthesized and cloned into the pcDNA3 vector by Genscript were used.

### Drosophila genetics

We obtained the following fly line from the Bloomington *Drosophila* Stock Center: *elav-GAL4* (8765), *UAS-ND30 (NDUFS3)* RNAi (44535), *tau KO* (64782), *nSyb-GAL4* (51635), *esg-Gal4* (93857), and *UAS-mito-roGFP2-ORP1* (67667). We received the following fly line from Vienna *Drosophila* Stock Center: *UAS-tau* RNAi (101386), *UAS-ATG1* RNAi (16133), *UAS-Sirt2* RNAi (103790). The sources of the other fly stocks are as follows: Dr. T. Littleton (*Mhc-GAL4*), Dr. J. Knoblich (*UAS-SoNar*), Dr. Mel Feany (UAS-tau-R406W), Dr. S. Birman (*TH-GAL4*), Dr. H Jacobs (*UAS-AOX*), Dr. Stephan L Helfand (*UAS-Sirt4*), and Dr. William Saxton (*UAS-mito-GFP*). All other fly lines are lab stocks maintained in the Lu lab. The indicated UAS RNAi and OE fly lines were crossed to *Mhc*-Gal4, *elav*-Gal4, or *esg*-Gal4 driver lines for muscle, pan-neuronal, or gut expression, respectively. Fly culture and crosses were performed according to standard procedures. Adult flies were generally raised at 25°C and with 12/12 hr dark/light cycles. For heat stress assay, flies were raised at 32°C. Fly food was prepared with a standard receipt (Water, 17 L; Agar, 93 g; Cornmeal, 1,716 g; Brewer’s yeast extract, 310 g; Sucrose, 517 g; Dextrose, 1033 g).

### Drug treatments in *Drosophila*

Adult flies were treated with 10 µM CPT or DES (Sigma #112402, 10 mM) for the indicated times. Newly emerged flies were treated with VDAC inhibitor DIDS (MedChemExpress #HY-D0086, 100 µM), Hsp70 inhibitor PES-CI (Sigma-Aldrich #5310670001, 60 µM), or Hsp90 inhibitor Geldanamycin (MedChemExpress #HY-15230, 10 µM) for 2-3 weeks. For the GeneSwitch flies, RU486 (Cayman Chemical #10006317, 100 µg/ml) was added to the food to induce tau expression in newly eclosed flies. Drug-containing fly food was prepared by adding drugs to microwave-melted fly food kept at a warm temperature (∼ 50°C). After thorough mixing with the chemicals, the fly food was aliquoted into plastic vials in small volume, left at room temperature to harden, and then stored at 4°C. Flies were changed to fresh drug-containing food every 2-3 days.

### Aversive taste memory assay in *Drosophila*

We performed the taste memory assay as described previously^81^. Briefly, flies were starved for 12-18h in an empty vial on wet Kimwipe paper. Flies were anesthetized on ice and fixed on a glass slide by applying nail polish to their wings. 10-15 flies were used for each experiment. Flies were incubated in a humid chamber for 2h to allow recovery from the procedure. In the pretest phase, flies were presented with 500 mM sucrose stimuli (attractive tastant) to their legs using Kimwipe wick. Flies that showed positive proboscis extension to the stimulus were used for the next phases. In the training phase, the flies were presented with 500 mM sucrose stimuli to their legs while being simultaneously punished with 50 mM caffeine (aversive tastant) applied to their extended proboscis. Training was repeated 15 times for each fly. The last phase was the test phase where the flies were given 500 mM sucrose to their legs at different time intervals (0, 5, 15, 30, 45, and 60 min), and proboscis extension was recorded. Each experiment was carried out ≥ 4 times.

### Climbing activity and wing posture assays in *Drosophila*

Climbing assay was performed as described previously^43^. Briefly, around 10-20 flies were transferred to a clean plastic vial. The flies were allowed to get accustomed to the new environment for 3-4 min and subsequently measured for the distance of bang-induced vertical climbing (negative geotaxis response). The performance was scored as percentage of flies crossing the 8 cm mark within 12 seconds. Each experiment was performed ≥ 4 times.

### *Drosophila* lifespan analysis

Flies were reared in plastic vials containing cornmeal medium. Flies were anesthetized using CO_2_ and collected at a density of 20 flies/vial. All flies were kept at humidified incubators at 12h on/off light cycle at 25°C. Flies were flipped into fresh vials every 3 days and the number of dead animals was recorded. Each set of experiment was carried out ≥ 4 times.

### Heat stress exposure in mice and flies

Two-month-old C57BL/6 WT and tau KO mice were transferred to a chamber maintained at 43°C and 60 ± 10 % humidity for 15 min once a day for 14 days. Mice were started for the battery of behavioral tests following the last heat exposure.

Newly eclosed flies were transferred to an incubator maintained at 32°C and 60 % humidity. All the behavioral and molecular assays were performed after two weeks of incubation.

### Mitochondrial purification

Intact mitochondria from *in vitro* cultured human cells, animal and human tissues were purified and quality controlled for the absence of contamination by other organelles according to established procedures. For analysis of fly samples, either brain or thoracic muscle were dissected. Samples were homogenized using a Dounce homogenizer. After two steps of centrifugation (1,500 g for 5 minutes and 13,000 g for 17 minutes), the mitochondria pellet was resuspended and washed twice with HBS buffer (5 mM HEPES, 70 mM Sucrose, 210 mM Mannitol, 1 mM EGTA, 1x protease inhibitor cocktail), then resuspended and loaded onto Percoll gradients. After centrifugation (16,700 rpm, 15 minutes, Beckman SW-40Ti rotor), the fraction between the 22% and 50% Percoll gradients containing intact mitochondria was carefully transferred into a new reaction tube, mixed with 2 volumes of HBS buffer, and pelleted by centrifuging at 20,000g for 20 minutes at 4°C to collect the mitochondrial samples for further analyses.

For PK treatment, equal volume of purified mitochondria was resuspended and digested with different concentrations of PK or PK along with 1% Triton X-100. PK treatment was performed on ice for 30 min. The reaction was blocked by adding SDS buffer before, followed by SDS PAGE and western blot for further analysis.

Mitochondrial sub-fractionation was performed using mouse brain mitochondria. Briefly, the brain tissue was homogenized in buffer A (300 mM Sucrose, 0.1 mM EGTA, 10 mM HEPES, pH 7.4) with a Dounce homogenizer. The homogenate was then centrifuged at 600g for 5 min. The supernatant was collected and filtered through cheesecloth and centrifuged at 3,300g for 10 min. The pellet was suspended in buffer B (10 mM KH_2_PO_4_, pH 7.4) and the mixture was incubated at 4°C for 15 min. Equal volume of buffer C (32% Sucrose (w/v), 30% Glycerol (v/v), 10 mM MgCl_2_, 10 mM KH_2_PO_4_, pH 7.4) was added to the mixture and incubated for another 15 min. The mixture was then centrifuged at 12,000g for 10 min and supernatant (S1) and pellet (P1) were collected separately. The pellet (P1) was resuspended in 3 ml of buffer B and mixed for 30 min at 4°C by gentle shaking, and subjected to ultracentrifugation at 160,000g for 30 min. The supernatant (S2), containing mitochondrial matrix, and pellet (P2) containing inner membrane were collected. S1 was also subjected to ultracentrifugation at 160,000g for 30 min, and the resulting supernatant (S3), containing inter membrane space (IMS), and pellet (P3) containing outer membrane (OM) were collected. The protein samples were further processed for SDS PAGE and western blot analysis.

### NAD^+^/NADH measurements

NAD^+^/NADH was measured essentially as described, using an NAD^+^/NADH quantification colorimetric kit according to the manufacturer’s instructions (AAT Bioquest #15273). Briefly, cell pellets or tissue samples were lysed using the lysis buffer for 15 min at 37◦C and lysates were collected after centrifugation at 12,000g for 15 min. For the measurement of NAD^+^ amount, NAD^+^ extraction solution was added into the lysates and incubated at 37◦C for 15 minutes. Thereafter neutralization solution was added to neutralize the NAD^+^ extracts. Absorbance was monitored at 460 nm after adding NAD/NADH working solution and 1hr incubation at room temperature with protection from light. To measure total NAD^+^ and NADH amount - NAD(H) (total), control extraction solution was added into the lysates and incubated at 37◦C for 15 minutes. Thereafter again control extraction solution was added. Absorbance was monitored at 460 nm after adding NAD/NADH working solution and 1hr incubation at room temperature with protection from light. The ratio of NAD^+^ /NADH was determined by the following equation: ratio = NAD^+^/ NAD(H) (total) – NAD^+^. Each experiment was performed ≥ 4 times.

NAD^+^/NADH was also assayed with another commercial kit (WST-8) by following the manufacturer’s protocol (MedChemExpress #HY-K0313). Briefly, cell or tissue samples were homogenized with the NAD^+^/NADH extraction solution on ice and lysates were collected after centrifugation at 12,000 rpm for 10 minutes. Total NAD(H) was measured by mixing 20 µl of test samples with 90 µl of ADH working solution followed by 10 minutes of incubation at 37°C. Then 10 µl of chromogenic liquid was added to the mixture and measured at 450nm after 30 minutes of incubation at 37°C. To measure NADH, 50-100 µl samples were heated at 60°C for 30 minutes to decompose NAD^+^. Then 20 µl of sample was mixed with 90 µl of ADH and incubated for 10 minutes at 37°C. After 10 µl of chromogenic liquid was added to the mixture, the samples were incubated for 30 minutes and detected at 450nm. The NAD^+^/NADH ratio was calculated as: NAD^+^/NADH = Total NAD(H) - NADH/NADH.

NAD^+^/NADH measurement by mass spectrometry was performed at the Quantitative Metabolite Analysis Center (QMAC), Benioff Center for Microbiome Medicine, UC San Francisco. Lyophilized mitochondrial samples (60 μl each) were resuspended in 1 mL of extraction buffer (acetonitrile : methanol : water (2:2:1)) containing the internal standard, 2-amino-3-bromo-5-methylbenzoic acid (Sigma-Aldrich, cat. #631531). The mouse brain samples were homogenized using Precellys 24-bead homogenizer pre-cooled with dry ice at 6400rpm, 3 x 20s with 30s rest.

All samples remained on ice for 30 min after homogenization. Precipitates and other cellular debris were pelleted after centrifugation at 21,000 rpm for 10 min. Supernatants were transferred to separate Eppendorf tubes and dried under vacuum. Samples were kept in -80° C until the day of data acquisition. Samples were reconstituted in 200 μl of water, vortexed briefly, and filtered using a 0.2 mM plate filter (3,400 rpm for 10 min). Purified mitochondria and mouse brain samples were diluted respectively 2- and 10-fold and immediately analyzed.

Targeted Metabolomics For NAD^+^ and NADH: A 5 μl aliquot of each sample was subjected to targeted metabolomic analysis on a Shimadzu 30-AD UPLC in series with a SCIEX 7500 Triple Quad Mass Spectrometer. Analytes were chromatographically separated using a Kinetex F5 2.6 μm, 2.1 x 150 mm LC column (Phenomenex, cat #00F-4723-AN) with a mobile phase scheme of [A] water + 0.1% formic acid and [B] acetonitrile + 0.1% formic acid. The LC method was set to a constant flow rate of 200 μl/min at 30o C, and the timed linear gradient program consisted of: time=0 min, 0.2% B, time=2 min, 0.2% B, time=5 min, 2% B, time=11 min, 25% B, time=13 min, 98% B, time=16 min, 98% B, time=16.1 min, 0.2% B, time=20 min, 0.2% B. Data was collected in MRM mode with positive and negative polarity switching using the following source parameters: CUR = 40, GS1 =45, GS2 = 70, CAD gas = 9, Temperature =375°C, ISVF = 2000V (positive) and -2000V (negative). Optimized MRM pairs were determined from standards and consisted of a quantifier and 2 qualifiers for each analyte: NAD^+^ 664.0 / 428.0 m/z (quantifier) with CE =38, 664.0 / 524.0 m/z (qualifier) with CE =20, and 666.0 / 136.0 m/z (qualifier) with CE =43; NADH 666.0 / 649.0 m/z (quantifier) with CE =23, 664.0 / 514.0 m/z (qualifier) with CE =32, and 666.0 / 348.0 m/z (qualifier) with CE =30; 2-amino-3-bromo-5-methylbenzoic acid 231.0 / 104.0 with CE =35. In negative mode, additional MRM pairs were monitored for 2-amino-3-bromo-5-methylbenzoic acid 229.9 / 81.0 with CE = -35 and 227.9 / 79.0 with CE = -35. Other method parameters include a cycle time of 0.165 seconds, dwell time of 10 ms/mrm pair, settling time = 15 ms, and pause time of 5.007ms. Raw data was processed using a built-in SCIEX OS software package (version 2.1.6.59781) for peak picking, integration, and determination of calculated concentration.

We used a genetically encoded NAD^+^/NADH sensor SoNar to measure NAD^+^/NADH by live imaging in *Drosophila*. Briefly, the *UAS-Gal4* system was used to express the *UAS-SoNar* transgene^56^ in neurons using the pan-neuronal *elav*-Gal4 driver. Flies with or without drug treatment were dissected at the indicated ages, and brain samples were kept in Schneider’s media and immediately visualized on a Leica SP8 confocal microscope with 405 nm or 488 nm laser to detect NADH or NAD^+^ signals, respectively. Since the SoNar-NADH signal is sensitive to photo bleaching, we first detected NADH signal using 405nm laser. The signal intensity was measured using the NIH ImageJ software.

### ROS measurement

The Amplex Red Hydrogen Peroxide/Peroxidase Assay Kit (Cat#A22188, Thermo Fisher) was used for the measurement of ROS according to the manufacturer’s instructions. Briefly, for whole lysates, cell pellets were gently homogenized in PBS with a loose-fitting homogenizer, and the homogenates were added to 96 well plate. Then the working solution of 100 µM Amplex Red reagent/0.2 U/ml HRP were added and incubated for 30 min at room temperature. Absorbance was monitored at 560 nm, and the protein concentration of the samples was determined to normalize ROS levels. For fly brain tissues, mouse cortex tissues, and human brain tissue samples, tissues were homogenized in PBS with a loose-fitting homogenizer, and homogenates were centrifuged at 2,000 rpm for 5 minutes. The supernatant containing intact cells was used for ROS determination as described above.

For ROS measurement by CM-H2DCFDA staining, CM-H2DCFDA (Cat#C6827, Invitrogen; 5 µM) was added to live neurons or freshly dissected brain tissues of flies and allowed to incubate for 30 min. Tissue samples were immediately imaged on a Leica SP8 confocal microscope with an excitation laser at 492–495 nm and detection at 517–527 nm using 20x or 40x oil-objective lens.

*In vivo* live imaging of ROS was performed in *Drosophila* using the genetically-encoded mito-roGFP-ORP1 reporter. Briefly, flies with the mitochondria-specific ROS reporter *UAS-mito-roGFP-ORP1* expressed in the brain under *elav-Gal4* control were dissected in insect Schnider medium and imaged under Excitation at 488 (reduced) or 405 (oxidized) nm and with Emission at 510 nm. Average intensity of individual brain was quantified using image J.

### *In vitro* mitochondrial RET and FET and tau import assays

Isolated mitochondria were incubated in an assay medium (125 mM KCl, 20 mM HEPES, 2 mM K_2_HPO_4_, 1 mM MgCl_2_, 0.1 mM EGTA, 0.025% BSA, pH7.0). To induce FET, 2.5 mM Malate and 2.5 mM Glutamate were supplemented as substrates. To induce RET, 5 mM Succinate and 1ug/ml Oligomycin were supplemented into the mitochondrial samples. To more closely mimic *in vivo* conditions where mitochondria are likely to be in state 3 or nearly state 3 respiration in the presence of ADP, FET or RET derived ROS and NAD^+^/NADH ratio change were also examined in the presence of ADP. For ROS detection, 1 uM Amplex Red and 5U/ml horseradish peroxidase were added into the reaction media. Finally, fluorescence was recorded by excitation at 560 nm and emission at 590 nm.

Treatment of purified mitochondria with recombinant tau protein was performed under conditions conducive to protein import as previously described^101^. Briefly, mitochondria isolated from 6-month-old C57BL6 mouse muscle tissues, HEK293T cells or hiPSC neurons after treatment with VDAC, Hsp70, or Hsp90 chemical inhibitors or RNAi-mediated knockdown of VDAC1, Hsp70, or Hsp90, or *Drosophila* fed with vehicle or VDAC, Hsp70, or Hsp90 chemical inhibitors, using HBS buffer in the presence of proteinase inhibitors. Freshly prepared mitochondria were incubated with recombinant human tau (2N4R) pretreated with GSK-3β to induce tau phosphorylation (abcam Cat#269022) or with recombinant phospho-mutant tau (tau-S404A) (abcam Cat#269016) in an import buffer (125 mM KCl, 20 mM HEPES, 2 mM K_2_HPO_4_, 1 mM MgCl_2_, 0.1 mM EGTA, pH7.0). Treatment was performed in the import buffer for 30 min at 37°C. The presence of tau/p-tau inside mitochondria was examined by western blot after PK treatment to remove peripherally associated tau. RET and FET assays were performed as described above after 30 min of incubation of recombinant tau with the mitochondria.

### Mitochondrial Respirometry

Mitochondria were isolated and purified as described above. Purified mitochondria were seeded into 24-well cell culture microplates (Seahorse Bioscience, Agilent), and centrifuged at 2,254 g at 4°C for 20 min. The assay was performed in mitochondria assay medium (115 mM KCl, 10 mM KH_2_PO_4_, 2 mM MgCl_2_, 3 mM HEPES, 1mM EGTA, FA-free BSA 0.2% pH 7.2), along with complex I assay medium (11 mM Pyruvate, 11 mM Malate, 11 mM L-Proline pH 7.2.). The hydration of a Seahorse XF96 sensor cartridge was performed one day prior to the assay. The loading of compounds (ADP, Oligomycin, FCCP, Antimycin A & Rotenone) and calibration of the cartridge sensor were performed on the day of the assay. Following instrument calibrations, cell culture plates containing mitochondria were transferred to the XFe24 analyzer (Seahorse Bioscience, Agilent) to record the OCR at the appropriate temperature.

### C-I activity assay

Complex I activity assay was performed with the Mitocheck complex I assay kit from Cayman Chemicals (Mitocheck, CI activity assay kit, #700930), following the manufacturer’s protocol.

### Succinate assay

Mitochondria succinate level was examined with the Succinate Assay kit from abcam (cat#ab204718), following the manufacturer’s protocol.

### RT-PCR

A total of 100 ng RNA was reverse transcribed into cDNA using the iScript cDNA synthesis kits (Cat#1708890, Bio-Rad). Template cDNA was amplified using PowerUp SYBR Green Master Mix (Cat#A25741, Biosystems) with the following cycling parameters: Activation 95°C, 10 min; Denature 95°C, 15 s, Annealing/extension 60°C, 1 min for 40 cycles. Quantitative RT-PCR was carried out on the Applied Biosystems Step One Real-Time PCR System (Cat#4376600, Applied Biosystems). Relative quantity values were calculated using the 2^−ΔΔCt^ method. β-actin was used to normalize the expression levels of each sample. The following gene-specific primers were used:

Cst7 Forward: GCCGAACTACATGCAGGAAG

Cst7 Reverse: AGTCACTGGCAGAGGAGAAC

Axl Forward: TGCACAAGATCAGAGCTGGA

Axl Reverse: CGGCCATGAACTTCACTAGC

Lpl Forward: TTCAACCACAGCAGCAAGAC

Lpl Reverse: CTGGATAATGTTGCTGGGCC

Itgax Forward: TGTGATGAGCCAGCTTCAGA

Itgax Reverse: TGGGTGGTGAACAGTTCTGT

Clec7a Forward: ACAGTACACCAGACACAGGG

Clec7a Reverse: TGGCCAGACAGCATAAGGAA

Cox6a2 Forward: AGACAGAGAAGGACAGTGCC

Cox6a2 Reverse: AGCAGTTAAGGGAGCAGAGG

Ank Forward: TCTGTGTCATTCCTCTGCGT

Ank Reverse: CTCCCACAAACCCTGCTAGA

Csf1 Forward: GGGCCTCCTGTTCTACAAGT

Csf1 Reverse: AGGAGAGGGTAGTGGTGGAT

NF-κB Forward: CACCGGATTGAAGAGAAGCG

NF-κB Reverse: AAGTTGATGGTGCTGAGGGA

TNF alpha Forward: CCCTCACACTCAGATCATCTTCT

TNF alpha Reverse: GCTACGACGTGGGCTACAG

NLRP3 Forward: ATTACCCGCCCGAGAAAGG

NLRP3 Reverse: TCGCAGCAAAGATCCACACAG

3R and 4R tau Forward: AAGTCGCCGTCTTCCGCCAAG

3R and 4R tau Reverse: GTCCAGGGACCCAATCTTCGA

### Immunofluorescence staining

Immunostaining of adult fly muscle was performed as previously described^102^. Briefly, fly thoraxes were dissected and fixed with 4% paraformaldehyde (Electron Microscopy Sciences, Cat#15710) in phosphate buffered saline and 0.3% Triton X-100 (PBS-T). The tissues were washed three times with PBS-T. Samples were incubated for 30 min at room temperature in blocking buffer: 0.5% goat serum in PBS-T. The following primary antibodies were added, and samples were incubated overnight at 4°C: anti-Ubiquitin (Abcam ab140601, 1:1000); anti-P62 (Abcam ab178440, 1:1000). The samples were washed three times with PBS-T and subsequently incubated with the following secondary antibodies for 4h at 4 °C: Alexa Flour 488 (Invitrogen A32723, 1:200), Alexa Flour 594 (Invitrogen A11036, 1:200). The tissues were washed three times with PBS-T and mounted in slow fade gold buffer (Invitrogen S36936).

Immunostaining of adult brains was performed as previously described^43^. Briefly, brain tissues of adult flies were dissected and fixed on ice for 30-45 min in fixing buffer (940 µl of 1% PBS-T and 60 µl of 37% formaldehyde). Tissues were washed three times in 0.1% PBS-T and blocked overnight at 4°C in blocking buffer (1 ml 1xPBS, 0.1% Triton-X, 5 mg/ml BSA). This was followed by incubating for 16h at 4°C with the following primary antibodies: anti-TH (Pel-Freez P40101-150, 1:1000); anti-Ubiquitin (Abcam ab140601, 1:1000); anti-p62 (Abcam ab178440, 1:1000). Tissues were washed three times with 0.1% PBS-T and subsequently incubated with the following secondary antibodies for 4h at 4°C: Alexa Flour 488 (Invitrogen A32723, 1:200), Alexa Flour 594 (Invitrogen A11036, 1:200). The tissues were washed three times with PBS-T and mounted in SlowFade Gold Antifade Mountant (Invitrogen S36936) and viewed using a Leica SP8 confocal microscope.

Neurons grown in 96-well plates were washed with PBS and fixed with 4% paraformaldehyde (Cat#P6148, Sigma-Aldrich) for 10 min at room temperature. Cells were permeabilized with 0.1% Triton X-100 (Cat#X100, Sigma-Aldrich) and blocked with 10% normal goat serum (Cat# PI31873, Fisher Scientific) in PBS for 1h. Then neurons were incubated with primary antibodies overnight at 4°C, washed three-times with PBS, and incubated with secondary antibodies for 1h at room temperature. Cells were washed with PBS and incubated with Hoechst 33342 (Cat# 62249, Thermo Fisher) for 5min at room temperature, images were captured directly in 96-well plates by using a Leica DMi8 confocal microscope (https://www.leica-microsystems.com/products/light-microscopes/p/leica-dmi8-id/).

Mouse brains were immediately dissected after euthanasia and wash with cold PBS. Tissues were then fixed with 4% Paraformaldehyde (PFA) for 48h at 4°C and immersed in 30% Sucrose solution before embedding in tissue freezing medium. Frozen tissues including human samples were sectioned on a Leica CM1950 cryostat at a thickness of 20 µm. The sections were used for immunostaining, and all fluorescent images were acquired on a Leica DMi8 confocal microscope. The branch length of Iba1-immunostained microglia was analyzed by Fiji neuron J plugin (https://imagej.nih.gov/ij/). For ThioS staining, tissue sections were incubated with a Thio S solution (Cat#T1892, Sigma-Aldrich; 0.05% in distilled water) for 5 minutes after secondary antibody staining. Then the tissues were washed with 75% ethanol and followed by washing with distilled water before image capture.

Antibodies used were: mouse anti-tau PHF1 (Gift of Peter Davies, 1:500); mouse anti-tau CP13 (Gift of Peter Davies, 1:500); rabbit anti-NDUFV1 (Proteintech, 11238-1-AP, 1:500); p-S262 anti-tau antibody (Invitrogen, 44-750G, 1:500); rabbit anti-VDAC (Proteintech, 55259-1-AP, 1:500); rabbit anti-Iba1 (Novus Biologicals, NBP2-19019, 1:500); rat anti-CD8 (Invitrogen, MA1-145, 1:500); rabbit anti-GFAP (Proteintech, 16825-1-AP, 1:500); rabbit anti-NeuN (Proteintech, 26975-1-AP, 1:500); rabbit anti-Caspase-3 (Cell Signaling Technology, 9661T, 1:400); rabbit anti-Synapsin (Cell Signaling Technology, 5297T, 1:200); mouse anti-Tuj1 (Biolegend, 801201, 1:400); mouse anti-Tomm40 (Proteintech, 66658-1-lg, 1:400); rabbit anti-p62 (Sigma, P0067, 1:400); rabbit anti-γH2A (Cell Signaling Technology, 9631T, 1:200); goat anti rabbit/mouse/rat IgG (H + L) highly cross-adsorbed secondary antibodies, Alexa Fluor 488/568 (Invitrogen).

### Proximity Ligation Assay (PLA)

PLA reactions were performed in 4% PFA fixed rTg4510 mouse and human PSP brain sections. The protocol followed the manufacturer’s User Guide (Sigma-Aldrich). Briefly, sections were incubated with blocking solution for 1h at 37°C and then PHF-1 and NDUFV1 or PHF-1 and NDUFS3 primary antibodies were applied to sections overnight at 4°C. PHF-1, NDUFS3, or NDUFV1 primary antibody only was added as negative controls. The slides were washed, and Duolink PLA anti-Rabbit PLUS and anti-Mouse MINUS probes (Cat#DUO92002 and Cat#DUO92004, Sigma-Aldrich) were added for 1h at 37°C. The slides were washed, and the Duolink ligation far red buffer (Cat#DUO92013, Sigma Aldrich) was added for 30 min at 37°C. The slides were washed, and the amplification buffer was added for 100 min at 37°C. The slides were washed and mounted in mounting medium with DAPI counterstaining.

### Blue native polyacrylamide gel electrophoresis (BN-PAGE)

Mitochondria were isolated form brain tissues as described previously^102^. Isolated mitochondrial pellet was suspended in native PAGE sample buffer (Invitrogen, #BN20032) along with 1% digitonin and protease inhibitors. Following 15 min incubation on ice, samples were centrifuged at 20,000g for 30 min. The supernatant was collected and mixed with the G-250 sample additive and loaded onto 4-16% precast native PAGE gels (Invitrogen, #BN1004BOX). The NativeMark protein standard (Thermo Fisher, #LC0725) was run along with samples to estimate molecular weight. Electrophoreses was conducted using Native PAGE running buffer (Invitrogen, #BN2007) as anode and Native PAGE running buffer containing Cathode buffer additive (Invitrogen, #BN2002) as cathode. Gels were stained with Coomassie blue staining reagent to visualize protein bands. 2D BN-PAGE/SDS-PAGE was performed by first separating mitochondrial proteins by BN-PAGE on the first dimension, followed by SDS-PAGE on the 2^nd^ dimension. Briefly, BN PAGE gel strip was equilibrated with equilibrium buffer (125 mM Tris-HCl, 2% SDS, 1% DTT, and 10% glycerol) for 15-30 min with gentle agitation. The gel strip was placed on the top of the SDS-PAGE gel and proteins were separated by running the gel in SDS running buffer. The gel was further processed by Coomassie brilliant blue (CBB) staining of total proteins or western blot detection of specific proteins.

### Biochemical purification of complex I

Mitochondria complex I was purified from mice brain tissues as described previously^72^. Briefly, 100 mg mitochondrial membrane was treated with one cOmplete protease inhibitor tablet (Roche, #11836170001) and solubilized on ice at 4.5 mg/ml by the stepwise addition of lauryl maltose neopentyl glycol (LMNG, Anatrace, #NG310) from a 5% stock solution to a final concentration of 1.5%, followed by a further 20 min incubation, and centrifugation (8,500 g, 15 min, 4 °C). The supernatant was filtered through a 0.22 μm syringe-tip filter and loaded onto three 5 ml Q-Sepharose HP columns (Cytiva, #17115301) connected in series and pre-equilibrated with Buffer A (20 mM Tris-HCl (pH 7.5), 2 mM EDTA, 10% (v/v) Ethylene Glycol, 0.15% (w/v) LMNG, 0.02% Asolectin (total soy lipid extract, Avanti Polar Lipids, #541602), and 0.02% 3-[(3-cholamidopropyl)-dimethylammonio]-1-propanesulfonate (CHAPS, Sigma-Aldrich, #226947)). The columns were washed with 23% Buffer B (Buffer A + 1 M NaCl) then Buffer B was increased to 38% to elute complex I. Complex I-containing fractions were collected, pooled and concentrated using an Amicon Ultra-15 (100 MWCO) and then injected onto a Superose 6 Increase column (Cytiva, # 29091596) pre-equilibrated with Buffer C (20 mM Tris-HCl (pH 7.5), 150 mM NaCl, 10% (v/v) glycerol, 0.0375% LMNG). Complex I-containing fractions were concentrated using an Amicon Ultra-15 and flash frozen in liquid N2. The concentration of complex I was measured and the NADH:APAD+ and NADH:DQ oxidoreductase rates were determined using the complex I activity assay kit (Mitocheck, CI activity assay kit, Cayman Chemicals, #700930).

### Immunoblotting and co-immunoprecipitation

Around 5 fly thoraxes or 15 heads were homogenized in 75 µl of regular lysis buffer (50 mM Tris-HCl, 150 mM NaCl, 1% Triton X100, protease inhibitors) on ice. Samples were homogenized using a hand-held mechanical homogenizer for 30 secs. The homogenized samples were incubated on ice for 30 mins before centrifuging at 15,000 rpm for 20 mins at 4°C. 30 µl of the supernatant was mixed with 10 µl of 4x Lammaelli buffer (BioRad #161-0747) and boiled for 5 mins at 100°C. The protein lysate was cooled, centrifuged and loaded onto 4-12% Bis-Tris gel (Invitrogen, #NP0321) or 16% Tricine gel (Invitrogen, #EC66955) with 1x MES (Invitrogen, #NP0002) as running buffer as previously described^103^.

Neurons and brain tissues were lysed with RIPA buffer (Cat#89900, Thermo Fisher) supplemented with protease inhibitors (Sigma, #4693132001) and phosphatase inhibitors (Sigma, #4906845001) with sonication on ice until brain particulates became invisible. After centrifugation at 16,000g for 15 min at 4°C, the supernatant was transferred to new tubes. Protein concentration was measured using Pierce BCA protein assay kit (Thermo Fisher, #23225). Equal amounts of total protein were subjected to SDS-PAGE. After transferring the proteins to a PVDF membrane (Sigma, #IPVH00010), the membrane was blocked with 5% milk in TBS with 0.05% Tween (TBST) for 1 h at room temperature. The following primary antibodies and dilutions were used: mouse anti-tau PHF1, 1:500 (Gift of Peter Davies); mouse anti-tau CP13, 1:500 (Gift of Peter Davies); rat anti-dSDHA and rabbit anti-dNDUFV1 1:1000 (Gift from Dr. Edward Owusu-Ansah); p-S262 anti-tau antibody 1:500 (ThermoFisher #44-750G); mouse anti-dTau 1:500 (DSHB #5A6); rabbit anti-Tom20 1:1000 (abclonal #A19403); rabbit anti-cytochrome c 1:1000 (Proteintech #10993-1-AP); rabbit anti-LonP1 1:1000 (Proteintech #15440-1-AP); rabbit anti-Map2 1:1000 (Proteintech #17490-1-AP); rabbit anti-NDUFS1 1:1000 (Proteintech #12444-1-AP); rabbit anti-NDUFA10 1:1000 (Proteintech #32947-1-AP); rabbit anti-SDHA 1:1000 (Proteintech #14865-1-AP), rabbit anti-ATP5α 1:1000 (abcam #ab14748); mouse anti-tau AT100, 1:500 (Invitrogen, MN1060); mouse anti-tau HT7, 1:1000 (Invitrogen, MN1000); mouse anti-Tau7, 1:1000 (Sigma, MAB2239); rabbit anti-tau Thr231, 1:500 (Invitrogen, 44-746G); rabbit anti-NDUFV1, 1:1000 (Proteintech, 11238-1-AP); rabbit anti-NDUFS3, 1:1000 (Proteintech, 15066-1-AP); rabbit anti-GFAP, 1:500 (Proteintech, 16825-1-AP); rabbit anti-IL-1β, 1:500 (Proteintech, 16806-1-AP); rabbit anti-Cleaved Caspase-1, 1:500 (Cell Signaling Technology, 4199T); rabbit anti-ASC/TMS1, 1:500 (Cell Signaling Technology, 67824T); rabbit anti-Drp1, 1:500 (Cell Signaling Technology, 4494S); rabbit anti-Opa1, 1:500 (Cell Signaling Technology, 80471T); rabbit anti-Glu2B, 1:500 (Cell Signaling Technology, 4212T); rabbit anti-p-GSK3β, 1:500 (Cell Signaling Technology, 5558T); rabbit anti-GSK3β, 1:500 (Cell Signaling Technology, 12456S); rabbit anti-MARK4, 1:500 (Cell Signaling Technology, 4834S); rabbit anti-MFN2, 1:500 (ProSci, 7863); mouse anti-PSD95, 1:500 (BioLegend, 810302); mouse anti-PP2A, 1:500 (BD Biosciences, 610556); mouse anti-Tau46, 1:500 (Sigma, T9450); rabbit anti-p-PAR-1(T408)^90^, 1:500; rabbit anti-VDAC1, 1:500 (Proteintech, 55259-1-AP); rabbit anti-Hsp70, 1:200 (Santa Cruze, sc-25837); rabbit anti-Hsp90, 1:500 (Proteintech, 13171-1-AP). Antibodies were incubated overnight at 4°C. The corresponding HRP-conjugated secondary antibodies were detected by ECL western blot reagents (Revvity, NEL105001EA). Immunoblot bands was analyzed with Fiji software (https://imagej.nih.gov/ij/).

Co-immunoprecipitation (co-IP) assays were performed essentially following a protocol described before^43^. Briefly, cell pellets or tissue samples were homogenized in lysis buffer [50 mM Tris-HCl, pH7.4, 150 mM NaCl, 5 mM EDTA, 10% glycerol, 1% Triton X-100, 0.1mg/ml cycloheximide, 1x RNase inhibitor, and Complete protease inhibitor cocktail (cat#B14012, Bimake)]. Additional Phosphatase Inhibitor Cocktail (cat# B15001, Bimake) was applied if phosphorylation signal was to be detected. After centrifugation at 10,000 g for 5 min, the supernatant was subjected to immunoprecipitation using the indicated primary antibodies, or affinity gels at 4°C for 6 hours with gentle shaking. In general, 500 µg of total proteins was used for co-immunoprecipitation assays. Subsequently, protein A-Sepharose beads were added to the mixture and incubated at 4°C for 2 hours with gentle shaking. The Sepharose beads were washed three times (10 minutes each) at 4°C in lysis buffer, mixed with 2x SDS Sample buffer, and loaded onto SDS-PAGE gels for western blot analysis. In general, 1/5 of IP samples was loaded for western blot analysis.

### General design of mouse behavioral tests

In general, 8-month-old male and female rTg4510 and littermates, 2-month-old tau KO and C57BL/6 mice were evaluated for differences in behavior. All mice were acclimated to the testing room for one to two hours prior to testing. All behavioral equipment was cleaned with 70% ethanol between each animal testing.

### CPT treatment of wild type and diseased mouse models

Approximately 4-month-old rTg4510 mice were administered vehicle (veh) and 25 mg/kg, 50 mg/kg, or 100 mg/kg CPT by oral gavage once a day for 4 months following a 6 days/week dosing schedule. C57BL/6 wild type mice at 4-months of age were administered veh or CPT (100 mg/kg) by gavage using once a day for 2 months. Heat stress exposed C57BL/6 and tau KO mice were administered veh or 100 mg/kg CPT by oral gavage once a day for 14 days. CPT was obtained from Cerepeut Inc. under a Materials Transfer Agreement between Cerepeut Inc. and Stanford University.

### Magnetic Resonance Imaging (MRI)

The MRI studies were performed at the Neuroscience Preclinical Imaging Lab in Stanford University, on a 7T Bruker BioSpec system equipped with BGA-12S gradient insert and interfaced to ParaVision 360 V. 3.5. A 4-element receive-only mouse CryoProbeTM and 86mm volume coil were used for MR imaging. Animals were anesthetized with 3% isoflurane mixed in air and O2 and maintained with 1-1.5% isoflurane during the imaging procedure. Core body temperature was maintained at 37°C using a warm water circulator pad and a temperature controller (Thermo Scientific). Animal’s respiration was also monitored via a pneumatic pad placed under the animal (SA instruments, NY). T2w Turbo RARE sequence was used to obtain transverse structural MRI for volume measurements (TR/TE = 3300/30 ms, FOV = 20 x 20 mm^2^, matrix = 335 x 335 yielding 60 µm in-plane resolution, slice thickness = 0.4 mm, RARE Factor = 4, NEX = 3, scan time = 13 min 42 sec). MR images were quantified by using Horos software (https://horosproject.org/about/).

### Morris water maze test

Morris water maze (MWM) was an open circular pool with a diameter of 120 cm. It was filled to a depth of 0.5 m with room temperature water. The pool was divided into four equal quadrants and the environmental cues remained in the same position throughout the experiment. A circular platform (diameter, 12 cm) was placed approximately 5 mm below the water surface to make it invisible to the mice. The position of the platform remained the same during the training trials. The mice were placed into the pool from 4 different quadrants per trial for 5 consecutive days. Each mouse was given 60 s to locate the submerged platform; they were allowed to sit on the platform for 20 sec before being removed. If a mouse failed to locate the platform within 60 sec, it was gently guided to the platform and allowed to sit on the platform for 20 sec before being removed. The time to find the platform was recorded as the escape latency. The average time from the four trials was considered the daily result for each group. The probe test was performed on day 6, in which mice were allowed to swim freely for 1 min in the absence of the hidden platform. The quadrant where the platform was previously located was assigned as the “target” quadrant. The time spent in the target quadrant/ total time was calculated.

### Open field test

Mice were placed into center of an open-field arena (40 × 40 × 40 cm, W × L × H) and allowed to explore for 10 minutes. An overhead camera was used to track movement, while travel distance and velocity were recorded and analyzed with the AnyMaze software (https://www.any-maze.com/).

### Elevated plus maze

The elevated plus maze comprised of two open arms (35 × 5 cm) and two closed arms (35 × 5 × 10 cm) that extended from a central platform (5 × 5 cm) and was elevated 50 cm from the floor. Mice were tested by placing them in the center of the maze facing an open arm, and their behavior was tracked for 5 min with an overhead camera and data analyzed with AnyMaze software (https://www.any-maze.com/). Percentage of open arm time was calculated as open arm time/open arm time + closed arm time.

### Novel object test

The novel object recognition task was conducted in a testing chamber (40 × 40× 40 cm) with two different kinds of objects. Both objects were generally consistent in height and volume but are different in shape and appearance. The test ran for three days, each followed by 24 hours delay. On Day 1, mice were habituated to the testing chamber for 10 min. On Day 2, the animals were exposed to the familiar area with two identical objects placed at an equal distance for 10 min. On Day 3, the mice were allowed to explore the testing chamber in the presence of the familiar object and a novel object for 5 min to test long-term recognition memory. The discrimination index for the novel object was calculated as the [(novel object exploration time - familiar object exploration time) / novel object exploration time + familiar object exploration time.

### Fear conditioning test

This test was conducted in a sound attenuated chamber with a metal grid floor capable of delivering an electric shock, and freezing was measured with an overhead camera and Anymaze software (https://www.any-maze.com/). Mice were initially placed into the chamber and the baseline freezing time was record for 2 minutes. An 80-dB white noise served as the conditioned stimulus (CS) and was presented for 30 sec. During the final 2 sec of this noise, mice received a mild foot shock (0.5mA), which served as the unconditioned stimulus (US). After 1 minute, another CS-US pair was presented. The mouse was removed 30 sec after the second CS-US pair and returned to its home cage. Twenty-four hours later, each mouse was returned to the test chamber, and freezing behavior was recorded for 5 min (context test). Percent of time freezing was calculated as test freezing time/5 min – baseline freezing time/ 2 min.

### Computational modeling of protein structure and protein-protein interaction

The structures of mouse NDUFS3 (UniProt: Q9DCT2) and human 2N4R tau (UniProt: P10636-8) and their interaction were modelled using AlphaFold2 and AlphaFold2 multimer through ColabFold built in UCSF ChimeraX-1.9 (https://www.cgl.ucsf.edu/chimerax/). The initial modeling predicted only one interaction interface between NDUFS3 and the C-terminal sequence of tau covering S396/S400/S404. This information was used to make refined predictions in AlphaFold3 (https://alphafoldserver.com) using the C-terminal domain of tau (aa 370-441) and full-length NDUFS3 and by incorporating phosphate groups at the S396, S400, and S404 positions. This resulted in the binding poses presented in Extended Data Fig. 7. Surface charge of NDUFS3 was colored in UCSF ChimeraX (https://www.cgl.ucsf.edu/chimerax/). The charged surface residues of NDUFS3 predicted by AlphaFold3 to interact with PHF-1 p-tau were found to be solvent exposed in the published mouse complex I cryo-EM structure (PDB: 6g2j).

### Quantification and statistical analysis

Images and Western blots were analyzed using Fiji (https://imagej.nih.gov/ij/). MRI data were analyzed by using the Horos software (https://horosproject.org/about/). Statistical analysis was performed with the GraphPad Prism 10 (https://www.graphpad.com/scientific-software/prism/). Differences between two groups were analyzed using two-tailed unpaired Student’s *t* tests. Differences among more than two groups were analyzed using one-way ANOVA followed by Tukey’s multiple-comparison test. Two-way ANOVAs were used to analyze between-subjects designs with two variable factors followed by Tukey’s multiple-comparison test. Lifespan was analyzed by Kaplan Meier survival analysis. All quantitative graphs with error bars are presented as means ± SEM. *p < 0.05, **p <0.01, ***p < 0.001, ****p < 0.0001.

## Resource Availability

### Lead contact

Further information and requests for resources and reagents should be directed to and will be fulfilled by the lead contact, Bingwei Lu

### Materials availability

All unique/stable reagents generated in this study are available from the lead contact with a completed Materials Transfer Agreement.

### Data and code availability

- All data are available in the main paper, Extended Data Figures, and Supplementary Information. Source data are provided with this paper.
- This paper does not report original code.
- Any additional information required to reanalyze the data reported in this work paper is available from the lead contact upon request.

---

**Extended Data Fig. 1.**
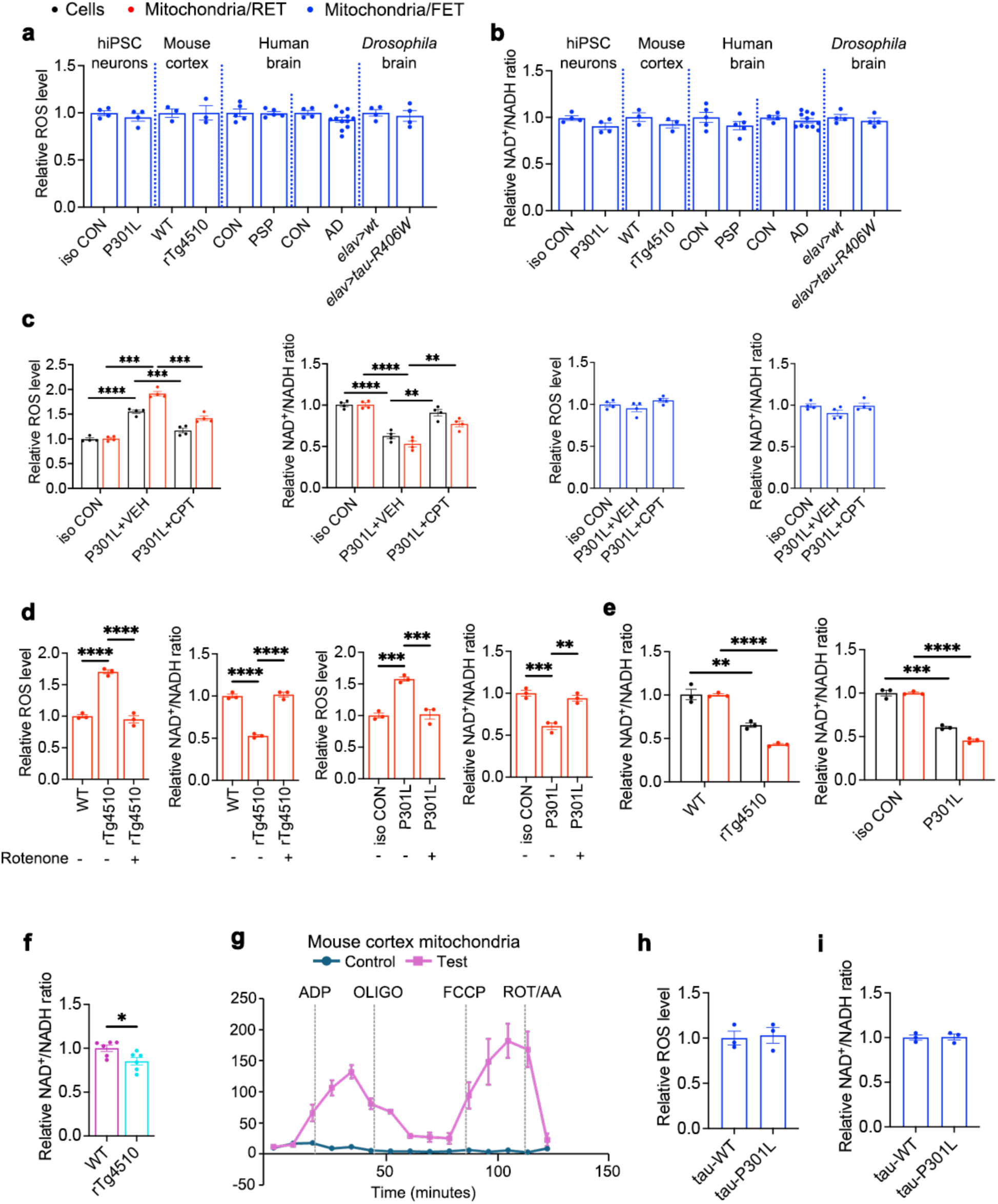
RET and FET activities in patient brain tissues and fly, mouse, and hiPSC-derived neuron models of tauopathy. **a, b**, Measurements of ROS level (**a**) and NAD^+^/NADH ratio (**b**) in mitochondria purified from tauopathy human iPSC neurons, rTg4510 mouse brains, human PSP and AD postmortem brains, and *Drosophila* brains expressing human tau-R406W, all respiring under FET condition. **c**, RET and FET assays of cells or purified mitochondria from isogenic WT control and tau-P301L iPSC neurons with or without CPT treatment. **d**, RET assay of purified mitochondria from tau-P301L iPSC neurons and rTg4510 mouse brains, with or without rotenone treatment during the assay. **e**, **f**, measurements of NAD^+^/NADH using the NAD^+^/NADH Assay Kit (WST-8) on rTg4510 mouse brain and tau-P301L iPSC neuron mitochondria (**e**), or measurement of NAD^+^/NADH in rTg4510 mouse brain mitochondria using targeted metabolomics by mass spectrometry (**f**). While the degree of measured changes between genotypes is variable, presumably reflecting methodological differences, the trend is the same. **g**, Mitochondrial respirometry analysis by Seahorse XF analyzer of the functional integrity of mitochondria purified from mouse brain tissues with the same method used to purify mitochondria for the RET and FET assays. **h**, **i**, Measurements of ROS level (**h**) and NAD^+^/NADH ratio (**i**) in mitochondria purified from human iPSC neurons expressing tau-WT or tau-P301L, respiring under the FET condition. All data are means ± SEM; statistical significance was determined by two-tailed unpaired Student’s *t* test (**a**, **b**, **e**-**i**) or one-way ANOVA with Tukey’s post hoc test (**c, d**). *p<0.05; **p<0.01; ***p<0.001; ****p<0.0001. For cell culture and fly studies, n=3-5 biological repeats. For mouse and patient tissue studies, each data point represents an individual mouse or patient sample.

**Extended Data Fig. 2.**
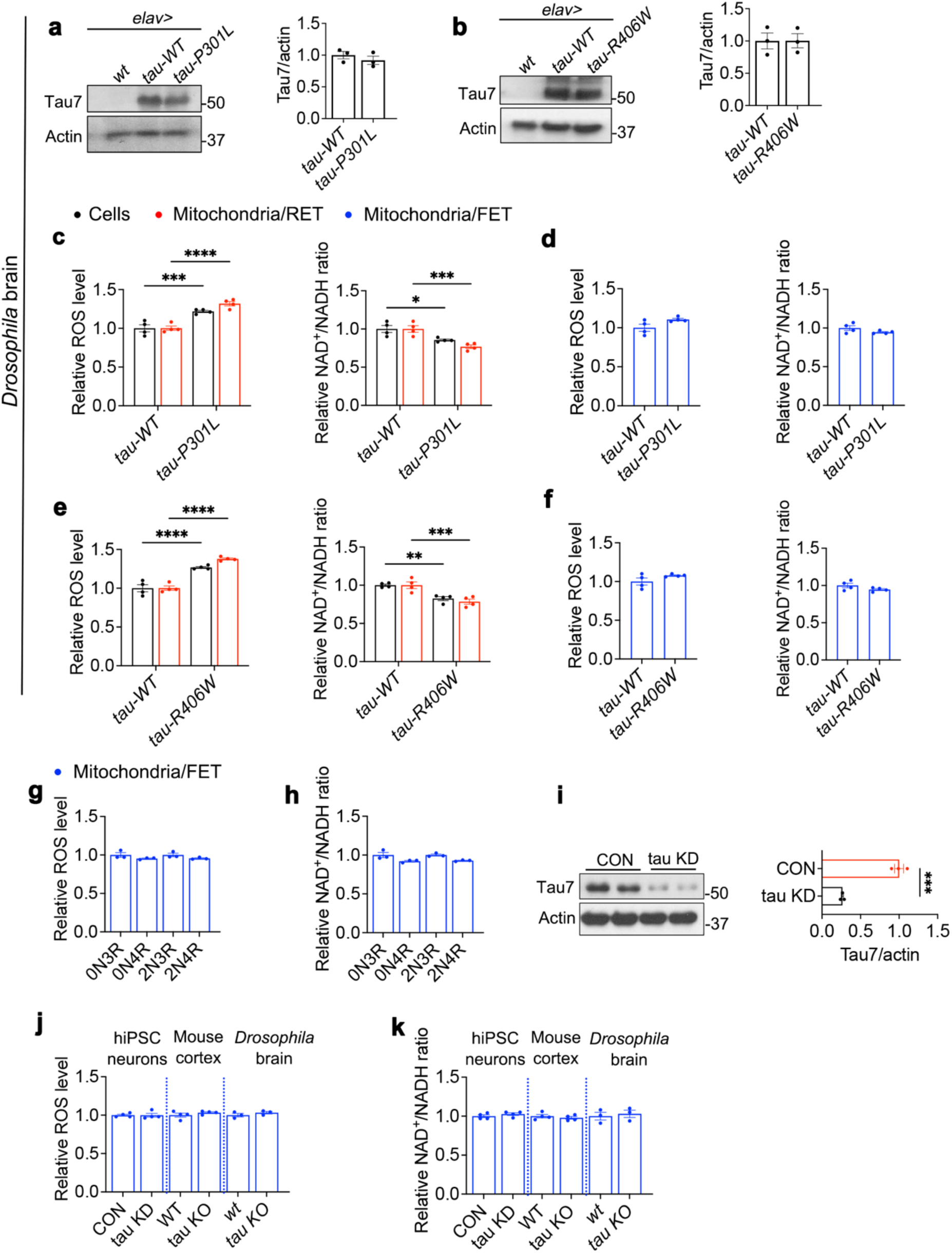
RET and FET-related changes in *Drosophila*, mouse, and human iPSC-derived neuron models after tau manipulation. **a**, WB analysis and quantification of tau-WT and tau-P301L expression in transgenic fly brain. **b**, WB analysis and quantification of tau-WT and tau-R406W expression in transgenic fly brain. **c**, **d**, Measurements of ROS and NAD^+^/NADH ratio of tau-WT or tau-P301L transgenic fly brain cells or mitochondria purified from these cells and respiring under RET (**c**) or FET (**d**) conditions. **e**, **f**, Measurements of ROS and NAD^+^/NADH ratio of tau-WT or tau-R406W transgenic fly brain cells or mitochondria purified from these cells and respiring under RET (**e**) or FET (**f**) conditions. **g**, **h**, Measurements of ROS level (**g**) and NAD^+^/NADH ratio (**h**) in mitochondria purified from human iPSC neurons overexpressing the various tau isoforms and respiring under FET condition. **i**, WB and quantification showing efficiency of tau KD in human iPSC neurons. **j**, **k**, Measurements of ROS level (**j**) and NAD^+^/NADH ratio (**k**) in mitochondria purified from control and tau KD human iPSC neurons, control and tau KO mouse cortex, or control and tau KO fly brains, respiring under the FET condition. All data are means ± SEM; statistical significance was determined by two-tailed unpaired Student’s *t* test. *p<0.05; **p<0.01; ***p<0.001; ****p<0.0001. For cell culture and fly studies, n=3-5 biological repeats; For mouse studies, each data point represents an individual mouse.

**Extended Data Fig. 3.**
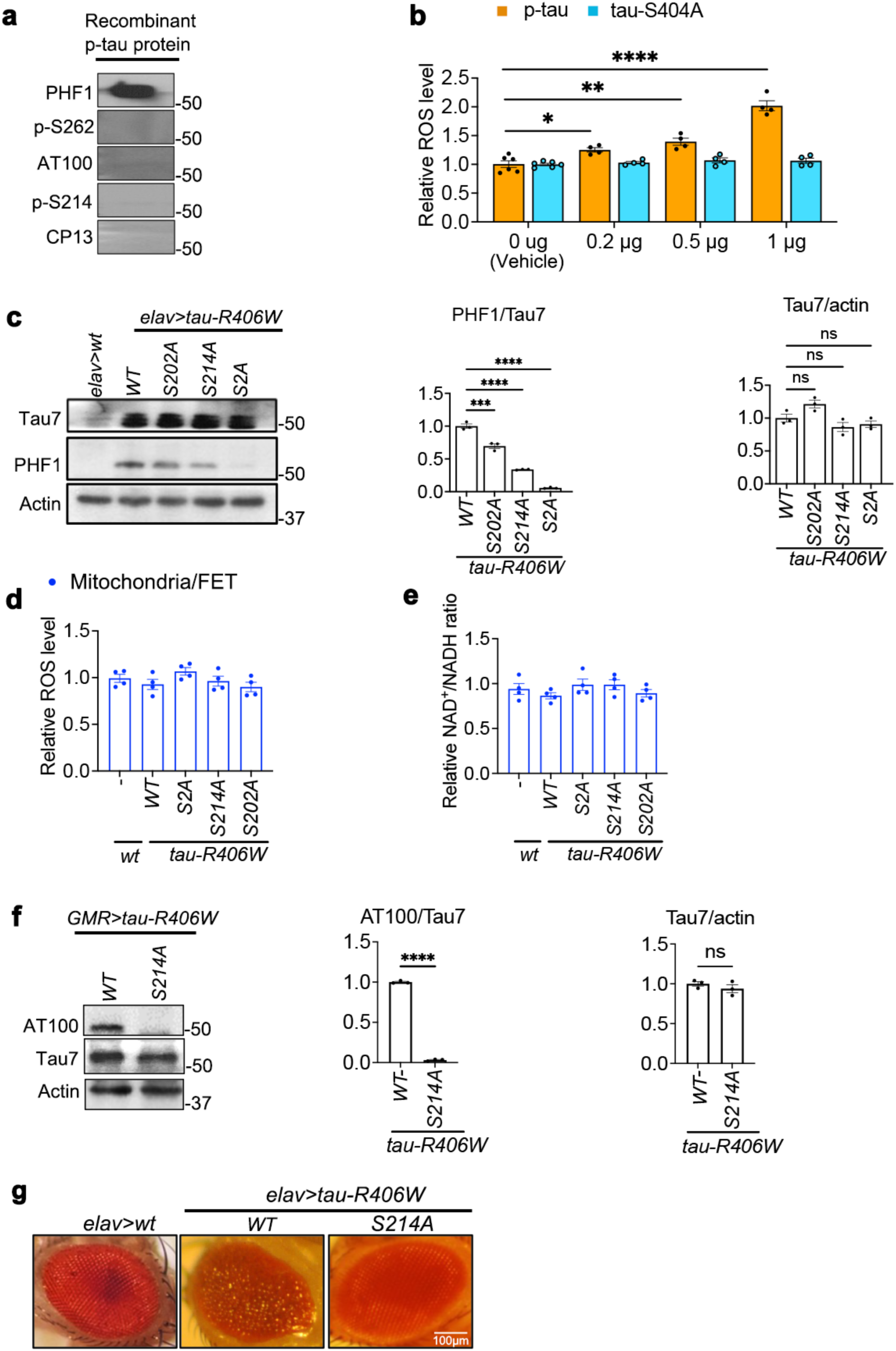
Data related to the phosphorylation-dependent function of tau in promoting RET. **a,** WB characterization of the recombinant human tau protein pretreated with GSK-3β and used in the *in vitro* mitochondrial import assays. **b**, Dose-dependent effect of phosphorylated (p-tau) and non-phosphorylatable (tau-S404A) recombinant tau proteins on RET-ROS. **c**, WB and quantification of the expression levels and extent of PHF-1 phosphorylation of the various tau-R406W phospho-mutants expressed in the *Drosophila* brain. **d, e**, Measurements of ROS (**d**) and NAD^+^/NADH ratio (**e**) in mitochondria purified from transgenic flies expressing WT or phospho-mutant forms of tau-R406W and respiring under the FET condition. **f**, WB analysis showing abolishment of the AT100 phospho-epitope by the S214A mutation in tau-R406W. **g**, Microscopy images of a control *Drosophila* eye (*elav>wt*) and *Drosophila* eyes expressing tau-R406W, or the tau-R406W-S214A phospho-mutant. The roughness of the eye surface caused by tau-R406W-induced photoreceptor degeneration was abolished by the S214A mutation. All data are means ± SEM; statistical significance was determined by one-way ANOVA with Tukey’s post hoc test (**c-e**), or two-tailed unpaired Student’s *t* test (**b, f**). *p<0.05; **p<0.01; ***p<0.001; ****p<0.0001. Each data point represents an independent experimental repeat.

**Extended Data Fig. 4.**
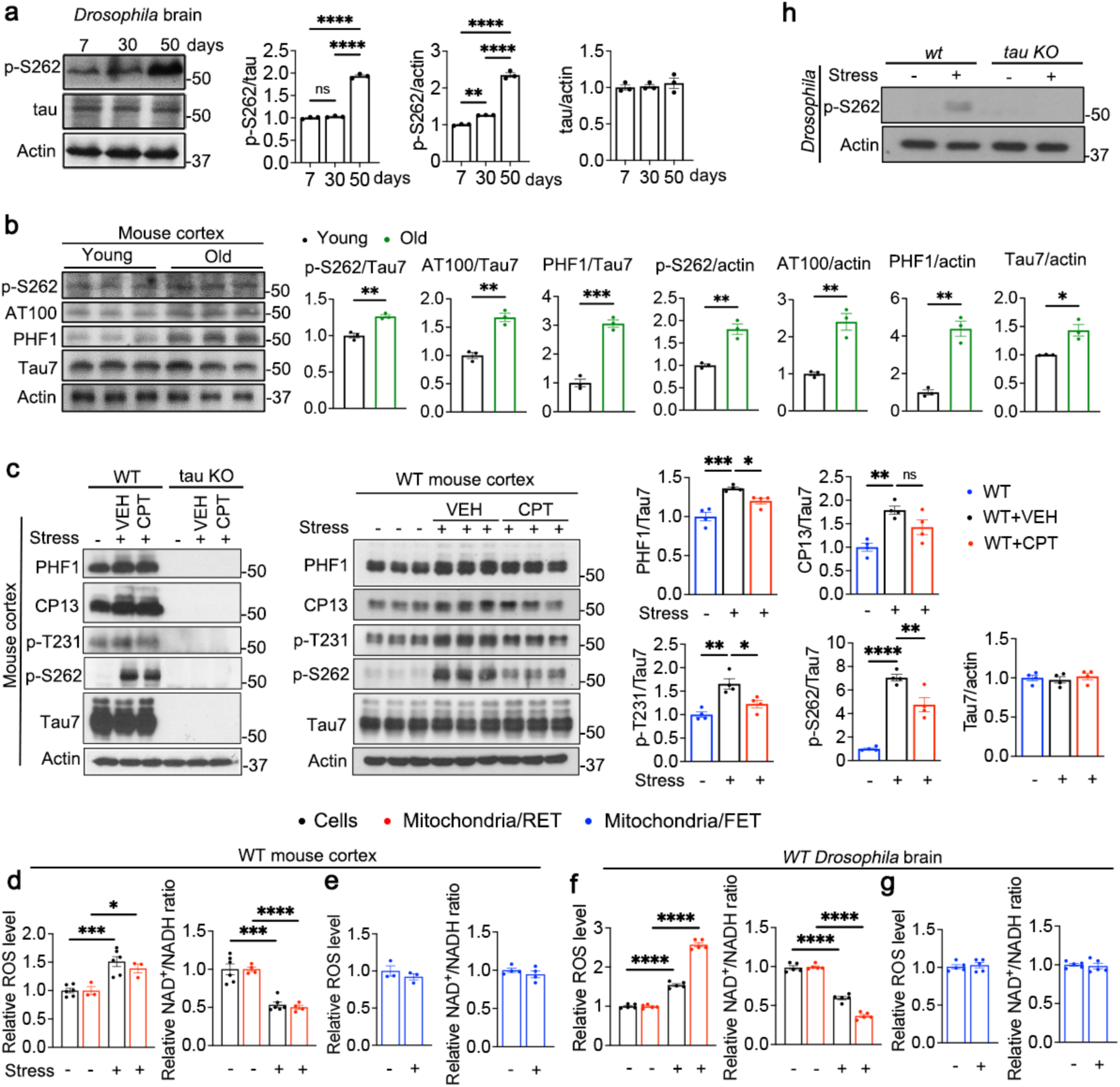
RET activation and tau hyperphosphorylation during stress. **a,** WB and quantification showing effect of aging on S262 phosphorylation of endogenous fly tau. **b**, WB and quantification showing effect of aging on endogenous mouse tau phosphorylation at S262, AT100, and PHF-1 sites. p-tau level is normalized with total tau or actin in **a** and **b**. **c**, WB and quantification of p-tau levels in WT mouse cortex of non-stressed control, heat-stressed, and heat-stressed plus CPT-treated animals, as well as the cortex of non-stressed control, heat-stressed, and heat-stress plus CPT-treated tau KO mice. **d**, **e**, Measurements of ROS and NAD^+^/NADH ratio of non-stressed or stressed WT mouse cortex cells, or mitochondria purified from these cells and respiring under RET (**d**) or FET (**e**) conditions. **f**, **g**, Measurements of ROS and NAD^+^/NADH ratio of non-stressed or heat-stressed WT *Drosophila* brain cells or mitochondria purified from these cells and respiring under RET (**f**) or FET (**g**) conditions. **h**, WB analysis of heat stress-induced p-S262 tau level in WT and tau KO fly brain. All data are means ± SEM; statistical significance was determined by one-way ANOVA with Tukey’s post hoc test (**a, c**), or two-tailed unpaired Student’s *t* test (**b, d, e, f, g**). *p<0.05; **p<0.01; ***p<0.001; ****p<0.0001. For fly studies, n=3-5 biological repeats; For mouse studies, each data point represents an individual mouse.

**Extended Data Fig. 5.**
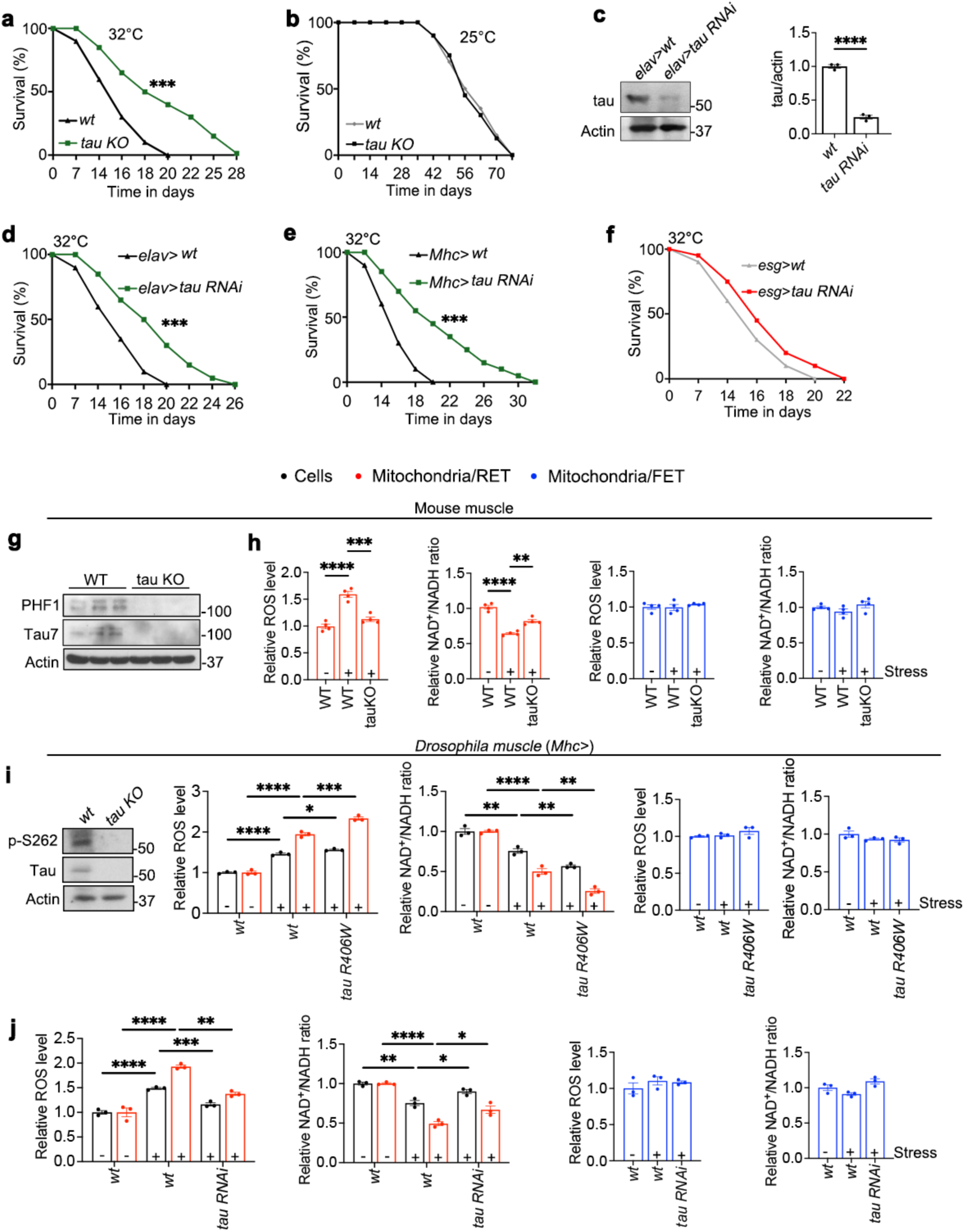
Muscle expression of tau and its importance to fitness. **a, b,** Survival curves of control and tau KO flies raised at 32°C (**a**) or 25°C (**b**). **c**, WB and quantification showing the knockdown efficiency of tau RNAi in the fly brain. **d**-**f**, Survival curves of control and various tissue-specific (**d**: *elav*> neuronal; **e**: *Mhc*> muscle; **f**: *esg*> gut) tau RNAi flies raised at 32°C. **g**, WB showing tau expression in the mouse muscle that is absent in tau KO mice. Note that mouse muscle tau appeared to be running at a higher molecular weight than neuronal tau, suggesting that it might represent a different tau isoform. **h**, RET and FET assays showing the effect of tau KO on heat stress-induced RET. **i**, WB showing tau expression in the fly muscle. **j**, **k**, RET and FET assays showing tau-R406W overexpression promotes (**j**), whereas tau KO inhibits (**k**) heat stress-induced RET in the fly muscle. All data are means ± SEM; statistical significance was determined by one-way ANOVA with Tukey’s post hoc test (**h, j, k**), two-tailed unpaired Student’s *t* test (**c, g, i**), or Kaplan Meier survival analysis (**a, b, d, e, f**). Each group contains 20 flies in survival assay. *p<0.05; **p<0.01; ***p<0.001; ****p<0.0001. Each data point in **h, j, k** represents an independent experimental repeat.

**Extended Data Fig. 6.**
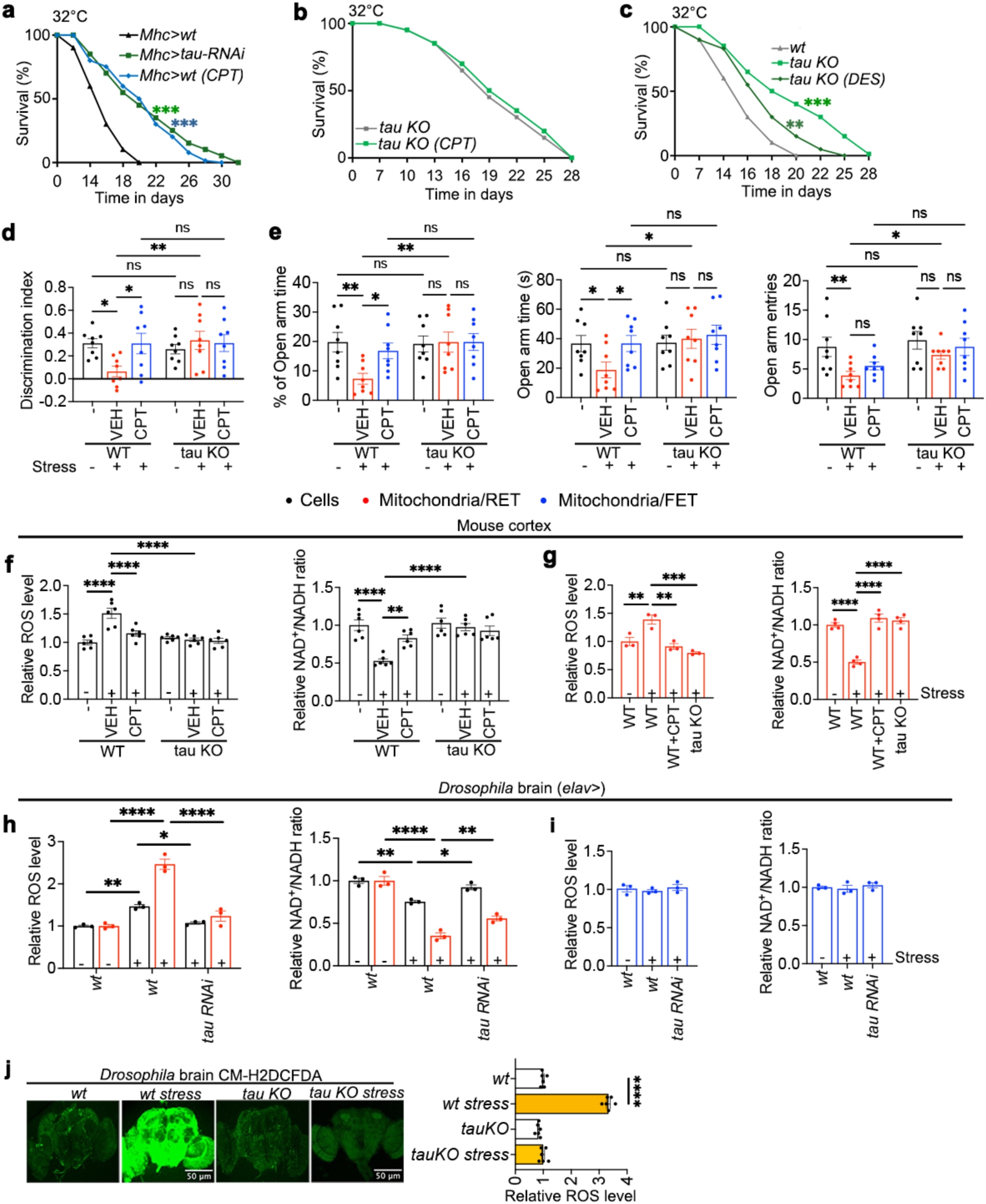
Effect of tau knockout or knockdown on RET-related fitness and survival in flies and mice. **a-c,** Survival curves of muscle-specific tau RNAi flies (**a**) or tau KO flies (**b, c**) raised at 32°C and treated with CPT (**a, b**) or DES (**c**). **d**, **e**, Behavior of WT and tau KO mice under heat stress and with or without CPT treatment in the novel object recognition (**d**) and elevated plus maze (**e**) tests. **f**, **g**, RET assays of cells (**f**) or purified mitochondria (**g**) from the cortex of WT and tau KO mice under heat stress and with or without CPT treatment. **h**, **i**, RET (**h**) and FET (**i**) assays of cells or purified mitochondria from brain tissues of control and neuronal tau RNAi flies under heat stress. **j**, Images and quantification showing that heat stress-induced brain ROS in wild type flies was abolished by tau KO. All data are means ± SEM; statistical significance was determined by two-way ANOVA with Tukey’s post hoc test (**d, f, j**), two-way ANOVA with Fisher’s LSD test (**e**), one-way ANOVA with Tukey’s post hoc test (**g, h, i**), or Kaplan Meier survival analysis (**a-c**). Each group contains 20 flies in survival assay. *p<0.05; **p<0.01; ***p<0.001; ****p<0.0001. For fly studies, n=3-5 biological repeats. For mouse studies, each data point represents an individual animal.

**Extended Data Fig. 7.**
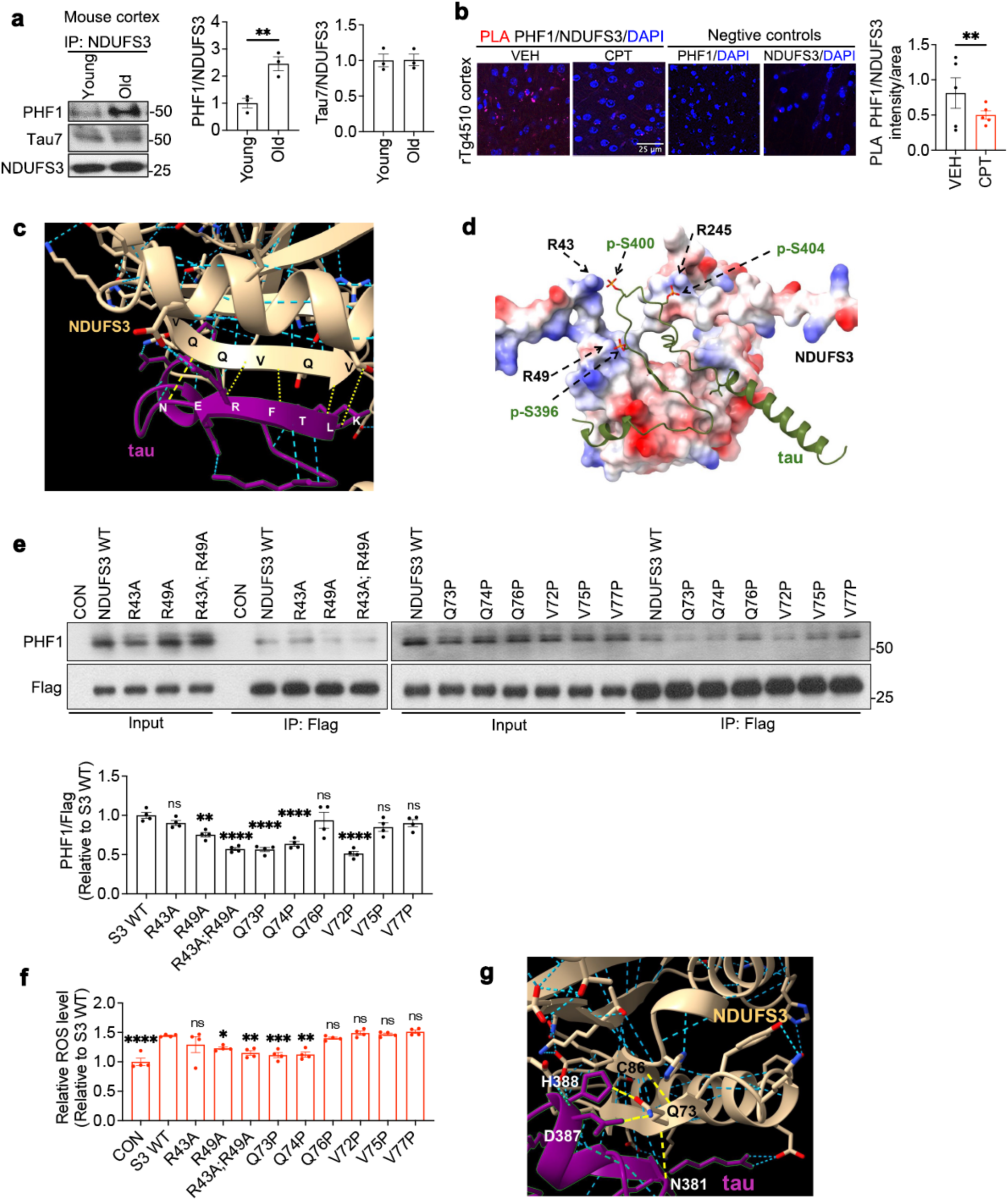
Physical interaction between tau and mitochondrial C-I protein NDUFS3 and effect of perturbed tau-NDUFS3 interaction on RET. **a,** Co-IP assay and quantification showing increased p-tau/NDUFS3 interaction in aged mouse brains. **b**, PLA and quantification showing PHF-1 tau/NDUFS3 interaction in rTg4510 mouse brain that is attenuated by CPT. **c**, **d**, AlphaFold-predicted β-strand interaction between the C-terminal KLTFRNE of tau and the VQQVQV peptide of NDUFS3 (**c**) and electrostatic interaction between the p-S396/pS400/pS404 residues in the PHF-1 epitope of tau and positively charged R49, R43, and R245 residues of NDUFS3, respectively (**d**). Hydrogen bonds formed between backbone carbonyl and amide groups are shown in yellow dashed lines in **c**. The surface charges of NDUFS3 are shown in blue for positive charges and red for negative charges in **d**. R245 is not surface exposed in the solved mouse C-I cryo-EM structure so it was not analyzed in this work. **e**, WB and quantification showing co-IP between PHF-1 p-tau and the Flag-tagged WT and mutant NDUFS3 proteins expressed in HEK293T cells co-transfected with tau-P301L. **f**, RET-ROS measurement in HEK293T cells co-expressing tau-P301L and the WT or mutant forms of NDUFS3. Note that although the V72P mutant perturbed p-tau/NDUFS3 interaction, its effect on RET-ROS was not significant, possibly due to effect of this mutation on NDUFS3 function and ROS production by additional mechanisms. **g**, AlphaFold predicted hydrogen-bonding between the Q73 of NDUFS3 and residues in tau and NDUFS3 through sidechain and backbone groups. Hydrogen bonds formed by Q73 are highlighted in yellow dashed lines. All data are means ± SEM; statistical significance was determined by two-tailed unpaired Student’s *t* test (**a, b**), or one-way ANOVA with Dunnett’s multiple test (**e, f**). *p<0.05; **p<0.01; ***p<0.001; ****p<0.0001. Each data point in **e** and **f** represents an independent experimental repeat.

**Extended Data Fig. 8.**
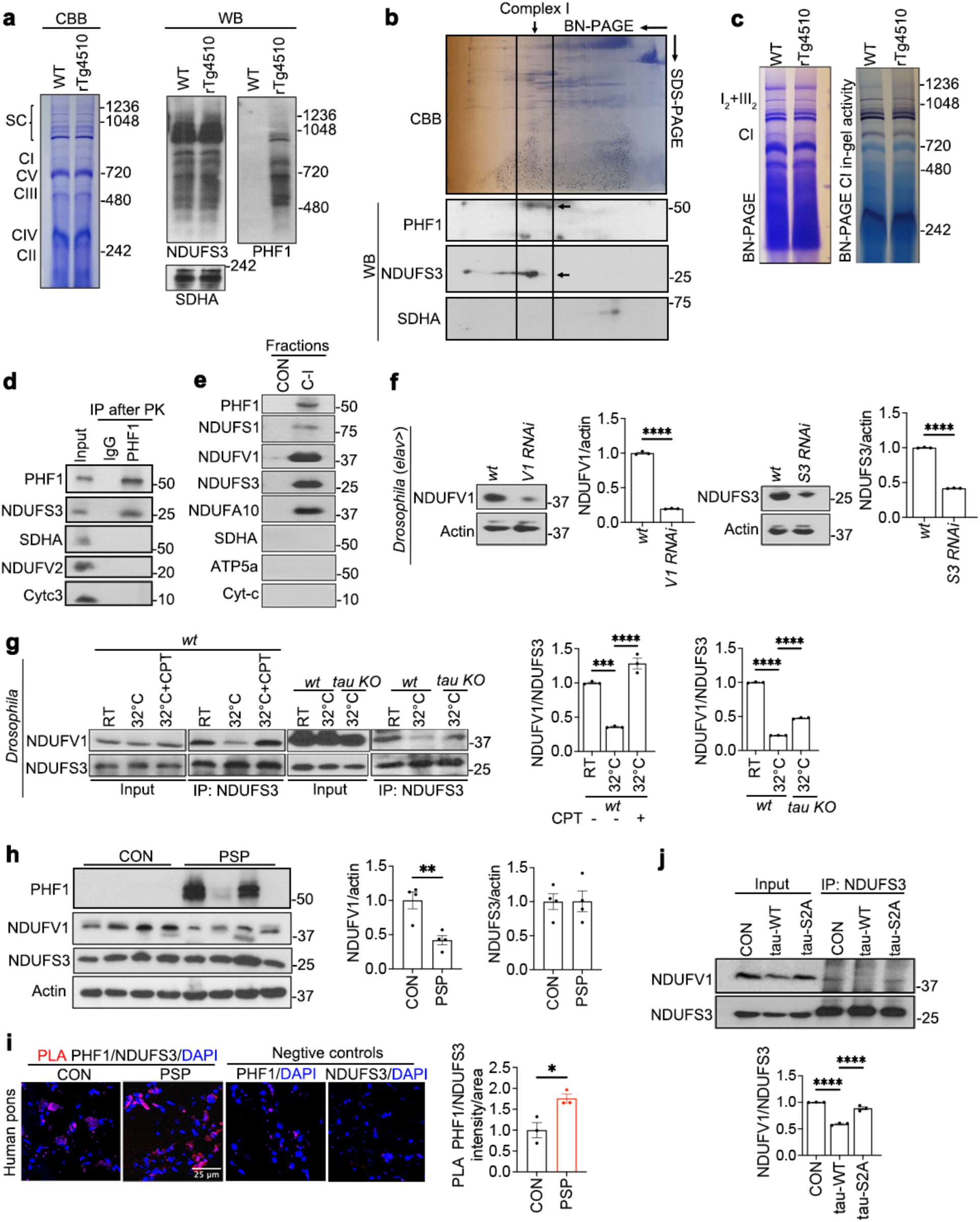
Physical interaction between tau and NDUFS3 and the effect of tau on NDUFS3 interaction with NDUFV1. **a**, BN-PAGE separated mitochondrial membranes from WT and rTg4510 mouse brains were stained with CBB (left) or probed with NDUFS3, PHF-1, and SDHA antibodies by WB. **b**, Mitochondrial membrane from rTg4510 mouse brain separated by 2D-PAGE, with BN-PAGE and SDS-PAGE as first and second dimensions, respectively, and stained with CBB (upper) or probed with NDUFS3, PHF-1, and SDHA antibodies by WB (lower). C-I position is marked. **c**, In-gel C-I activity assay of BN-PAGE separated brain mitochondrial membranes. **d**, Co-IP assay using PK-treated mitochondria from rTg4510 mouse brain. **e**, WB analysis of control and C-I fractions purified by high-performance ion exchange and size exclusion chromatography from rTg4510 mouse brain. SDHA, ATP5A, and Cyt *c* serve as specificity controls. **f**, WB and quantification verifying antibody specificity using NDUFS3 and NDUFV1 RNAi samples. **g**, Co-IP assay and quantification showing the effect of heat stress on NDUFS3-NDUFV1 interaction in fly brain tissues and the effect of CPT treatment or tau KO. **h**, WB analysis and quantification of NDUFV1 and NDUFS3 in control and PSP patient brain samples. **i**, PLA assay of PHF-1 tau/NDUFS3 interaction in PSP brain tissue pons region. **j**, Co-IP assay and quantification of the effect of tau-WT and tau-S2A on NDUFS3/NDUFV1 interaction in transfected HEK293T cells. All data are means ± SEM; statistical significance was determined by two-tailed unpaired Student’s *t* test (**f, h, i**), or one-way ANOVA with Tukey’s post hoc test (**g, j**). *p<0.05; **p<0.01; ***p<0.001; ****p<0.0001. Each data point in **j** represents an independent experimental repeat. For patient tissue studies, each data point represents an individual patient sample. For mouse tissue studies, each data point represents an individual mouse sample. **i**: n=3 mice/group, each data point represents an average of 6 images/mouse.

**Extended Data Fig. 9.**
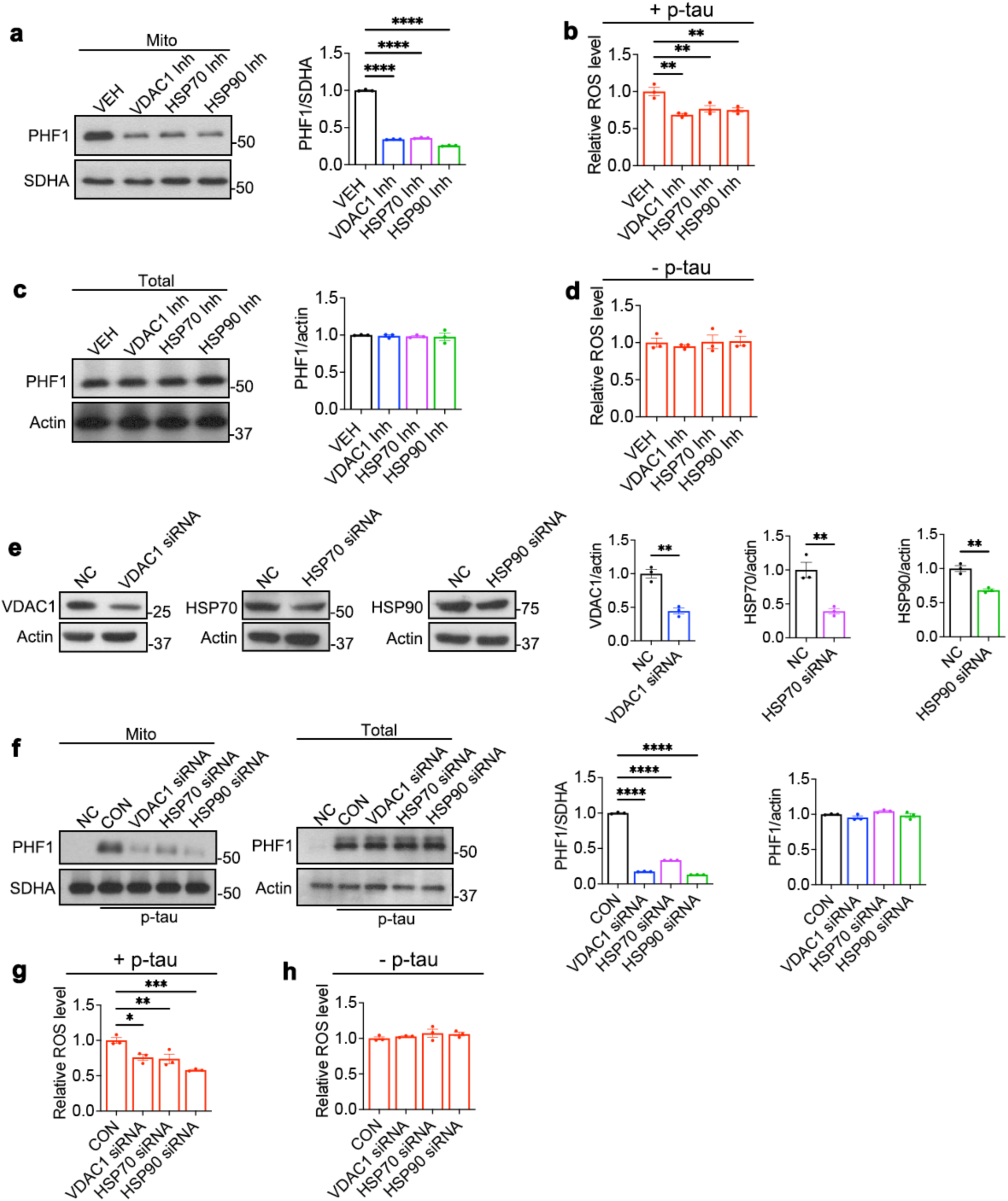
Effect of inhibiting tau entry into mitochondria on RET *in vitro*. **a,** Representative WB and quantification showing effect of VDAC, Hsp70, and Hsp90 inhibitor treatment on tau entry into mitochondria in the *in vitro* import assay. Mitochondrial p-tau level is normalized with the C-II protein SDHA. **b**, RET-ROS measurements after p-tau import into control or inhibitor-treated mitochondria. **c**, Representative WB and quantification showing total PHF-1 tau level in the *in vitro* import assay mixture. Total p-tau level is normalized with actin, which is known to be associated with mitochondria. **d**, RET-ROS measurements in control or inhibitor-treated mitochondria without p-tau import. **e**, WBs and quantification showing knockdown efficiency by VDAC1, Hsp70, and Hsp90 siRNAs. **f**, Representative WBs and quantification showing the effect of VDAC1, Hsp70, and Hsp90 siRNAs on tau entry into mitochondria in the *in vitro* import assay and total PHF-1 tau level in the *in vitro* import assay. **g**, **h**, RET-ROS measurements in mitochondria from control or siRNA treated cells with (**g**) or without (**h**) p-tau import into mitochondria. All data are means ± SEM; statistical significance was determined by one-way ANOVA with Dunnett’s multiple test (**a, b, c, d, f, g, h**), or two-tailed unpaired Student’s *t* test (**e**). *p<0.05; **p<0.01; ***p<0.001; ****p<0.0001. Each data point in **a**-**h** represents an independent experimental repeat.

**Extended Data Fig. 10.**
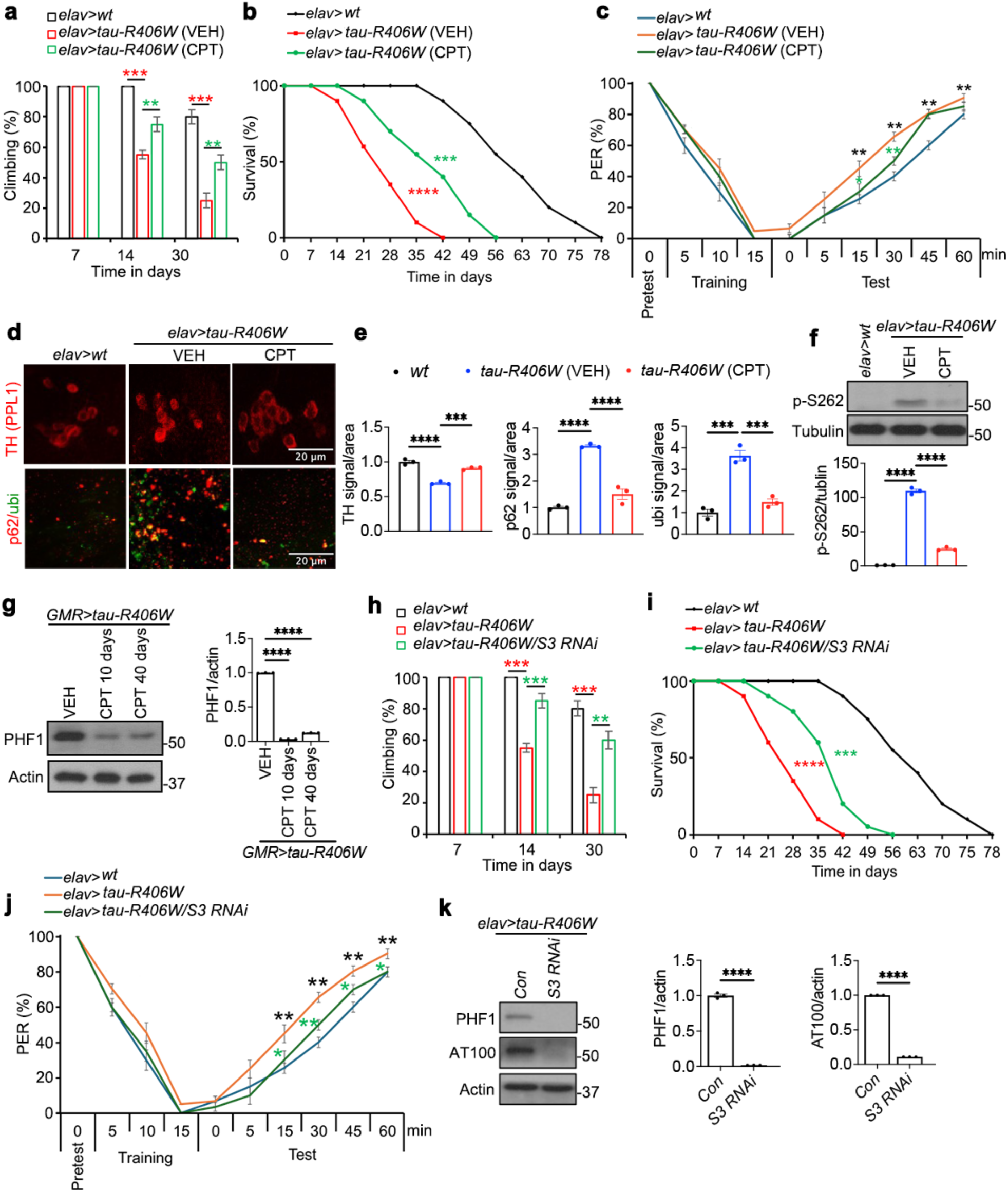
Effect of RET inhibition on behavioral deficit, neuropathology, and survival in a tauopathy fly model. **a,** climbing activity assay of control flies and neuronal tau-R406W expressing flies with or without CPT treatment at different ages. **b**, Survival curve of control flies and neuronal tau-R406W expression flies with or without CPT treatment. **c**, Aversive taste memory assay of control flies and neuronal tau-R406W flies with or without CPT treatment. **d**, **e**, Immunostaining (**d**) and quantification (**e**) of signal intensities of TH-positive PPL1 cluster dopaminergic neurons and p62 and ubiquitin (ubi) positive protein aggregates in the brains of control flies and neuronal tau-R406W flies with or without CPT treatment. **f**, **g**, WB and quantification showing effect of CPT on p-S262 (**f**) and PHF-1 p-tau (**g**) level in pan-neuronal *elav-Gal4*> (**f**) or photoreceptor neuron-specific *GMR-Gal4*> (**g**) tau-R406W expression flies. **h**-**j**, Effect of RET inhibition by NDUFS3 RNAi on climbing activity (**h**), survival (**i**), and aversive taste memory (**j**) of pan-neuronal tau-R406W expression flies. **k**, WB and quantification showing effect of NDUFS3 RNAi on p-tau levels in pan-neuronal tau-R406W flies. All data are means ± SEM; statistical significance was determined by one-way ANOVA with Tukey’s post hoc test (**a, e, f, g, h**), group analysis using multiple *t* test with Sidak-Bonferroni multiple comparison (**c**, **j**), two-tailed unpaired Student’s *t* test (**k**), or Kaplan Meier survival analysis (**b, i**). *p<0.05; **p<0.01; ***p<0.001; ****p<0.0001. Each data point represents independent experimental repeat.

**Extended Data Fig. 11.**
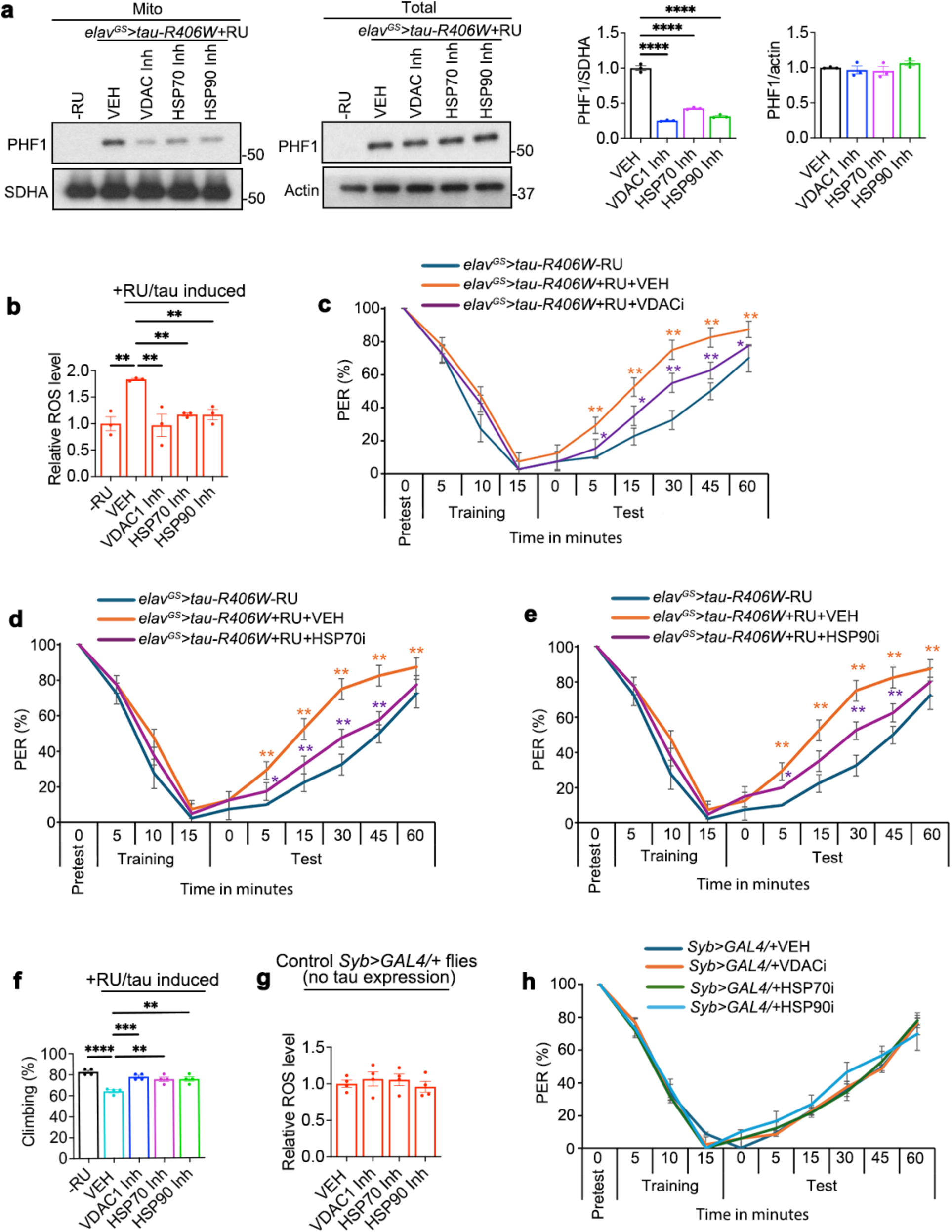
Effect of inhibiting tau entry into mitochondria on RET and behavior in a tauopathy *Drosophila* model. **a,** Representative WBs and quantification showing effect of VDAC, Hsp70, and Hsp90 inhibitors on the mitochondrial level of PHF-1 tau and total PHF-1 tau in the *elav-GS>tau-R406W* flies. In this GeneSwitch inducible model, tau-R406W expression is tightly controlled by the addition of RU486 to the fly food. Mitochondrial fractions or total cell lysates were used for WB, and mitochondrial or total PHF-1 p-tau was normalized by SDHA or actin. **b**, RET-ROS measurements in the mitochondria from control or inhibitor treated *elav-GS>tau-R406W* flies after tau induction by RU486. **c**-**e**, aversive taste memory assays in VDACi (**c**), Hsp70i (**d**), and Hsp90i (**e**) treated *elav-GS>tau-R406W* flies after tau induction by RU486. **f**, Climbing activity assay in VDACi, Hsp70i, and Hsp90i treated *elav-GS>tau-R406W* flies after tau induction by RU486. **g**, **h**, RET (**g**) and aversive taste memory (**h**) assays of control flies without tau transgene expression. All data are means ± SEM; statistical significance was determined by one-way ANOVA with Dunnett’s multiple test (**a, b, f, g**), or group analysis using multiple *t* test with Sidak-Bonferroni multiple comparison (**c**, **d**, **e**, **h**). *p<0.05, **p<0.01, ***p<0.001, ****p<0.0001. Each data point represents independent experimental repeat. Three sets of flies with 10-12 flies in each set were used for the behavioral assays.

**Extended Data Fig. 12.**
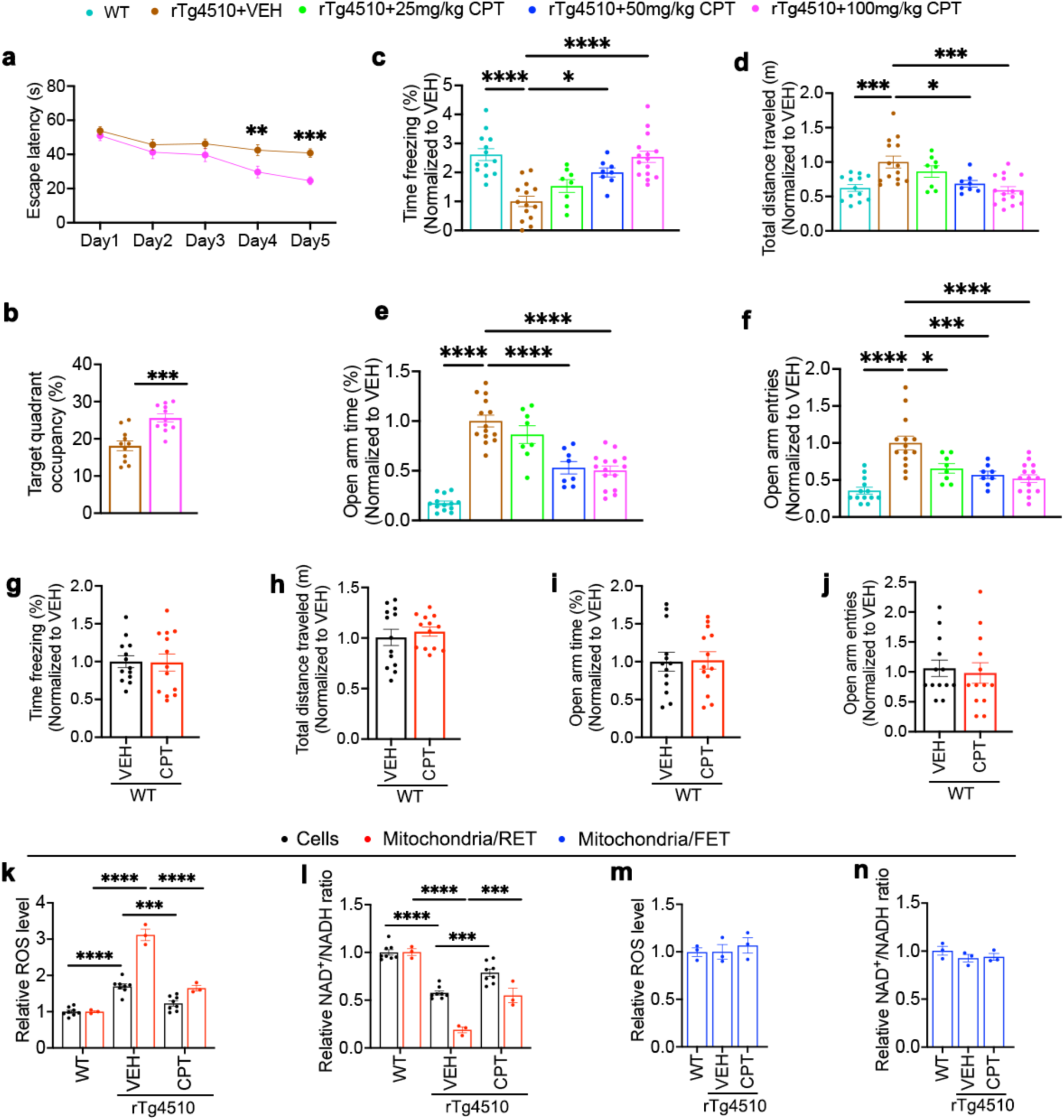
Effect of RET inhibition on behavioral deficits in rTg4510 tauopathy mice. **a, b,** Effect of CPT treatment on escape latency (**a**) and target quadrant occupancy (**b**) in the Morris water maze (MWM) assay of rTg4510 mice. **c**-**f**, Dose-response of oral CPT administration at 25, 50, or 100 mg/kg on behavior in the contextual fear conditioning (**c**), open field (**d**), and elevated plus maze (EPM) (**e**, **f**) tests. **g**-**j**, Effect of CPT treatment on behavior in the contextual fear conditioning (**g**), open field (**h**), and EPM (**i**, **j**) tests of C57BL/6 wildtype mice. **k**-**n**, RET (**k**, **l**) and FET (**m**, **n**) assays of brain cells or purified mitochondria from rTg4510 mice treated with vehicle or CPT. All data are means ± SEM; statistical significance was determined by one-way ANOVA with Tukey’s post hoc test (**c-f**, **k-n**), or two-tailed unpaired Student’s *t* test (**a, b, g-j**). *p<0.05; **p<0.01; ***p<0.001; ****p<0.0001 (**a, b**: n=11 mice/group; **c-f**: WT, n=13; VEH, n=14; 25 mg/kg CPT and 50 mg/kg CPT, n=8 mice each; 100 mg/kg CPT, n=15 mice; **g-j**: n=11 mice/group; **k-n**, each data point represents an individual mouse).

**Extended Data Fig. 13.**
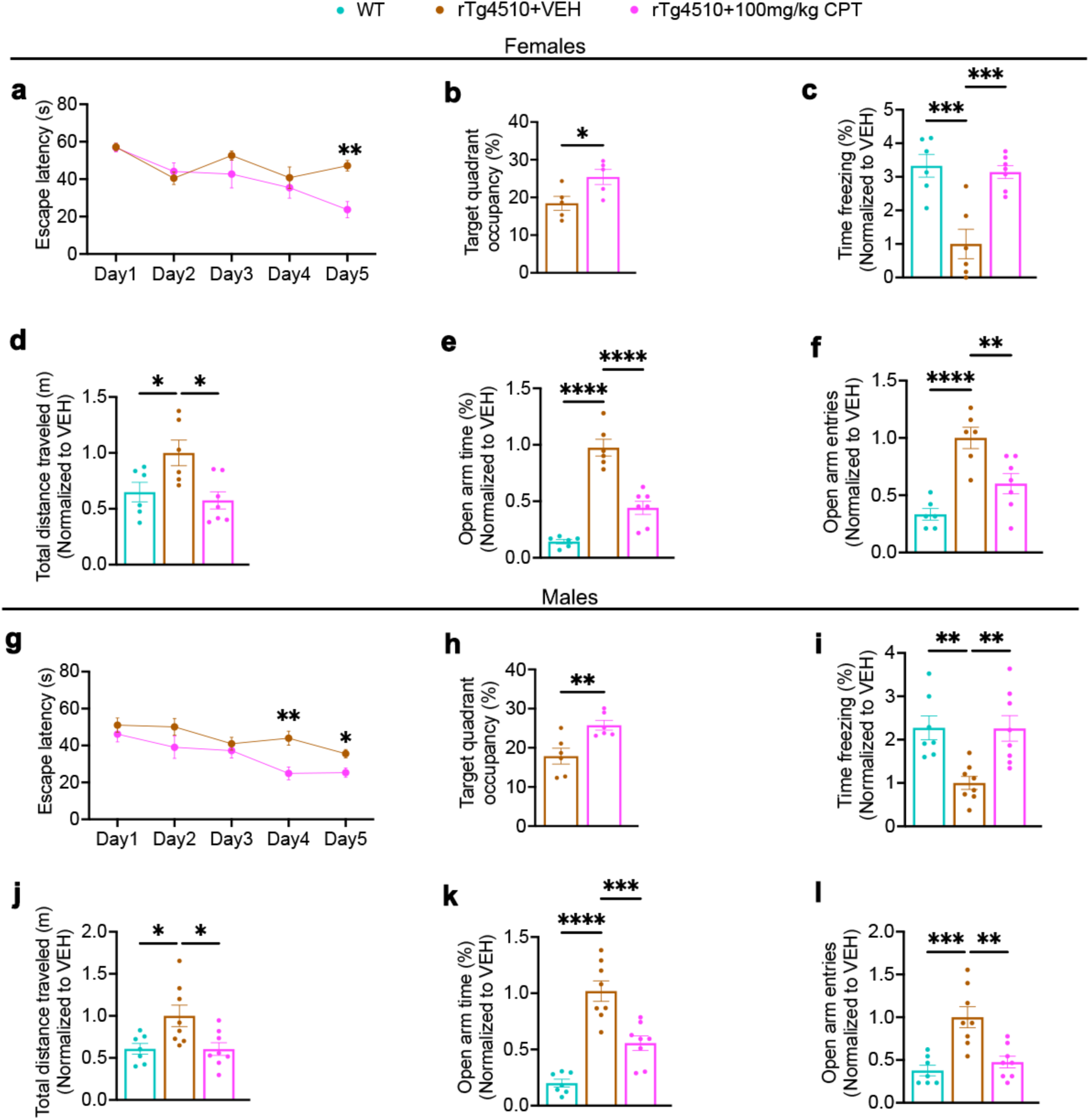
No sex-specific effect of RET inhibition on behavioral deficits in rTg4510 tauopathy mice. Female (**a-f**) and male (**g**-**l**) mice were separately analyzed for the MWM (**a, b, g, h**), fear conditioning (**c, i**), open field (**d, j**), and EPM (**e, f, k, l**) tests. All data are means ± SEM; statistical significance was determined by one-way ANOVA with Tukey’s post hoc test (**c-f**, **i-l**), or two-tailed unpaired Student’s *t* test (**a, b, g, h**). *p<0.05; **p<0.01; ***p<0.001; ****p<0.0001 (**a, b**: n=5 mice/group; **c-f**: WT, n=6; VEH, n=6; CPT, n=7 mice; **i-l**: WT, n=7; VEH, n=8; CPT, n=8 mice).

**Extended Data Fig. 14.**
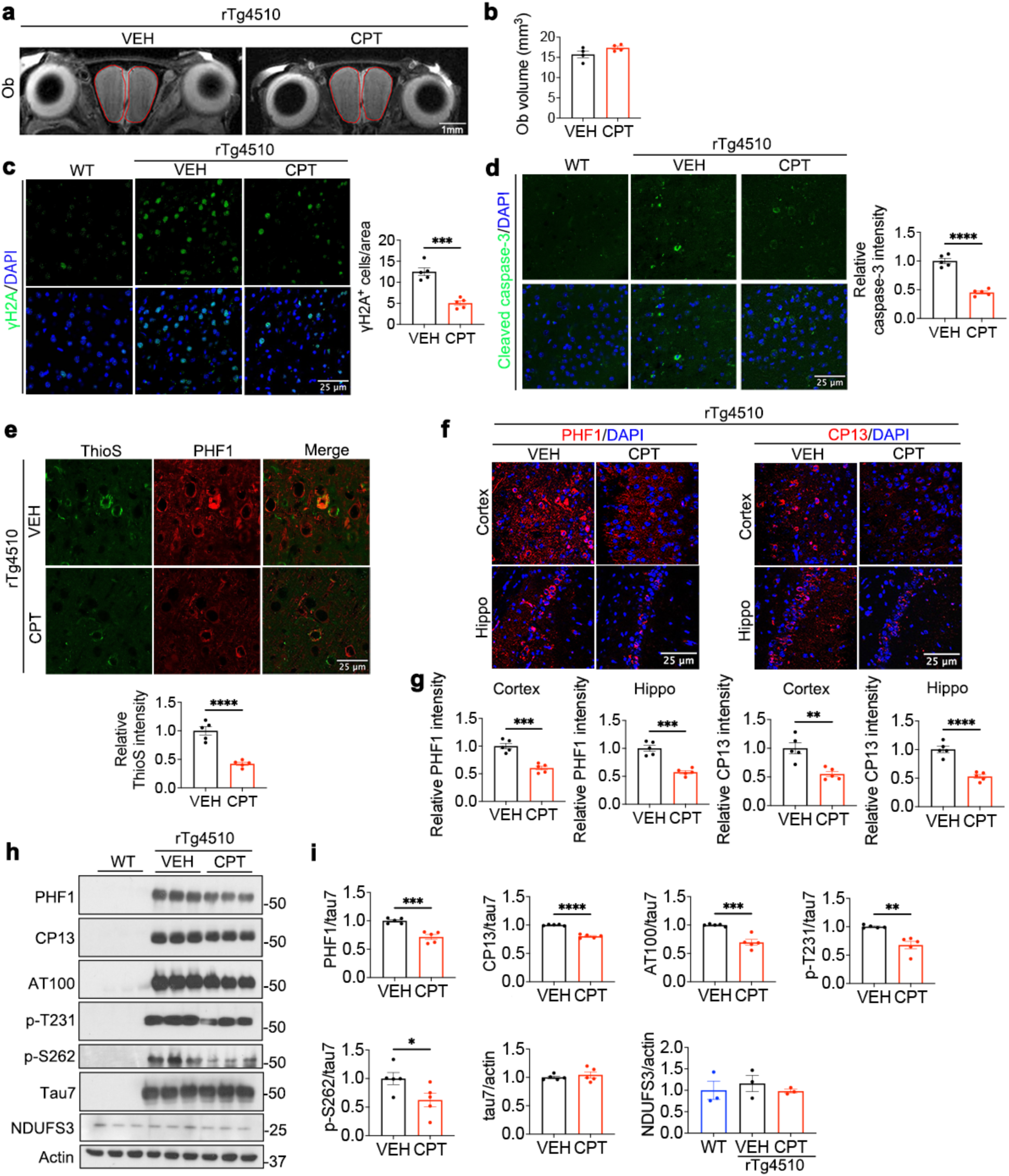
Effect of RET inhibition on tau-induced neuropathology and tau phosphorylation in rTg4510 tauopathy mice. **a, b**, Representative MRI images (**a**) and data quantification (**b**) of olfactory bulb (OB) volume. **c-e**, Representative images and data quantification of DNA-damage marker γH2AX (**c**), apoptosis marker cleaved caspase 3 (**d**), and tau aggregation as indicated by ThioS staining (**e**) in brain sections of rTg4510 mice treated with vehicle or CPT. **f**, **g**, Representative images (**f**) and quantification (**g**) of PHF-1 and CP13 immunostaining in the hippocampal CA1 and cortex regions of rTg4510 mouse brain. **h, i**, Immunoblots (**h**) and data quantification (**i**) of p-tau (PHF-1, CP13, AT100, p-T231, p-S262), total tau (HT7), and NDUFS3 levels in the cortex of 8-month-old rTg4510 mice. p-tau levels were normalized with total tau. All data are means ± SEM; statistical significance was determined by two-tailed unpaired Student’s *t* test; NDUFS3/actin in **i** was quantified by one-way ANOVA with Tukey’s post hoc test. *p<0.05; **p<0.01; ***p<0.001; ****p<0.0001 (**b**: n=4 mice/group; **c-e**, **g**: n=5 mice/group, each data point represents an average of 6 images/mouse; **i**: each data point represents an individual mouse).

**Extended Data Fig. 15.**
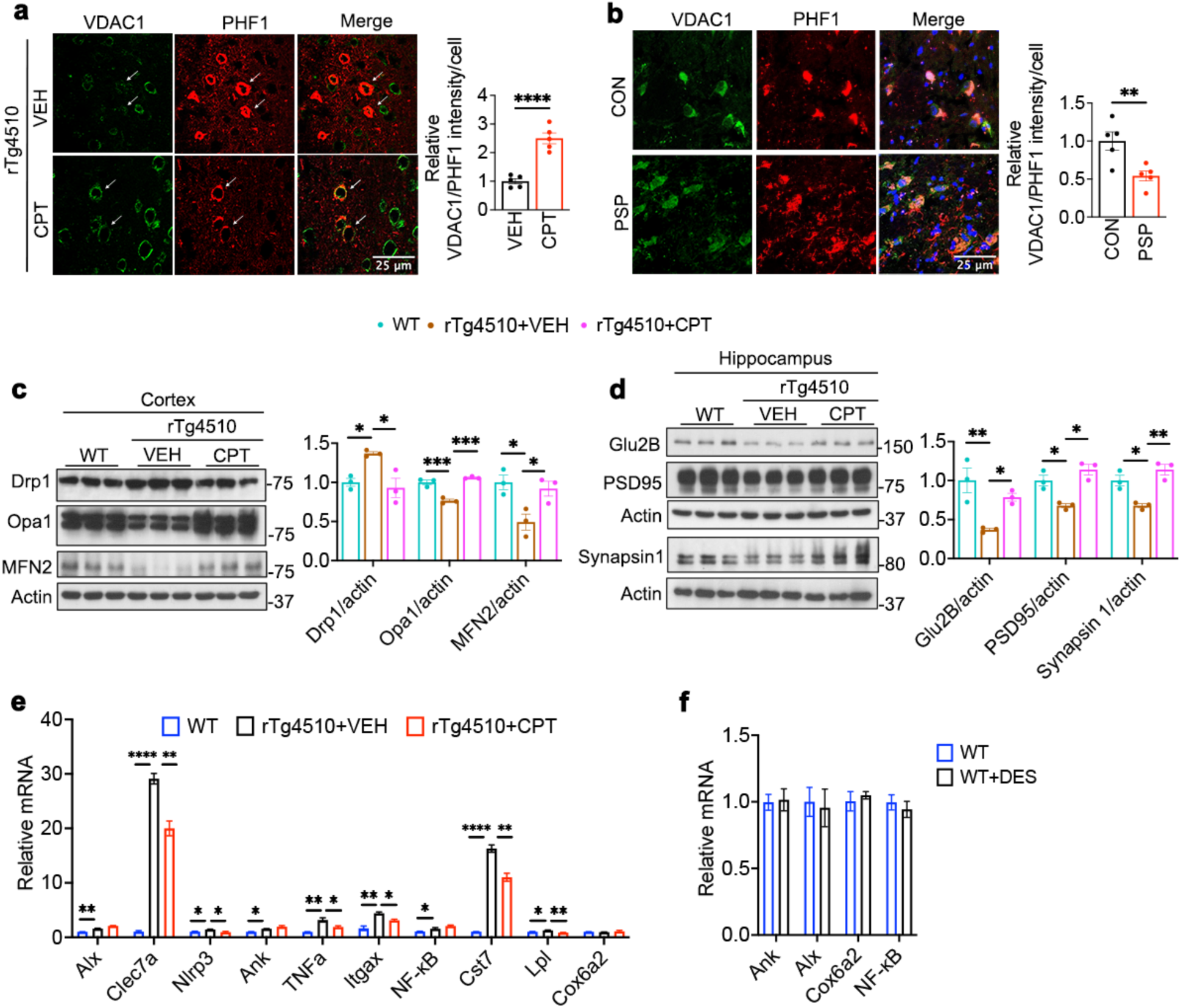
Further analysis of the effect of RET inhibition on tau-induced cellular and molecular pathology in rTg4510 mice. **a, b**, Representative images and data quantification of PHF-1 and VDAC1 levels in the cortex of rTg4510 mice with or without CPT treatment (**a**) or in PSP patient brain samples (**b**). **c**, **d**, WB and quantification of mitochondrial dynamics related proteins (**c**) or synaptogenesis related proteins (**d**) in the hippocampal tissues of rTg4510 mice with or without CPT treatment. **e**, **f**, RT-PCR analysis of neuroinflammation related transcripts in the hippocampal tissues of rTg4510 mice with or without CPT treatment (**e**), or wild type mice with or without DES treatment (**f**). All data are means ± SEM; statistical significance was determined by one-way ANOVA with Tukey’s post hoc test (**c, d, e**), or two-tailed unpaired Student’s *t* test (**a, b, f**). *p<0.05; **p<0.01; ***p<0.001; ****p<0.0001 (**a, b**: n=5 mice/group, each data point represents an average of 8 cells/mouse; **c-f**: n=3 mice/group).

**Extended Data Fig. 16.**
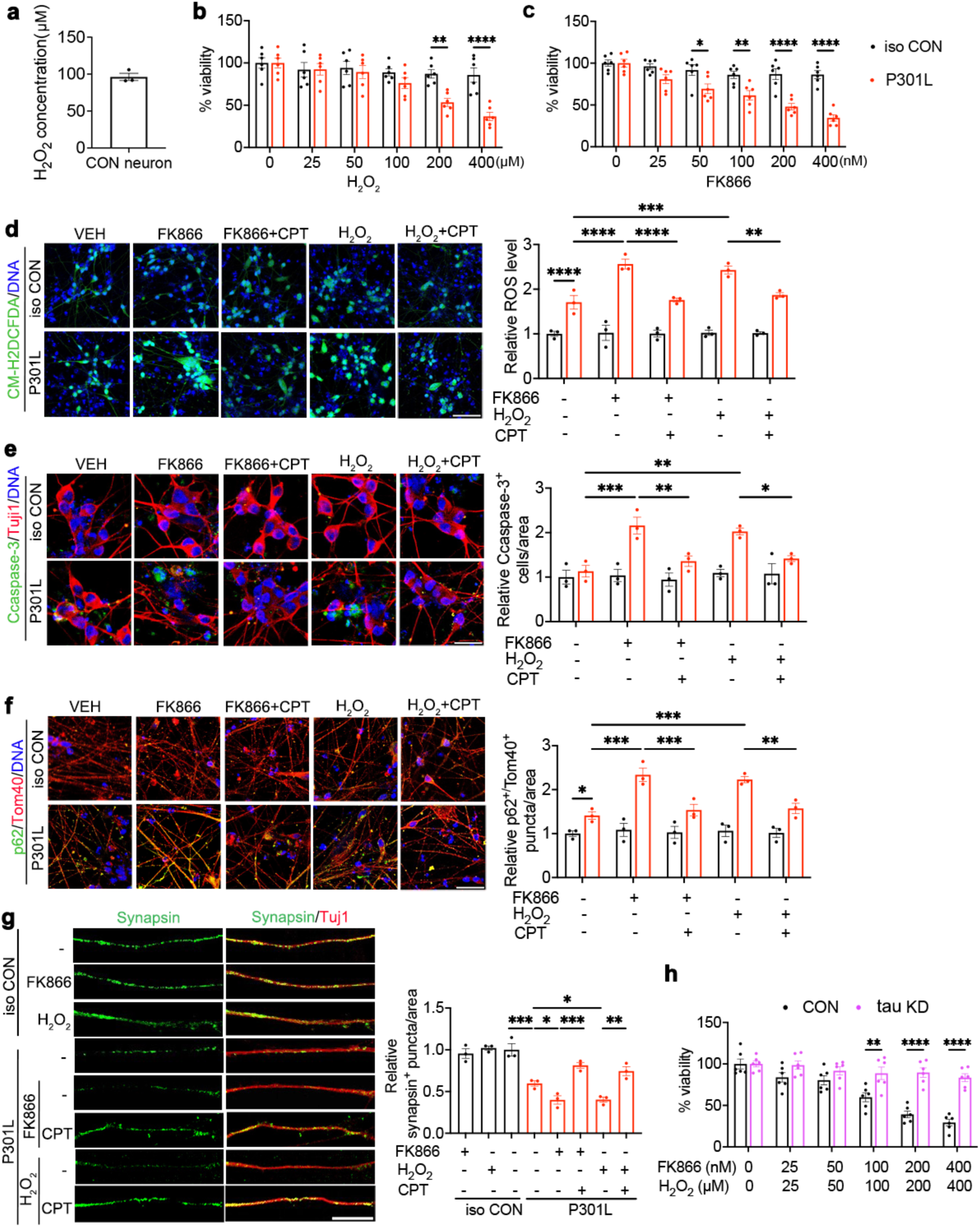
Effect of RET inhibition on neuronal health in tauopathy hiPSC-derived neuronal models of tauopathy. **a**, Measurement of H_2_O_2_ level in normal human iPSC neurons. **b**, **c**, Dose-dependent effect of H_2_O_2_ (**b**) and FK866 (**c**) on the viability of tau-P301L human iPSC neurons and isogenic wildtype controls after 24h treatment. **d**-**f**, Representative images and quantification of ROS (**d**), activated caspase 3 (**e**), and mitochondria-associated p62 aggregates (**f**) in tau-P301L and isogenic WT control iPSC neurons treated with H_2_O_2_ (200 µM) or FK866 (200 nM) individually or co-treated with CPT (2.5 µM). **g**, Representative images and quantification of Synapsin-positive puncta in tau-P301L and isogenic control iPSC neurons treated with H_2_O_2_ (200 µM) or FK866 (200 nM) individually or co-treated with CPT (2.5 µM). **h**, Effect of H_2_O_2_ and FK866 co-treatment on the viability of control and tau KD human iPSC neurons. All data are means ± SEM; statistical significance was determined by two-way ANOVA with Tukey’s post hoc test (**b, c, h**), or one-way ANOVA with Tukey’s post hoc test (**d, e, f, g**). *p<0.05; **p<0.01; ***p<0.001; ****p<0.0001 (**b, c, h**: n=6/group from 3 biological replicates and 2 wells/experiment; **d-g**: n=3 biological replicates, and each data point represents an average of 6 images/experiment).

**Extended Data Fig. 17.**
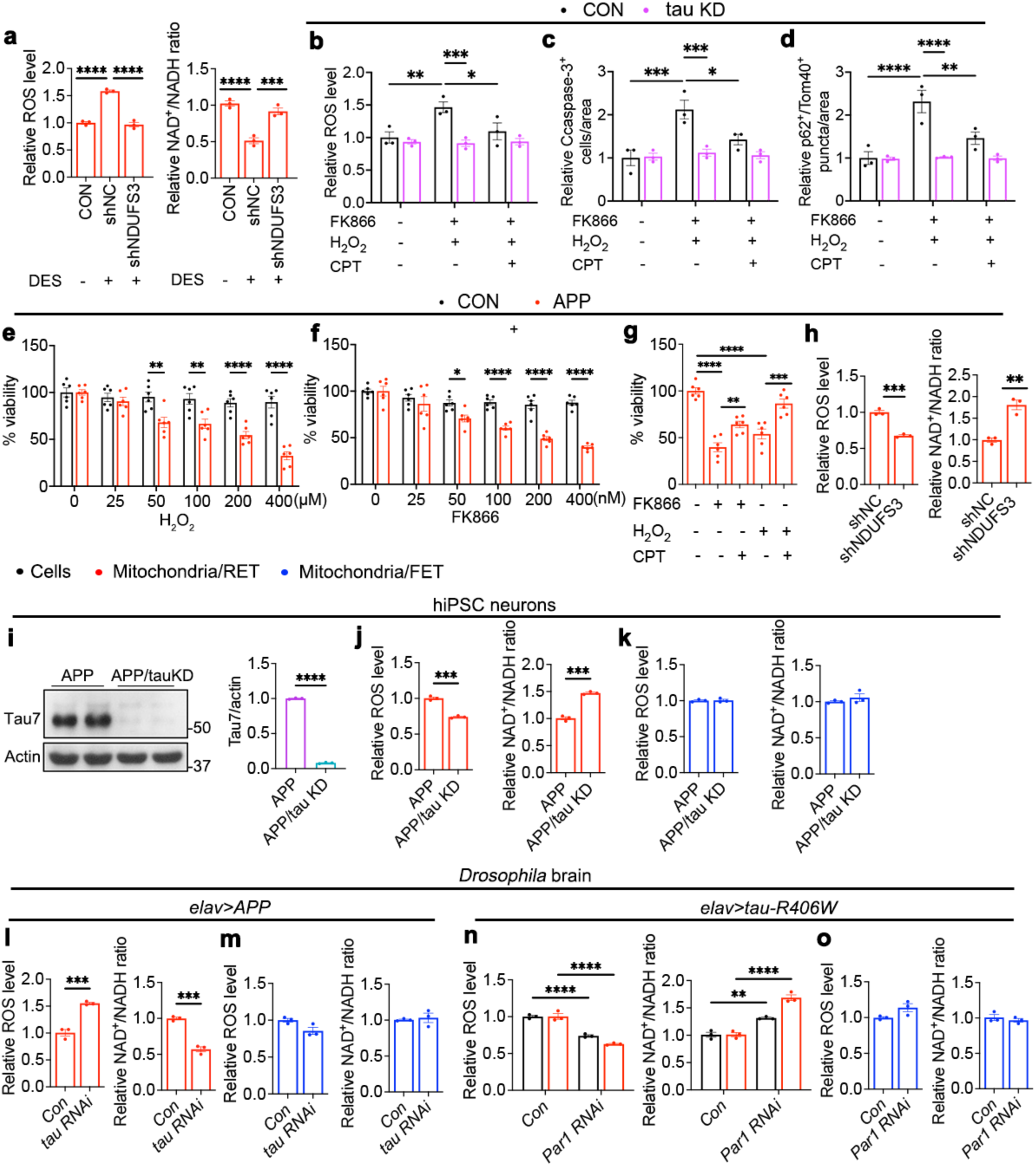
Effects of tau or NDUFS3 knockdown on neuronal health in tauopathy hiPSC neurons and PAR-1 knockdown on tau-related RET activation in *Drosophila*. **a**, Effect of NDUFS3 RNAi on DES-induced RET activation in normal iPSC neurons. **b-d**, Effects of treatment of control and tau KD iPSC neurons with H_2_O_2_ (200 µM) or FK866 (200 nM), individually or co-treated with CPT (2.5 µM), on ROS level (**b**), cleaved caspase 3-positive signal (**c**), and mitochondria-associated p62 positive aggregates (**d**). **e**-**g**, Effects of treatment of control and APP iPSC neurons with H_2_O_2_ (**e**) or FK866 (**f**), individually or co-treated with CPT (**g**), on cell viability. **h**, Effect of NDUFS3 RNAi on RET parameters in APP iPSC neurons. **i**, WB and quantification of tau KD efficiency in APP iPSC neurons. **j**, **k**, Effect of tau KD on RET (**j**) and FET (**k**) in APP iPSC neurons. **l**, **m**, Effect of tau RNAi on RET (**l**) and FET (**m**) in neuronal APP expressing fly brain. **n**, **o**, Effect of PAR-1 RNAi on RET (**n**) and FET (**o**) of brain cells or purified mitochondria of neuronal tau-R406W expressing fly brain. All data are means ± SEM; statistical significance was determined by two-way ANOVA with Tukey’s post hoc test (**b-f**), one-way ANOVA with Tukey’s post hoc test (**a, g**), or two-tailed unpaired Student’s *t* test (**h-o**). *p<0.05; **p<0.01; ***p<0.001; ****p<0.0001 (**b-d**: n=3 biological replicates, and each data point represents an average of 6 images/experiment; **e-g**: n=6/group from 3 biological replicates and 2 wells/experiment; **a, h, i-o**: n=3 biological replicates).

**Extended Data Fig. 18.**
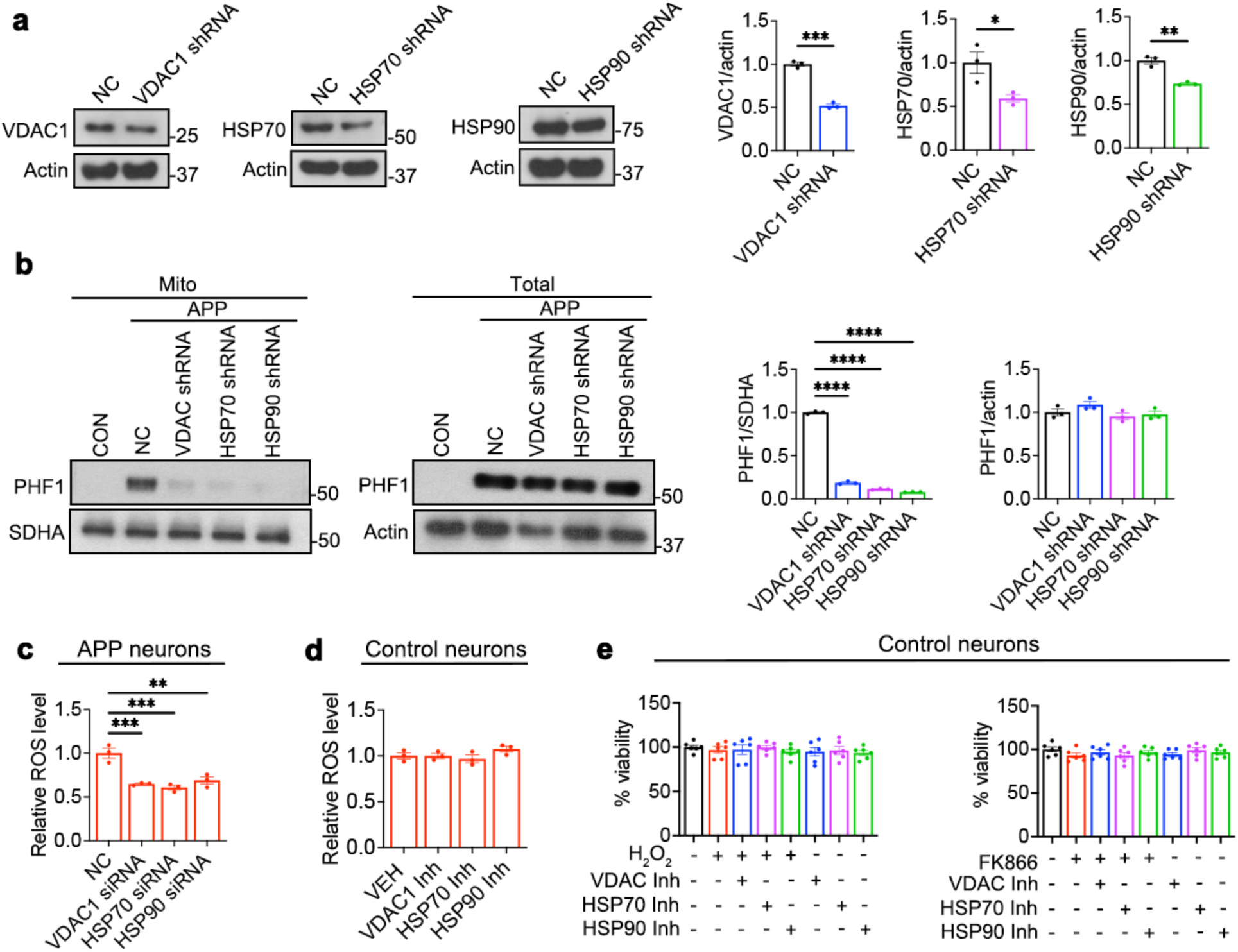
Effects of inhibition of tau entry into mitochondria on RET and stress sensitivity of hiPSC neurons. **a,** Representative WBs and quantification showing effect of VDAC1, Hsp70, and Hsp90 lenti-shRNAs on the expression of the target proteins. **b**, Representative WBs and quantification showing effect of VDAC1, Hsp70, and Hsp90 lenti-shRNAs on the levels of mitochondrially localized PHF-1 tau and total PHF-1 tau in APP hiPSC neurons. Mitochondrial fractions or total cell lysates were used for WB, and mitochondrial or total PHF-1 p-tau was normalized by SDHA or actin. **c**, RET-ROS measurements in mitochondria from control or VDAC1, Hsp70, and Hsp90 lenti-shRNA treated APP hiPSC neurons. **d**, **e**, RET (**d**) and stress sensitivity (**e**) assays of control iPSC neurons treated with VDAC1, Hsp70, and Hsp90 inhibitors. All data are means ± SEM; statistical significance was determined by two-tailed unpaired Student’s *t* test (**a**) or one-way ANOVA with Dunnett’s multiple test (**b-e**). *p<0.05; **p<0.01; ***p<0.001; ****p<0.0001 (**e**: n=6/group from 3 biological replicates and 2 wells/experiment).

**Extended Data Fig. 19.**
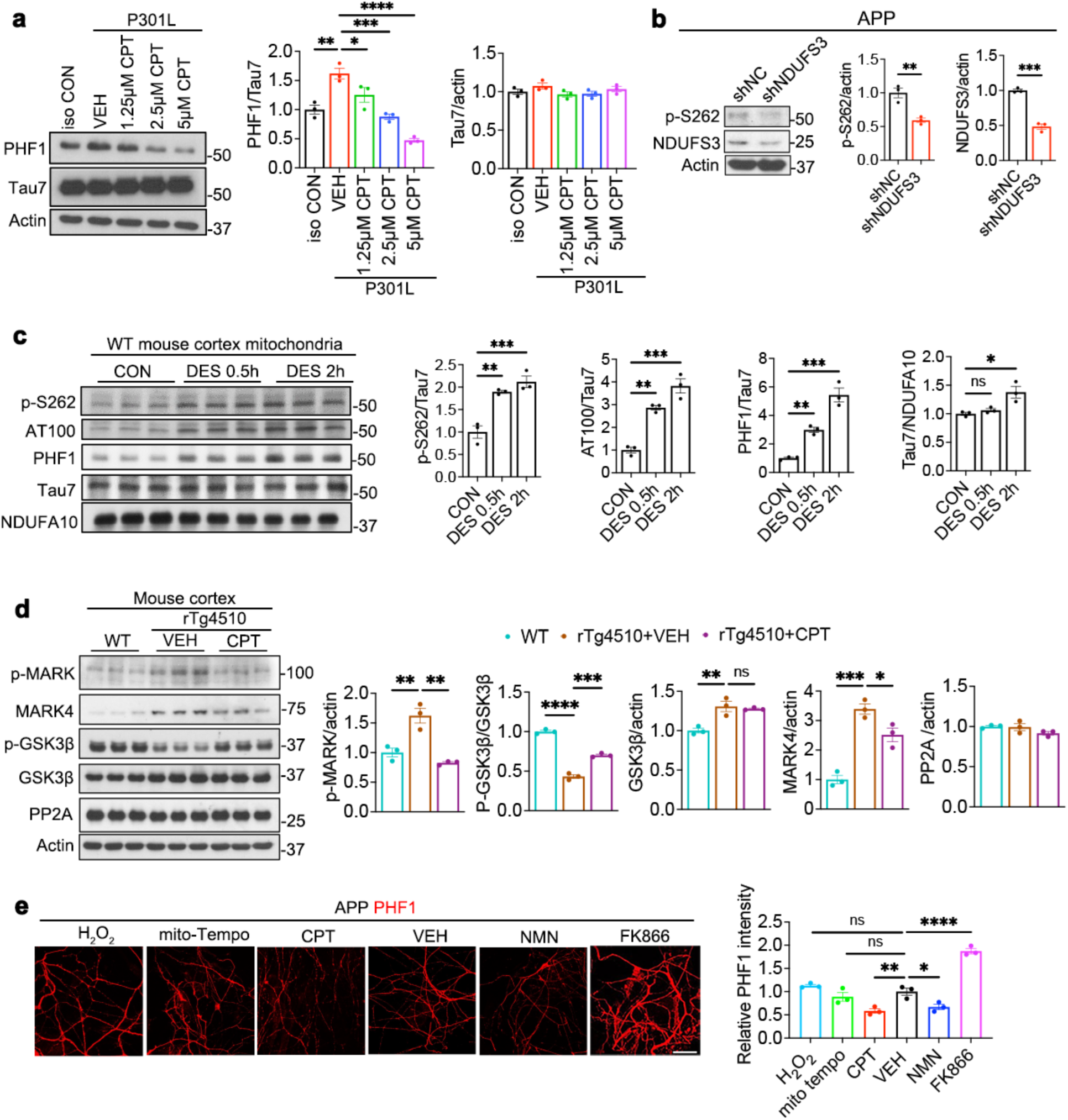
Mechanisms of feedback regulation of tau phosphorylation by RET. **a**, Representative WB and quantification showing dose-dependent effect of CPT treatment on tau phosphorylation in tau-P301L human iPSC neurons. **b**, Representative WB and quantification showing effect of NDUFS3 RNAi on tau phosphorylation in APP iPSC neurons. **c**, WB and quantification of the effect of RET activation by DES treatment on tau phosphorylation in the mitochondrial fraction of wildtype mouse cortex. **d**, WB and quantification of tau kinase and phosphatase levels in the cortex of rTg4510 mice with or without CPT treatment. **e**, Representative images and data quantification of PHF-1 p-tau signal in APP human iPSC neurons under the indicated treatment conditions. All data are means ± SEM; statistical significance was determined by one-way ANOVA with Tukey’s post hoc test (**a, c-e**), or two-tailed unpaired Student’s *t* test (**b**). *p<0.05; **p<0.01; ***p<0.001; ****p<0.0001 (**a, b**: n=3 biological replicates; **c, d**: n=3 mice/group; **e**: n=3 biological replicates with each data point representing an average of 6 images/experiment).

**Extended Data Fig. 20.**
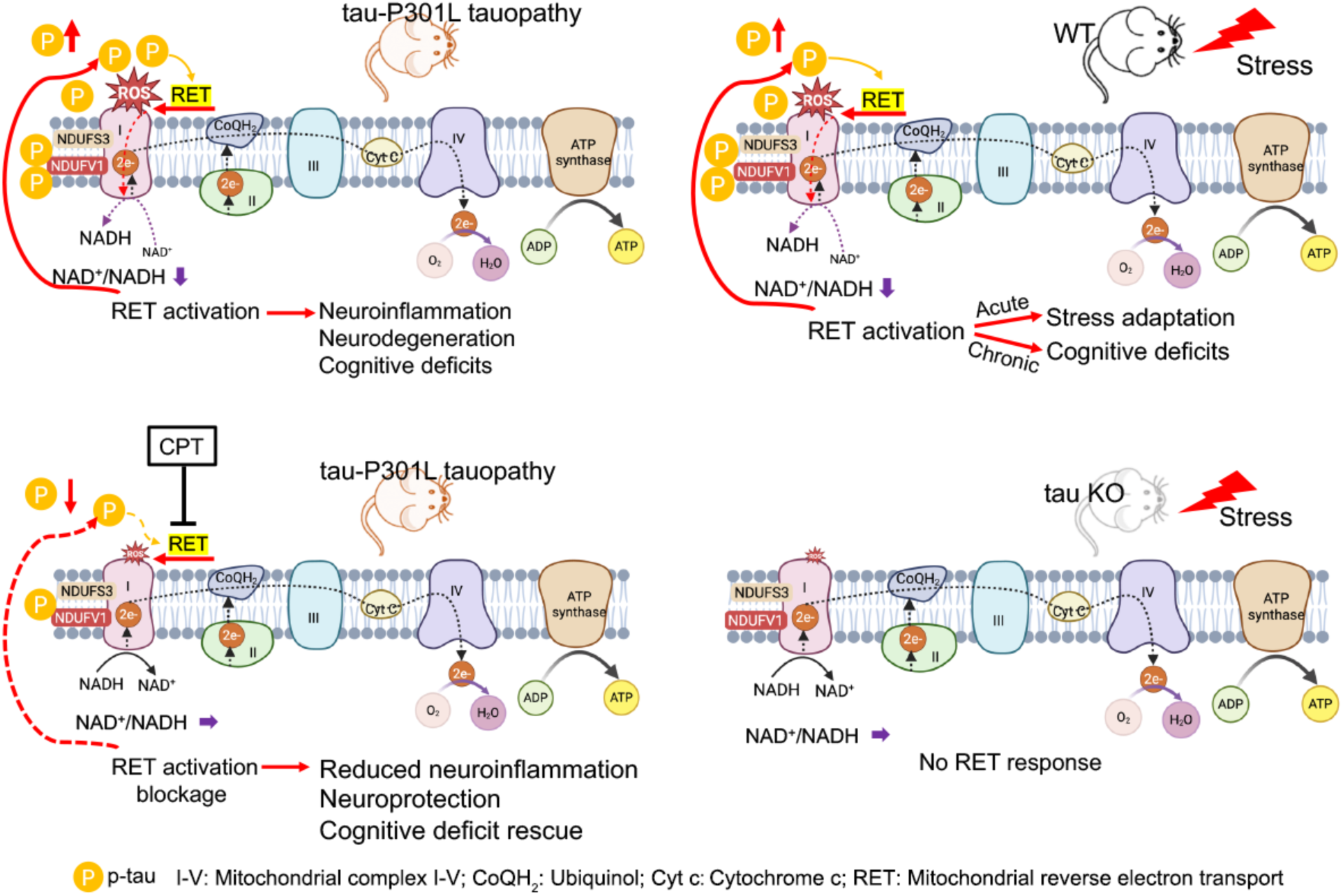
Schematic diagram summarizing the main findings of this study. Stress stimuli like heat stress will activate RET. This may serve some adaptive function initially, but chronic stress will cause aberrant RET activation, leading to tau hyperphosphorylation. p-tau further promotes RET, forming a vicious cycle that causes cognitive deficits. In tau KO animals, this feedback loop is broken, and the animals are spared from stress induced behavioral deficits. In tauopathy mice (e.g., rTg4510), pathogenic tau and elevated p-tau level can promote RET, which causes further tau phosphorylation, forming a vicious cycle that contributes to the tauopathy phenotypes. Inhibition of RET by CPT attenuates this vicious cycle and rescues the disease phenotypes of tauopathy, offering a new therapeutic approach. Black and red dashed lines depict FET along the ETC and RET along C-I, respectively. It remains to be determined whether CoQH_2_ and mitochondrial membrane potential, two previously characterized drivers of RET, are involved in tau-driven RET.

**Extended Data Table S1.**
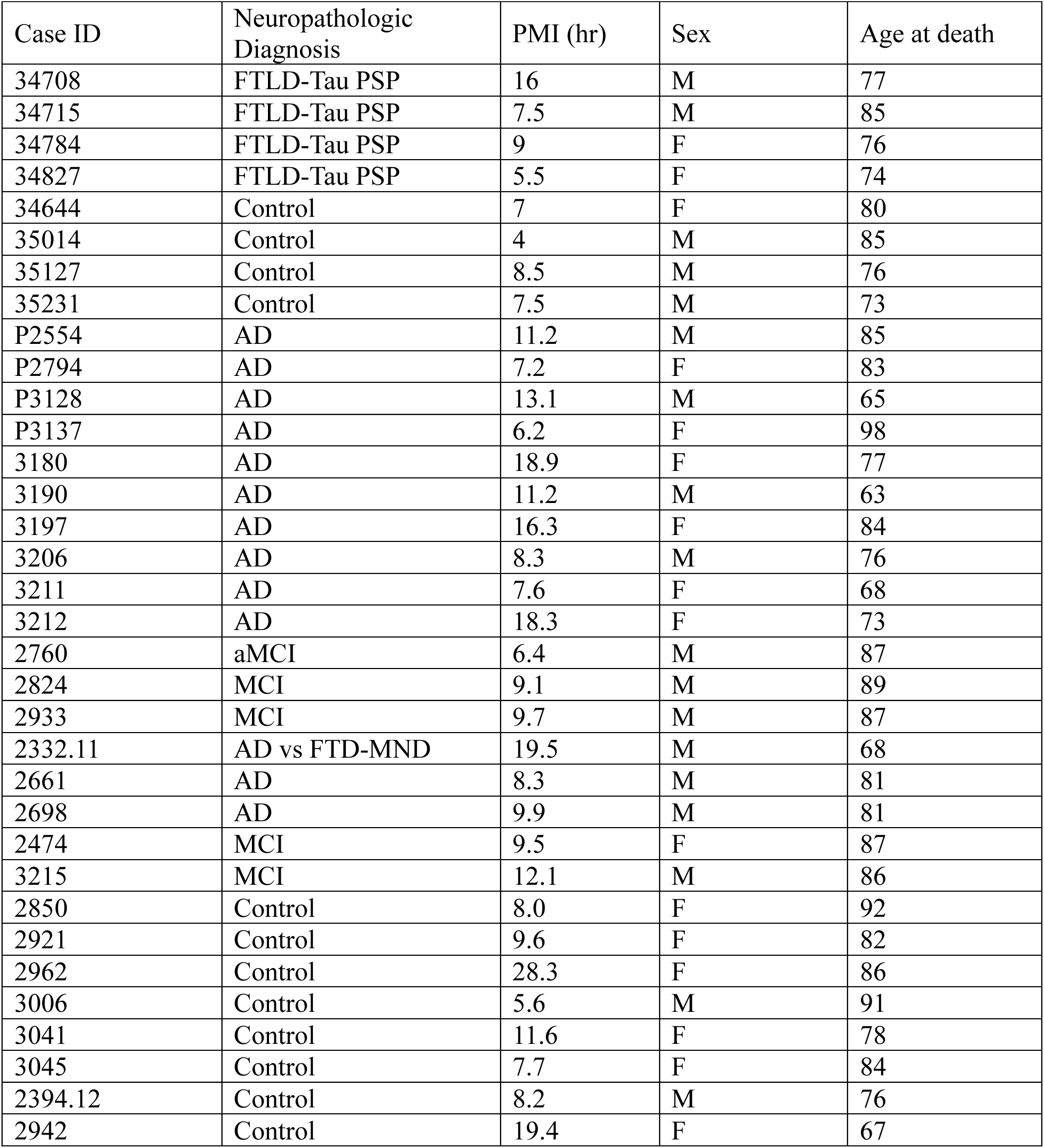
Demographic, pathological, and clinical details of the control and diseased individuals.

